# A stress-dependent TDP-43 SUMOylation program preserves neuronal function

**DOI:** 10.1101/2024.04.12.589206

**Authors:** Terry R. Suk, Caroline E. Part, Jenny L. Zhang, Trina T. Nguyen, Meghan M. Heer, Alejandro Caballero-Gómez, Veronica S. Grybas, Paul M. McKeever, Benjamin Nguyen, Steve M. Callaghan, John M. Woulfe, Janice Robertson, Maxime W.C. Rousseaux

**Author notes:** Authors contributed equally (Jenny L. Zhang, Trina T. Nguyen).

## Abstract

Amyotrophic Lateral Sclerosis (ALS) and Frontotemporal Dementia (FTD) are overwhelmingly linked to TDP-43 dysfunction. Mutations in TDP-43 are rare, indicating that the progressive accumulation of exogenous factors – such as cellular stressors – converge on TDP-43 to play a key role in disease pathogenesis. Post translational modifications such as SUMOylation play essential roles in response to such exogenous stressors. We therefore set out to understand how SUMOylation may regulate TDP-43 in health and disease. We find that TDP-43 is regulated dynamically via SUMOylation in response to cellular stressors. When this process is blocked *in vivo*, we note age-dependent TDP-43 pathology and sex-specific behavioral deficits linking TDP-43 SUMOylation with aging and disease. We further find that SUMOylation is correlated with human aging and disease states. Collectively, this work presents TDP-43 SUMOylation as an early physiological response to cellular stress, disruption of which may confer a risk for TDP-43 proteinopathy.

## Introduction

Altered proteostasis is one of the key hallmarks of aging and is particularly prevalent in neurodegenerative diseases[1]. Despite the highly heterogenous nature of neurodegenerative diseases – likely owing to the extreme genetic and environmental diversity acting upon individuals – a select few proteins are recurringly implicated in pathology[2]. TDP-43 is an essential RNA binding protein notorious for its involvement in neurodegenerative diseases including Amyotrophic Lateral Sclerosis (ALS) and Frontotemporal Dementia (FTD). In a diseased state, TDP-43 is found mislocalized from the nucleus and aggregated in the cytoplasm of neurons within the central nervous system (CNS) in ∼97% and ∼45% of ALS and FTD cases, respectively[3, 4]. Additionally, ALS and FTD exist on a clinical and genetic spectrum linked by TDP-43 dysfunction often referred together as ALS/FTD[5]. Beyond ALS/FTD, TDP-43 pathology is increasingly linked to other neurological disorders including Limbic-predominant Age-related TDP-43 Encephalopathy (LATE), Alzheimer’s Disease (AD), Chronic Traumatic Encephalopathy (CTE), and Stroke[6–9]. Thus, TDP-43 pathology has garnered much attention to better understand the causes of these diseases and to uncover potential routes of therapeutic intervention.

The accumulation of cytoplasmic TDP-43 aggregates is considered a late-stage event in neurodegeneration. Increasing evidence suggests that the partial mislocalization or complete depletion of TDP-43 from the nucleus to the cytoplasm is an early event in ALS/FTD pathogenesis, functioning independently – however tightly associated – with aggregation[10–16]. As a result, recent efforts have focused on the outcomes of TDP-43 mislocalization as early markers of dysfunction and/or mechanisms driving disease. Indeed, loss of function due to mislocalization of TDP-43 causes aberrant cryptic splicing in genes including *STMN2* and *UNC13A* that drive the progression of ALS[12–15, 17]. To wit, cryptic mis-splicing of *STMN2* was recently found to correlate strongly with TDP-43 pathology[18, 19]. Additionally, once in the cytoplasm, TDP-43 can exert additional toxicity through gain of function effects by sequestering critical proteins into cytoplasmic aggregates thus disrupting crucial pathways leading to cellular demise[11, 20]. Together, dysregulation of TDP-43 is sufficient to drive neuronal dysfunction ultimately leading to neurodegeneration. However, the mechanisms instigating TDP-43 pathogenesis remain convoluted.

Despite the overwhelming prevalence of TDP-43 pathology in ALS/FTD and related diseases, mutations in the gene encoding TDP-43 (*TARDBP*) are only present in less than 1% of all ALS and FTD cases[21]. It is becoming evident that not one, but many genetic and/or environmental factors affect pathways converging on TDP-43 in ALS/FTD. Many ALS/FTD causative genes exert toxicity by disruption of key pathways such as nucleocytoplasmic transport and cellular stress responses[22]. The cellular stress response is a critical pathway tightly linked to ALS/FTD and TDP-43 supported via genetic, experimental, and epidemiological evidence [23–36]. On the one hand, ALS/FTD-linked mutations disrupting various steps of the stress response pathways converge on the dysregulation of TDP-43 resulting in pathology[23–25, 30–33, 36]. On the other hand, exogenous insults such as aging, head injuries, viral infections, and other exposures may confer susceptibility or precipitate ALS/FTD and can serve as pre-clinical models of TDP-43 proteinopathy[8, 37–39]. Motor neurons, the primary vulnerable cell type in ALS, are thought to be particularly susceptible to stress due to the high levels of excitotoxicity experienced throughout one’s lifetime[40]. Indeed, prolonged cellular stress due to chronic stress exposure or failures in stress recovery can result in TDP-43 pathology[28, 41]. However, much less is known about the molecular pathways acting upon TDP-43 in the cell stress response and how they might be linked to age-related neurodegeneration. Uncovering these mechanisms will help to better understand how environmental insults that occur throughout aging converge on TDP-43 and cause susceptibility to disease.

Post translational modifications (PTMs) play key roles in regulating protein function and have been tightly linked to TDP-43 (dys)function and disease. TDP-43 is modified by an array of PTMs including phosphorylation, ubiquitination, acetylation, and polyADP-ribosylation[42–46]. Abnormal TDP-43 phosphorylation and ubiquitination are pathognomonic of ALS and TDP-43 related FTD[42]. Recent studies have suggested that SUMOylation – the covalent conjugation of a Small Ubiquitin-like Modifier (SUMO) to target lysine residues[47, 48] – by SUMO1 may have a role in regulating TDP-43 nucleocytoplasmic transport and RNA binding[49–52]. SUMO1 is best characterized for its roles in nucleocytoplasmic shuttling[53]. SUMO2 however is the only essential SUMO paralog and selectively plays roles in orchestrating cellular stress responses[54, 55]. SUMO2 has previously been observed in TDP-43 aggregates *in vitro* and is related with TDP-43 insolubility[56, 57]. However, it is unclear whether TDP-43 is a direct target of SUMO2 and what the implications are on TDP-43 function and disease pathogenesis.

Here, we show that TDP-43 is modified by SUMO2 selectively in response to cellular stressors. TDP-43 becomes SUMOylated within the nucleus early in response to stress, upstream of TDP-43 aggregation. Modification by SUMO2 is further correlated with dosage and duration of cellular stress and peaks during the recovery phase before it is cleared through the ubiquitin proteosome system. We further identified four E3 SUMO ligases that can modulate the levels of TDP-43 SUMOylation. We found that TDP-43 is SUMOylated by SUMO2 in a conserved region of the C-terminal domain at lysine (K) 408 directly adjacent to phosphorylation residues serine (S) 409/410 characteristically phosphorylated in TDP-43 aggregates. To understand the physiological consequences of blocking TDP-43 SUMOylation, we generated a knock in mouse model bearing a p.K408R point mutation in endogenous mouse *Tardbp* allele. Cortical neurons cultured from these mice display impaired stress recovery and accumulation of nuclear TDP-43. These mice do not show abnormalities in development but develop mild social and cognitive deficits as they age. Pathologically, we observe TDP-43 mislocalization and accumulation of phosphorylated TDP-43 in the spinal cord and significant denervation of neuromuscular junctions in aged mice. As SUMOylation of TDP-43 plays a protective role in mice during aging, we assessed human brain samples and observed a positive correlation between global SUMOylation and age inferring an increased demand on SUMO-related pathways during aging. Finally, we observed significant increase in TDP-43 and SUMO2 interactions in the prefrontal cortex from individuals diagnosed with ALS/FTD compared to unaffected controls suggesting SUMOylation is actively engaged in regulating TDP-43 in disease states.

## Results

### TDP-43 is SUMOylated in the nucleus in a context specific manner

Various acute stressors including oxidative (sodium arsenite, NaAsO_2_), hyperosmotic (D-sorbitol and sodium chloride), and heat shock have been demonstrated to induce TDP-43 mislocalization and aggregation *in vitro*, helping to uncover key mechanisms affected in disease[24, 58, 59]. As sodium arsenite is a commonly used stressor to interrogate TDP-43 alterations, we examined whether TDP-43 became SUMOylated in response to sodium arsenite treatment. Using an immunoprecipitation assay (“SUMOylation Assay”, Fig. 1A) to immunoprecipitate HA-SUMO bound proteins under denaturing conditions to disrupt non-covalent protein interactions, we found that TDP-43 becomes SUMOylated by SUMO2 and can form polySUMO chains specifically in response to stress (Fig. 1B). Interestingly we found that sodium arsenite and heat shock stressors, but not hyperosmotic stress, induced TDP-43 SUMOylation suggesting that SUMOylation is a context specific modifier of TDP-43 (Fig. S1A).

**Fig. 1:**
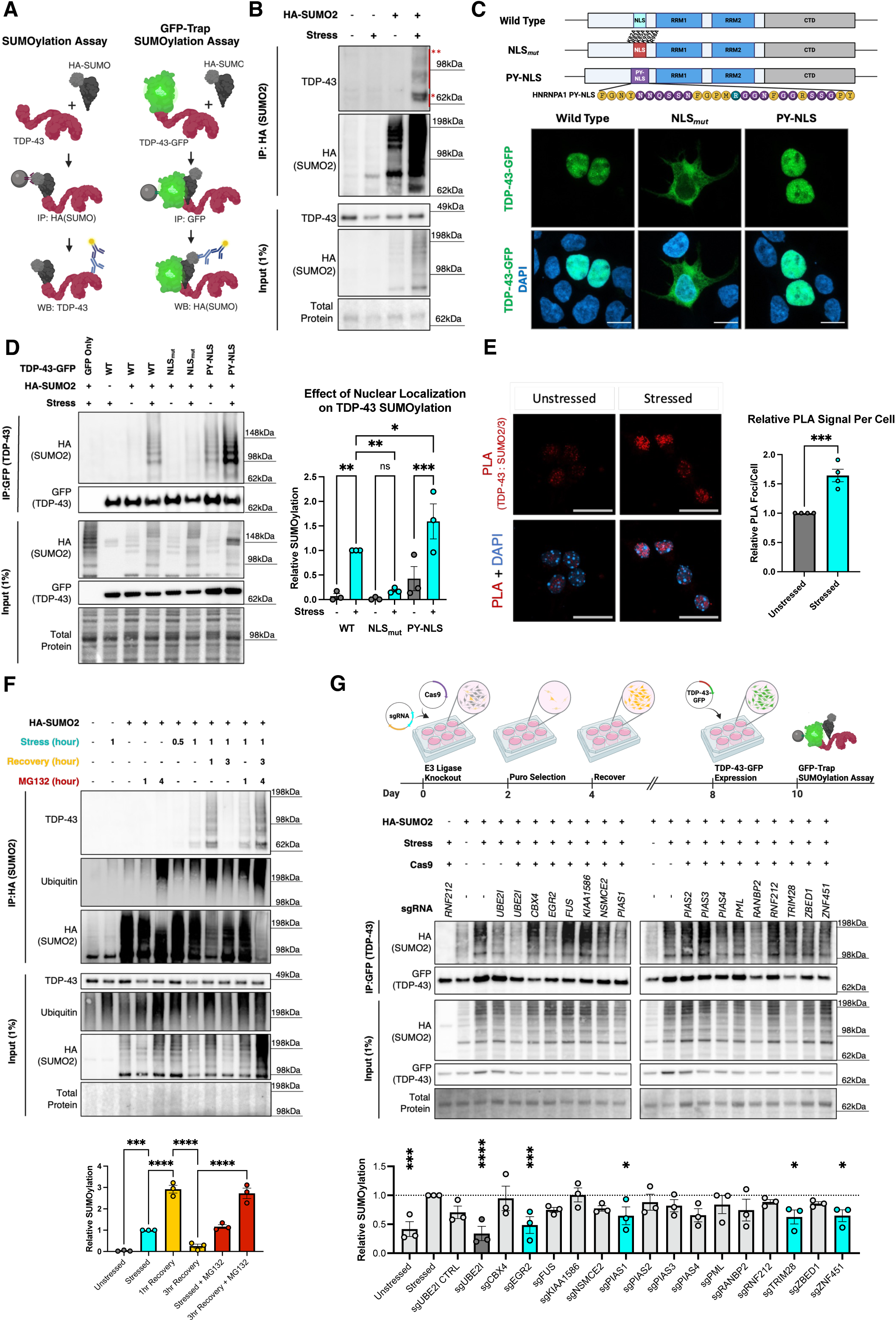
SUMOylation dynamically regulates nuclear TDP-43 in a stress-responsive manner. **(A)** Schematic of immunoprecipitation assays to detect TDP-43 SUMOylation. **(B)** Representative SUMOylation Assay western blot detecting TDP-43 SUMOylation specifically in response to 1 hour sodium arsenite (250 µM) stress. * = SUMOylated TDP-43, ** = PolySUMOylated TDP-43. **(C)** Schematic TDP-43-GFP Nuclear Localization Sequence (NLS) variants and representative florescent microscopy analysis of their subcellular localization. (Scalebar = 10 µm). Yellow amino acids represent those critical for PY-NLS function. Teal arginine residue was mutated from the native HNRNPA1. **(D)** Representative GFP-trap SUMOylation assay detecting loss of SUMOylation in response to TDP-43 mislocalization in response to 1 hour sodium arsenite stress (250 µM). (N=3) RM One-Way ANOVA with Fisher’s LSD test data presented as mean ± SEM, * p<0.05, ** p<0.005, *** p<0.0005. **(E)** Representative image and quantification of relative proximity ligation signal between TDP-43 and SUMO2/3 in murine primary cortical neuron cultures in response to 1 hour sodium arsenite (250 µM) treatment. Scalebar = 20 μm. (N = 4 per condition) Unpaired T-test data presented as mean ± SEM, *** p<0.0005 **(F)** Representative SUMOylation Assay of TDP-43 SUMOylation dynamics during stress (250 µM sodium arsenite) and recovery demonstrating a dependency on the ubiquitin proteosome for clearance of polySUMOylated TDP-43 by treatment with proteasome inhibitor MG132 (2µM). **(G)** Schematic and representative GFP-Trap SUMOylation assay screening for potential E3 ligases regulating TDP-43 SUMOylation in response to 1 hour sodium arsenite stress (250 µM). (N = 3) RM One-Way ANOVA with Fisher’s LSD test data presented as mean ± SEM, * p<0.05, ** p<0.005, *** p<0.0005, **** p<0.0001.

Previous reports have suggested that TDP-43 is SUMOylated under native conditions by SUMO1 to regulate nucleocytoplasmic transport and RNA binding[49]. Strikingly, we found that TDP-43 SUMOylation was selectively modified by SUMO2 and SUMO3, but not SUMO1, specifically under stressed conditions; consistent with the conserved roles of SUMO2 and SUMO3 in response to stress (Fig. S1B). Mature (covalently bound) SUMO2 and SUMO3 share nearly 100% amino acid homology making them difficult to differentiate and thus referred together as SUMO2/3. Since SUMO2 is the only essential and most abundantly expressed SUMO paralog in the CNS and sodium arsenite is commonly used to study TDP-43 pathobiology, we focused on TDP-43 SUMOylation by SUMO2 in response to sodium arsenite stress[54, 60].

To test whether TDP-43 SUMOylation is linked with disease-like states, we expressed ALS-linked mutant TDP-43 (TDP-43^Q331K^) and found that TDP-43 SUMOylation was significantly increased compared to wild type TDP-43 (Fig. S1C). Next, we disrupted the nuclear localization sequence in TDP-43 (TDP-43^NLSmut^) to mislocalize TDP-43 to the cytoplasm and found that in response to sodium arsenite, TDP-43^NLSmut^ blocks SUMOylation indicating that nuclear localization is essential for TDP-43 SUMOylation (Fig. 1C,D). Classically, lysine (K) residues, in addition to arginine (R) residues, are critical components of a traditional NLS to enable the interaction with importins to facilitate nuclear import of proteins; but they may also be direct targets of SUMOylation[61]. To determine whether loss of TDP-43 SUMOylation is mediated through loss of nuclear localization or disruption of lysine (K) residues in the NLS, we sought to restore TDP-43 localization to rescue TDP-43 SUMOylation. We surveyed the literature to identify a NLS that functions independent of lysine (K) residues. Previous studies have demonstrated that the RNA binding protein HNRNPA1 contains a non-conventional PY-NLS to facilitate nuclear localization [62, 63]. We replaced the native TDP-43 NLS with an HNRNPA1 PY-NLS (TDP-43^PY-NLS^) and found that TDP-43^PY-NLS^ localizes to the nucleus reflecting wild type TDP-43 (Fig. 1C). By performing a GFP-Trap SUMOylation assay we found that TDP-43^PY-NLS^ rescues the loss of SUMOylation observed when expressing TDP-43^NLSmut^ indicating that nuclear localization, and not the lysine (K) residues within the NLS, is required for stress-dependent TDP-43 SUMOylation (Fig. 1D).

To further validate our findings, and to visualize the interaction between TDP-43 and SUMO2 in primary cortical neurons. As TDP-43 and SUMO2 are widely expressed throughout the nucleus, co-localization analysis is challenging to infer SUMOylation events (Fig. S1D). To visualize the interaction between TDP-43 and SUMO2/3 we performed a proximity ligation assay (PLA) in murine primary cortical neurons cultures (Fig. 1E, Fig. S1E). We found that in response to sodium arsenite there is a significant increase in PLA signal between endogenous TDP-43 and SUMO2/3, consistent with the stress-dependent SUMOylation previously observed. Taken together, SUMOylation may be an important mechanism regulating nuclear TDP-43 upstream of mislocalization in response to stress.

### SUMOylation is an early event in response to stress and helps clear TDP-43 through the ubiquitin proteasome system during recovery

Prolonged cellular stress *in vitro* leads to mislocalization of TDP-43 and progressive transition into insoluble aggregates resembling aspects of pathology observed in disease[16, 28, 31, 33, 64]. As TDP-43 SUMOylation occurs within the nucleus upstream of nuclear egress, we postulated that SUMOylation would occur upstream of aggregation during prolonged sodium arsenite stress. To test this hypothesis, we performed a time course assay to characterize the dynamics of TDP-43 SUMOylation during prolonged treatment with sodium arsenite. We observed that TDP-43 becomes readily SUMOylated in the first 15-30 minutes of the acute stress response upstream of the accumulation of RIPA-insoluble, phosphorylated TDP-43 (Fig. S1F).

Furthermore, we observed an increase in TDP-43 SUMOylation that linearly correlated with the duration of stress (R^2^ = 0.6922, p<0.0001). To determine whether SUMOylation occurs in response to relative stress intensity, we performed a dose response assay and found that TDP-43 SUMOylation is proportional to relative stress intensity (Fig. S1G). Therefore, the increase in TDP-43 SUMOylation during prolonged stress is likely due to an increased stress burden over time.

To determine the fate of SUMOylated TDP-43, we performed a stress-recovery time-course assay where cells were stressed with sodium arsenite for up to 1 hour, then allowed cells to recover after washout. For the first hour during stress recovery, we observed an ∼3-fold increase in TDP-43 SUMOylation indicating that TDP-43 continues to be SUMOylated and polySUMOylated during the early stages of stress recovery (Fig. 1F). Furthermore, SUMOylated TDP-43 was not detectable after 3 hours of recovery, suggesting that most of the clearance occurs between 2-3 hours post stress. TDP-43 is known to interact with the SUMO-Targeted Ubiquitin Ligase RNF4 which can polyUbiquitinate SUMOylated proteins to be degraded through the UPS[65]. To determine whether the UPS functions to clear SUMOylated TDP-43 during stress recovery, we treated cells with the proteasome inhibitor MG132 and monitored TDP-43 SUMOylation during stress and recovery. We observed that inhibiting the UPS system by MG132 treatment prevented clearance of SUMOylated TDP-43 during post-stress recovery indicating that SUMOylated TDP-43 is marked for degradation by the UPS pathway.

Importantly, treatment with MG132 for 1 or 4 hours was not sufficient to induce TDP-43 SUMOylation by itself, consistent with the stress-selective nature inducing TDP-43 SUMOylation. Taken together, TDP-43 SUMOylation occurs early in the stress response which then leads to ubiquitin-targeted clearance during stress recovery demonstrating SUMOylation may be an important mechanism regulating nuclear TDP-43 proteostasis.

### Stress-dependent TDP-43 SUMOylation is mediated by select E3 SUMO ligases

E3 SUMO ligases are important mediators for SUMOylation of select substrates and enable spatiotemporal regulation of this modification. Thus, they may represent critical modulators of TDP-43 SUMOylation. To identify potential E3 SUMO ligases that may regulate TDP-43 SUMOylation, we first performed a literature search to identify potential ligases with evidence of mediating SUMOylation. We prioritized 15 candidate E3 SUMO ligases and using a dual sgRNA/Cas9 approach generated knockout cell lines for each of the candidate ligases and performed GFP-Trap SUMOylation assays to test the effects on TDP-43 SUMOylation (Fig. 1G)[66–80]. As a positive control, we knocked out the sole E2 SUMO ligase, UBC9 (encoded by *UBE2I*), which led to a near complete loss of TDP-43 SUMOylation demonstrating the functionality of the approach. We identified four SUMO E3 ligases whose knockout consistently reduced the levels of stress-induced TDP-43 SUMOylation: *EGR2 (KROX20), PIAS1, TRIM28 (KAP1),* and *ZNF451 (ZATT).* EGR2 is linked to the neuromuscular disorder Charcot Marie Tooth Disease and is an immediate early gene rapidly reacting to external cellular stimuli aligning with the early response of TDP-43 SUMOylation upon stress[81, 82]. PIAS1 is a canonical, highly conserved E3 SUMO ligase known to play roles in cellular stress responses[83]. TRIM28 is also a well characterized E3 SUMO ligase which is a predicted interactor of SUMOylated TDP-43 based on the GPS-SUMO2.0 algorithm[84–86]. ZNF451 is characterized as a stress dependent E3 SUMO ligase specifically promoting polySUMOylation with SUMO2/3. It is sometimes referred to as an “E4 SUMO elongase” which complements our data that TDP-43 is polySUMOylated in response to stress[80, 87]. Identification of these putative TDP-43 SUMO ligases supports the robustness of stress-induced TDP-43 SUMOylation and adds layers of nuance into the regulation of TDP-43 with variable routes of modulating TDP-43 functions.

### TDP-43 SUMOylation occurs in a conserved region of the C-terminal domain

SUMOylation occurs at lysine (K) residues often residing within a consensus SUMOylation motif, *Ψ*-K-x-D/E (*Ψ*=large hydrophobic residue, x = any amino acid), however it is increasingly recognized that SUMOylation at lysine residues in non-consensus motifs is not uncommon and plays significant roles in regulating protein function[85]. Using GPS-SUMO (https://sumo.biocuckoo.cn/) to predict likely SUMOylation sites on TDP-43, we identified a consensus SUMOylation motif within the first RNA recognition motif at K136 – thought to be the site targeted by SUMO1 – and a non-consensus SUMOylation motif within the C-terminal domain at K408[85, 86].

To map the residue of stress induced TDP-43 SUMOylation, we expressed TDP-43 mutating the candidate lysine (K) residues to arginine (R) residues to maintain similar charge and structure of the native amino acid sequence while blocking SUMOylation: K136R, K408R, and K136R/K408R “2KR”. By performing a GFP-trap SUMOylation assay, we observed that TDP-43^K136R^ did not block sodium arsenite-induced TDP-43 SUMOylation (Fig. 2A). Through fluorescent microscopy, we observed that TDP-43^K136R^ formed nuclear puncta with resemblance to RNA-binding deficient TDP-43 (5FL) (Fig. S2A). We predicted the structure of TDP-43 and TDP-43^K136R^ with AlphaFold3 in the presence of UG repeated RNA and found that K136R is predicted to slightly alter the tertiary structure of TDP-43 and affect the interaction with RNA (Fig. S2B and S2C). Thus, phenotypes observed in TDP-43^K136R^, as reported by others, may be facilitated by loss of RNA binding functions independent of loss of SUMOylation[50–52]. In contrast, through GFP-trap SUMOylation assays we found that TDP-43^K408R^ mutant significantly reduced TDP-43 SUMOylation by ∼50%, suggesting K408 is a major site of SUMOylation in response to sodium arsenite stress (Fig. 2A). Importantly, mining two independent unbiased mass spectrometry studies identifying sites of SUMOylation across the proteome in stressed and unstressed conditions also uncovered TDP-43 SUMOylation at K408, supporting our findings [88, 89]. Through florescent microscopy analysis, we found that expression of TDP-43^K408R^ did not induce nuclear TDP-43 puncta, but instead led to a slight albeit insignificant increase in cells presenting with mislocalized TDP-43 (Fig. S3D). However, overexpression of TDP-43^K408R^ did not uniformly induce mislocalization in all cells, consistent with the dynamic nature of TDP-43 SUMOylation and the requirement of stress to introduce TDP-43 SUMOylation responses. Finally, TDP-43^2KR^ significantly reduced TDP-43 SUMOylation to the same extent as TDP43^K408R^ suggesting that K408 and not K136 is a target of stress-induced SUMOylation (Fig. 2A). We further observed that TDP-43^2KR^ expression is less stable than TDP-43^K408R^ likely due to the structural changes mediated by K136R. Taken together, K408 is a major target of stress-induced SUMOylation.

**Fig. 2:**
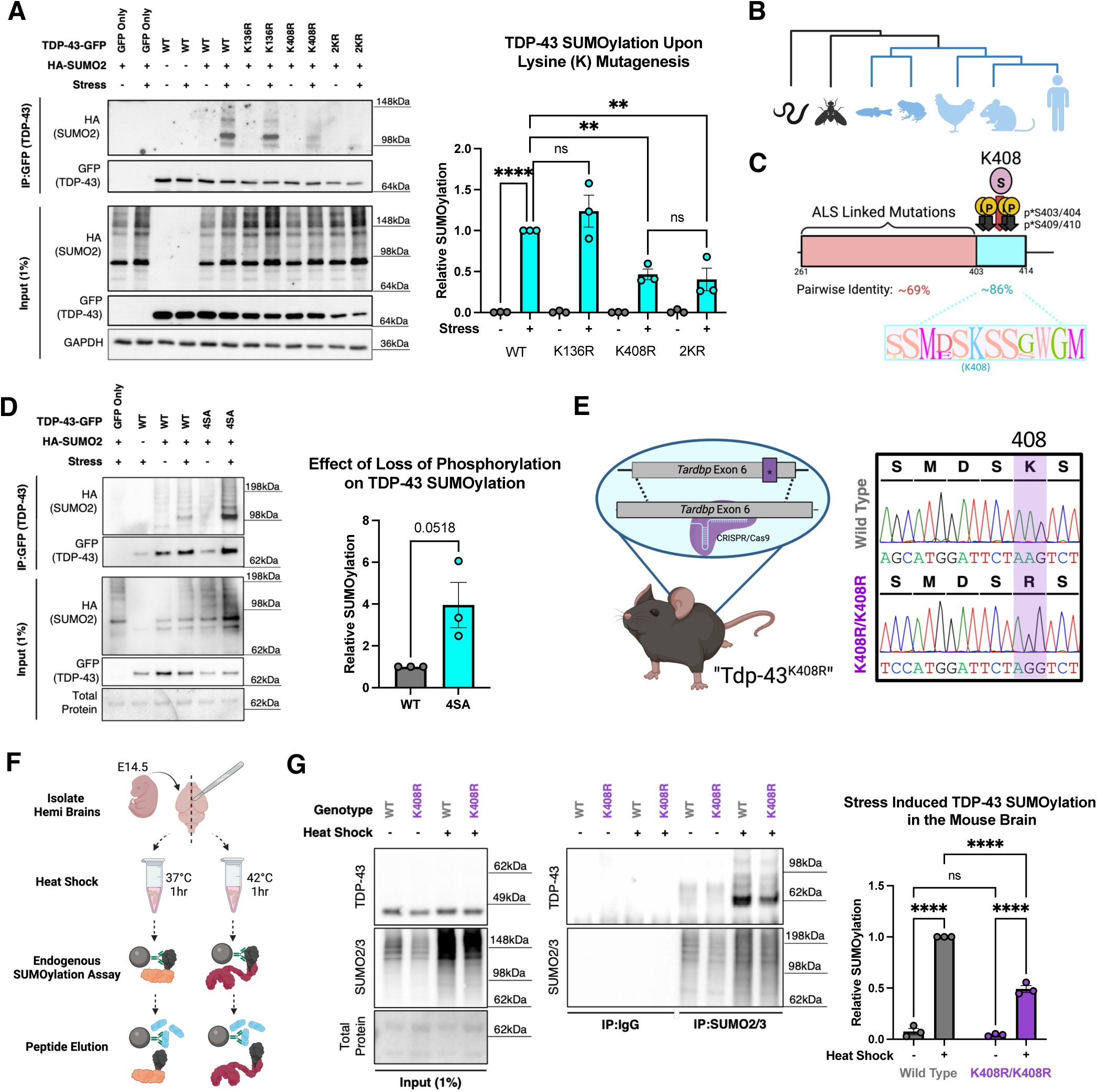
TDP-43 is SUMOylated at K408 in a conserved region of the C-terminal domain. **(A)** Representative GFP-Trap SUMOylation assay to map the site of TDP-43 SUMOylation in response to 1 hour sodium arsenite (250 μM) stress. One-Way ANOVA with Fisher’s LSD test data presented as mean ± SEM, ** p<0.005, **** p<0.0001. **(B)** Phylogenetic representation from alignment of representative TDP-43 paralogs highlighting the emergence and conservation of [human] K408 (Blue). **(C)** Pairwise identity of the [human] TDP-43 C-terminal domain from MUSCLE alignment of 300 amino acid sequences for TDP-43 paralogs from Humans to Actinopterygii from OrthoDB. Red represents the TDP-43 C-terminal domain (Intrinsically disordered domain). Cyan represents PTM-enriched region at the extreme C-terminus. **(D)** Representative GFP-Trap SUMOylation assay with phospho-dead TDP-43 “4SA” (S403/404/409/410A) highlighting antagonism between SUMOylation and phosphorylation. (N=3) Unpaired T-test data presented as mean ± SEM. **(E)** Schematic and sanger sequencing of the TDP-43^K408R^ knock in mouse line. **(F)** Schematic of endogenous SUMOylation assay from embryonic mouse brains. **(G)** Validation of loss of TDP-43 SUMOylation in the brains of TDP-43^K408R^ mice. 2-Way ANOVA with Fisher’s LSD test. Data presented as mean ± SEM, **** p<0.0001.

To address the evolutionary conservation of TDP-43 SUMOylation at K408, we aligned the amino acid sequences of TDP-43 with 300 orthologs from OrthoDB and found that human K408 is nearly completely conserved throughout jawed-vertebrates concurrent with the evolution of the C-terminal domain (Fig. 2B, Supplementary Table 1). While this domain harbors the majority of ALS/FTD causing mutations, the residues surrounding K408 display considerably high levels of conservation (∼86% pairwise identity) indicating that this motif may play important roles in regulating TDP-43 in vertebrates that may be altered by disease causing mutations (Fig. 2C, Supplementary Table 1). Consistent with these findings, no missense variants have been identified that disrupt the SUMOylation motif (based on gnomAD v4.0[90]) supporting its robust conservation in the healthy human population.

The residues surrounding TDP-43 K408, specifically S403, S404, S409, and S410 are characteristically phosphorylated in patients with TDP-43 proteinopathy[42]. To determine whether phosphorylation of TDP-43 at these serine residues interact with SUMOylation at K408, we expressed phospho-dead “4SA” TDP-43 (S403/404/409/410A) and observed that TDP-43^4SA^ does not lead to loss of SUMOylation following stress, but rather appeared to promote an increase in its SUMOylation (Fig. 2D). Whether phosphorylation functions allosterically to remove SUMO2/3 at K408 or antagonistically to block the SUMOylation recognition site remains unclear. The independent modification by either phosphorylation or SUMOylation may serve as a mechanism to differentially regulate TDP-43 function independently or sequentially in response to stimuli. Thus, while TDP-43 phosphorylation has gained significant attention since its initial discovery[42] – in part due to the availability of phospho-specific antibodies – we now show that the C-terminal domain contains a conserved “PTM-enriched region” where SUMOylation at K408 may precede phosphorylation, revealing additional complexity and nuance to the stimulus-induced regulation of TDP-43.

### C-terminal SUMO-mimetic destabilizes TDP-43

Promoting specific protein SUMOylation is rather difficult given the size of the SUMO molecule and the pleiotropy resulting from overexpressing SUMO proteins and their related machinery. To overcome this limitation and to mimic SUMOylation at K408, we expressed TDP-43-HA and TDP-43 with a C-terminal SUMO2 fusion (TDP-SUMO2-HA) in HEK293T cells. By western blot we found that C-terminal fusion of SUMO2 led to a significant reduction in TDP-43 levels (Fig. S3A). The reduction in TDP-43-SUMO2-HA suggesting that C-terminal fusion of SUMO2 destabilizes TDP-43 levels, consistent with our data showing that prolonged TDP-43 SUMOylation leads to its polyubiquitination and degradation (Fig. 1F). We further observed that overexpression of TDP-43-HA and TDP-43-SUMO2-HA both significantly increased phosphorylation of eIF2α compared to non-transfected controls despite significantly lower levels of TDP-43-SUMO2-HA.

Next, we aimed to gain insight into how C-terminal fusion of SUMO effects TDP-43 localization and stress response. We did not observe constitutive changes in TDP-43 localization suggesting that SUMO2 fusion does not drive TDP-43 nuclear export (Fig. S3B). By stressing cells for 1 hour with sodium arsenite we observed that TDP-43-SUMO-HA cells could elicit a stress response and form stress granules just like their WT counterparts. However, we were surprised to find a significant increase in the proportion of stress granules colocalizing with TDP-43-SUMO2-HA compared to TDP-43-HA alone. In addition, stress granules were generally larger and more irregular in size and shape in cells expressing TDP-43-SUMO2-HA. Taken together this suggests that SUMOylation of TDP-43 at the C-terminus converges on stress response pathways involving stress granules and the integrated stress response (i.e. eIF2α pathway). However, whether TDP-43 SUMOylation is involved in the upstream regulation or downstream modulation of these pathways remains unclear.

### Endogenous mutation of murine TDP-43 at K408 blocks stress-responsive TDP-43 SUMOylation in the mouse brain

To understand the cellular and physiological roles of TDP-43 SUMOylation, we generated a knock in mouse model harboring a missense (c.1223 A>G) point mutation in the endogenous mouse *Tardbp* locus resulting in the expression of Tdp-43 bearing a p.K408R missense mutation thereby blocking endogenous Tdp-43 SUMOylation (Fig. 2E, Fig. S4A,B). For simplicity herein, all genes/proteins will be referred to following their human nomenclature (e.g. mouse Tdp-43 versus human TDP-43, collectively “TDP-43”). Although TDP-43 and SUMO2 are independently essential for embryonic development, the “TDP-43^K408R^” mice are born at normal mendelian ratios without gross impairment indicating that TDP-43 SUMOylation at K408 is not essential for murine development (Fig. S4C)[54, 91].

In order to validate that mutation of endogenous TDP-43 K408 blocks stress responsive TDP-43 SUMOylation in TDP-43^K408R^ mice, we designed an ex vivo approach to test endogenous stress responsive TDP-43 SUMOylation in the mouse brain (Fig. 2F). Briefly, we dissected mouse embryos at E14 and divided each brain into two hemispheres. Both hemispheres were maintained in neurobasal complete media for one hour, with one hemisphere at 37 °C as a control, and the complementary hemisphere was heat shocked at 42 °C. The brain hemispheres were lysed under denaturing conditions and endogenously SUMOylated proteins were immunoprecipitated using antibodies targeted against endogenous SUMO2/3. SUMOylated proteins were eluted using a synthetic peptide reflecting the antibody epitope to competitively elute SUMOylated proteins. We observed that this approach reliably detects endogenous TDP-43 SUMOylation in response to heat shock yielding a doublet band at ∼65kDa and a corresponding decrease by ∼50% in TDP-43^K408R^ animals reflecting previous results in HEK293T cells (Fig. 2G, 1B). Thus, stress-induced TDP-43 SUMOylation is impaired in the TDP-43^K408R^ mouse brain.

### TDP-43 SUMOylation at K408 supports cellular stress response and recovery in neurons

As TDP-43 SUMOylation responds to, and is cleared during cellular stress and recovery respectively, we posited that SUMOylated TDP-43 plays key roles in responding and recovering from stress. We cultured murine TDP-43^+/+^ and TDP-43^K408R/K408R^ primary cortical neurons and performed stress and recovery assays to determine how the cell stress response may be affected. Since TDP-43 is critical for the formation of G3BP1 positive stress granules, we examined their formation and resolution following stress and recovery[92, 93]. We found that stress granules could form in TDP-43^K408R/K408R^ neurons to the same degree as TDP-43^+/+^ neurons, but there was a significant delay in stress granule clearance during recovery (Fig. 3A, Fig. S5A-C). When assessing TDP-43 solubility, we also observed a significant increase in RIPA insoluble phosphorylated TDP-43 at 1 hour recovery that was cleared by 3 hours post-stress recovery (Fig. 3B, S6A). Based on our previous results that SUMOylation is cleared by the UPS pathway and phosphorylation may function antagonistically with SUMOylation, the increase in phosphorylation during recovery may serve as an alternative method of proteostasis to compensate for loss of SUMOylation. We further observed a significant increase in nuclear TDP-43 foci that were specific to the recovery phase and were exacerbated after 3 hours recovery in the TDP-43^K408R/K408R^ neurons (Fig. 3C). Various forms of nuclear TDP-43 bodies are increasingly recognized to be aberrantly regulated in disease and stress states [19, 94–96]. We tested several likely markers of nuclear foci related to cell stress and recovery and found that these foci are neither anisosomes nor paraspeckles and they do not colocalize with Ubiquitin (Fig. S6B-D). Thus, these TDP-43 foci may represent a unique nuclear body related to recovery from cellular stress. Together, TDP-43^K408R/K408R^ neurons appropriately respond to, but not efficiently resolve acute cellular stress which may render cells vulnerable to further insult.

**Fig. 3:**
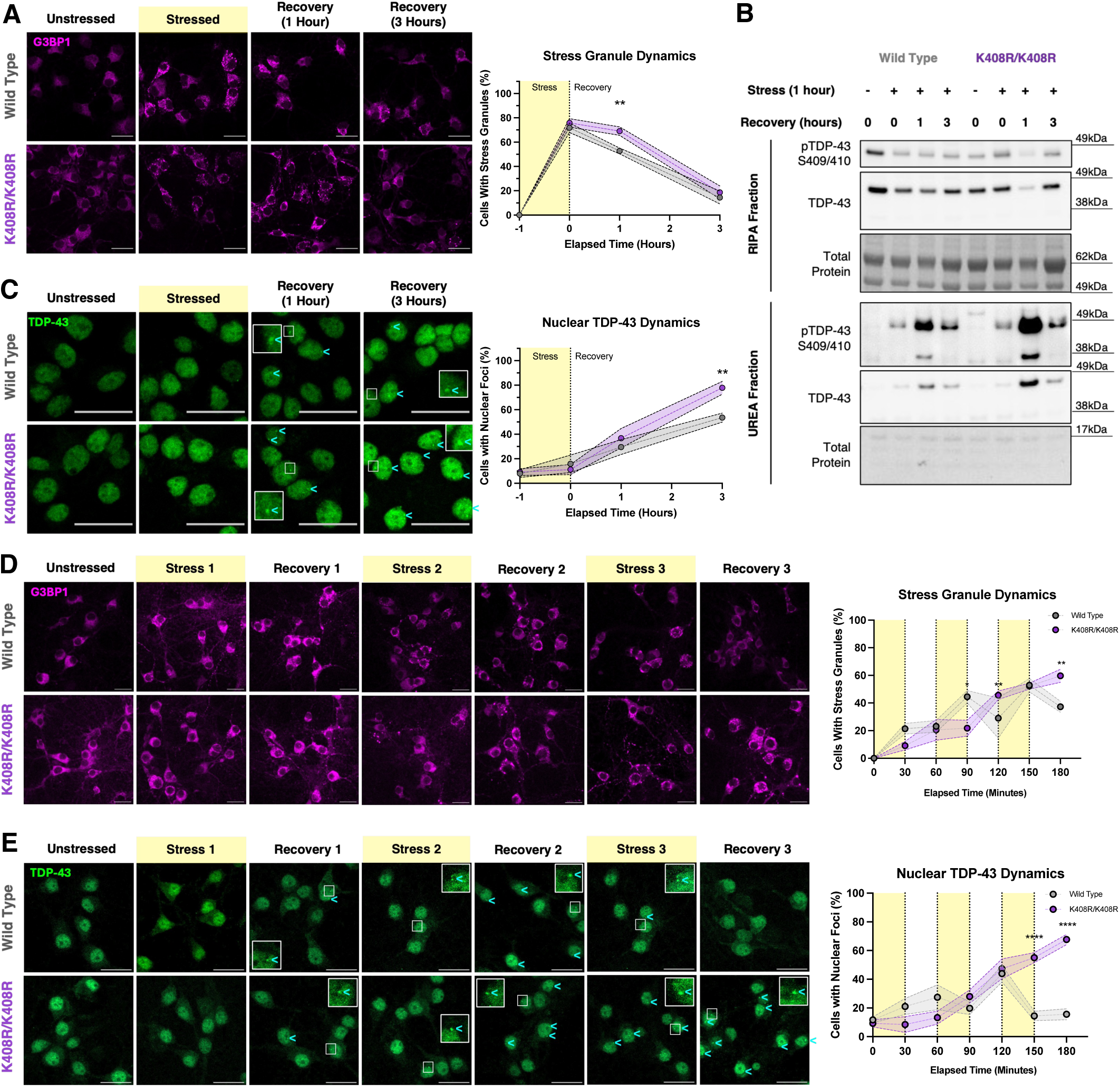
Blocking TDP-43 SUMOylation at K408 impairs the cellular stress response in neurons. **(A)** Representative images and quantification of G3BP1 stress granule dynamics in mouse primary cortical neurons (7 DIV) during stress (1 hour 250 μM sodium arsenite) and recovery. G3BP1 contrast was set for optimal visualization of stress granules. Scale Bar = 25 μm N=5, 2-Way ANOVA with Fisher’s LSD test data presented as mean ± SEM, * p<0.05, ** p<0.005. **(B)** Representative western blot and quantification of RIPA and UREA fractions from mouse primary cortical neurons (7 DIV) during stress (1 hour 250 μM sodium arsenite) and recovery. Quantification found in Fig. S6A. **(C)** Representative images and quantification of TDP-43 nuclear foci formation in mouse primary cortical neurons (7 DIV) during stress (1 hour 250 μM sodium arsenite) and recovery. Cyan arrowheads denote cells with nuclear TDP-43 foci. Scale Bar = 25 μm. N=5, 2-Way ANOVA with Fisher’s LSD test data presented as mean ± SEM, ** p<0.005. **(D)** Representative images and quantification of G3BP1 stress granule dynamics in mouse primary cortical neurons (7 DIV) during repeated stress (30-minute 250 μM sodium arsenite treatment) and recovery (30-minute washout), repeated 3 times. Scale Bar = 25 μm. N=3, 2-Way ANOVA with Fisher’s LSD test data presented as mean ± SEM, * p<0.05, ** p<0.005. **(E)** Representative images and quantification of TDP-43 nuclear foci formation in mouse primary cortical neurons (7 DIV) during repeated stress (30-minute 250 μM sodium arsenite treatment) and recovery (30-minute washout), repeated 3 times. Cyan arrowheads denote cells with nuclear TDP-43 foci. Scale Bar = 25 μm. N=3, 2-Way ANOVA with Fisher’s LSD test data presented as mean ± SEM, ****p<0.0001. For all immunofluorescent assays at least 50 cells were imaged and quantified per replicate (N=3-5).

Following the idea that SUMOylation of TDP-43 helps facilitate efficient recovery to protect cells from further insults, we questioned whether multiple short insults exacerbate the stress response in neurons. We designed a paradigm where primary cortical neurons were stressed for 30 minutes with sodium arsenite then allowed to recover for 30 minutes before receiving subsequent insults. We stressed and recovered the same neurons a total of 3 times over 3 hours and assessed stress granule dynamics in SUMO-competent (TDP-43^+/+^) and SUMOylation-deficient (TDP-43^K408R/K408R^) neurons. In TDP-43^+/+^ neurons we observed that during the first stress phase, stress granules formed in a proportion of neurons which were then retained during the first recovery (Fig. 3D). Subsequent rounds of stress further increased the proportion of cells containing stress granules which were roughly maintained or decreased during subsequent rounds of recovery suggesting that recovery pathways are being engaged and subsequent stress induces an additive effect without full recovery. In TDP-43^K408R/K408R^ neurons however we observed the initiation of stress granule formation in the first 30 minutes of stress that followed by inappropriate reactivation and recovery following subsequent rounds of stress. This supports that stress responses related to stress granule dynamics are altered in TDP-43^K408R/K408R^ likely related to inefficient recovery from stress. Additionally, we observed a striking increase in the proportion of cells presenting with nuclear TDP-43 foci after the third round of stress and recovery (Fig. 3E), though increases in TDP-43 mislocalization in this experimental paradigm were rather marginal and did not reach significance(Fig. S6E).

To better determine how blocking TDP-43 SUMOylation may respond to mild chronic stress, we treated neurons with low dose of sodium arsenite for 30 hours then allowed them to recovery for 3 days and assessed biochemical changes in TDP-43 solubility. In TDP-43^+/+^ neurons we observed a significant decrease in RIPA soluble TDP-43 after chronic stress followed by a marked increase in phosphorylated TDP-35 fragments in the urea-fraction (Fig. S6F). Upon recovery these fragments were cleared and insoluble TDP-43 returned to unstressed levels by 72 hours recovery. In TDP-43^K408R/K408R^ neurons however, we observed the same changes in response to chronic stress, however during recovery we observed significantly lower soluble TDP-43 levels and significantly increased phosphorylated TDP-35 fragments in the urea-fraction. Furthermore, we observed significant shift in the solubility of Ubiquitinated proteins during recovery. Taken together, blocking TDP-43 SUMOylation impairs the ability for cells to recover from cellular stress and may shift to other pathways involving phosphorylation or cleavage to regulate TDP-43.

### Blocking TDP-43 SUMOylation in vivo leads to sex-specific social and cognitive deficits

Following on our findings that blocking TDP-43 SUMOylation affect neurons abilities to recover from repeated or mild cellular stress, we posited that blocking TDP-43 SUMOylation in aging may render long lived cells (i.e. neurons) vulnerable across aging as mild life stresses accumulate over time [97]. Thus, we aimed to comprehensively characterize the TDP-43^K408R^ mice across their lifespan to uncover the baseline phenotypic changes that occur with aging as an innate stress upon cells. As TDP-43 SUMOylation is involved in supporting the normal cellular stress response and maintaining TDP-43 proteostasis, we hypothesized that disrupting this process leads to age-dependent physiological changes accelerated by impaired stress responses and TDP-43 accumulation. To ensure that the K408R mutation did not lead to post gestational developmental changes that would confound the presentation of age-related impairment, we assessed the weights and motor abilities of pups at P21. We did not observe any significant differences between TDP-43^+/+^ and TDP-43^K408R/K408R^ with respect to body weight or motor abilities indicating that blocking TDP-43 SUMOylation does not lead to gross developmental abnormalities (Fig. S7A-C).To determine whether blocking TDP-43 SUMOylation leads to phenotypes in an age-dependent manner, we assessed general wellness biweekly, including measuring body weight and hindlimb clasping scores, and performed a battery of 14 behavioral tests focusing on motor, social, and cognitive function related to ALS and FTD at three time points (2, 9 and 16 months, Fig. S7D-F, S8-10).Of note: TDP-43^K408R/+^ mice are generally omitted from figures for clarity to avoid confounding variables that may arise due to asymmetric regulation of wild type and K408R alleles due to the autoregulatory nature of TDP-43 (i.e. potential dominant negative effects conferred by the mutation and not partial loss of SUMOylation), but are accessible in Supplementary Table 2 and typically exhibit a milder phenotype than TDP-43^K408R/K408R^.

We were surprised to observe that blocking TDP-43 SUMOylation in TDP-43^K408R^ mice resulted in sex-specific cognitive and social deficits selective to female animals. Starting around 2 months of age, we observed significant barbering in female mice leading to alopecia in the K408R mutants (Fig. 4A). Consistent with this finding, we found that aged female TDP-43^K408R/K408R^ mice were significantly more submissive to their TDP-43^+/+^ counterparts in the tube test (Fig. 4B). Thus, blocking TDP-43 SUMOylation leads to social abnormalities in female mice. In assessing the cognitive behaviors, we observed significant risk taking and hyperactivity in the open field test at 2 months of age (Fig. 4C). This hyperactivity was age-dependent and selectively affected young female mice as they were significantly less active than their TDP-43^+/+^ counterparts at 9 months of age in the open field tests (Fig. 4C). Additionally, these female TDP-43^K408R/K408R^ mice presented with significant impairment during habituation at 9 months in the beam break test (Fig. 4D). As there was no change in motor abilities in the rotarod and digigait tests (Fig. S9C,D), we rationalized that the activity abnormalities were likely due to cognitive and anxiety impairments as opposed to motor impairments. By 16 months of age, female TDP-43^K408R/K408R^ mice performed significantly worse than TDP-43^+/+^ mice in the spontaneous Y-maze, inferring age-dependent cognitive decline in the domain of working memory (Fig. 4E). Together, female mice present with social and cognitive impairment in early age with slight impairments in cognitive performance with aging. There were no significant cognitive effects in male TDP-43^K408R/K408R^ mice, however mild impairments were observed on the rotarod test at 9 months of age (2-way ANOVA p=0.0273, Supplementary Table 2). This deficiency was not observed at 16 months of age, largely due to all mice performing significantly worse at the rotarod task, limiting the interpretation of this assay[98]. Taken together, the age-dependent cognitive and social impairment observed in female TDP-43 SUMO-deficient mutants highlight interesting and unexplored aspects of TDP-43 with important implications for understanding sex-specific roles of TDP-43 SUMOylation.

**Fig. 4:**
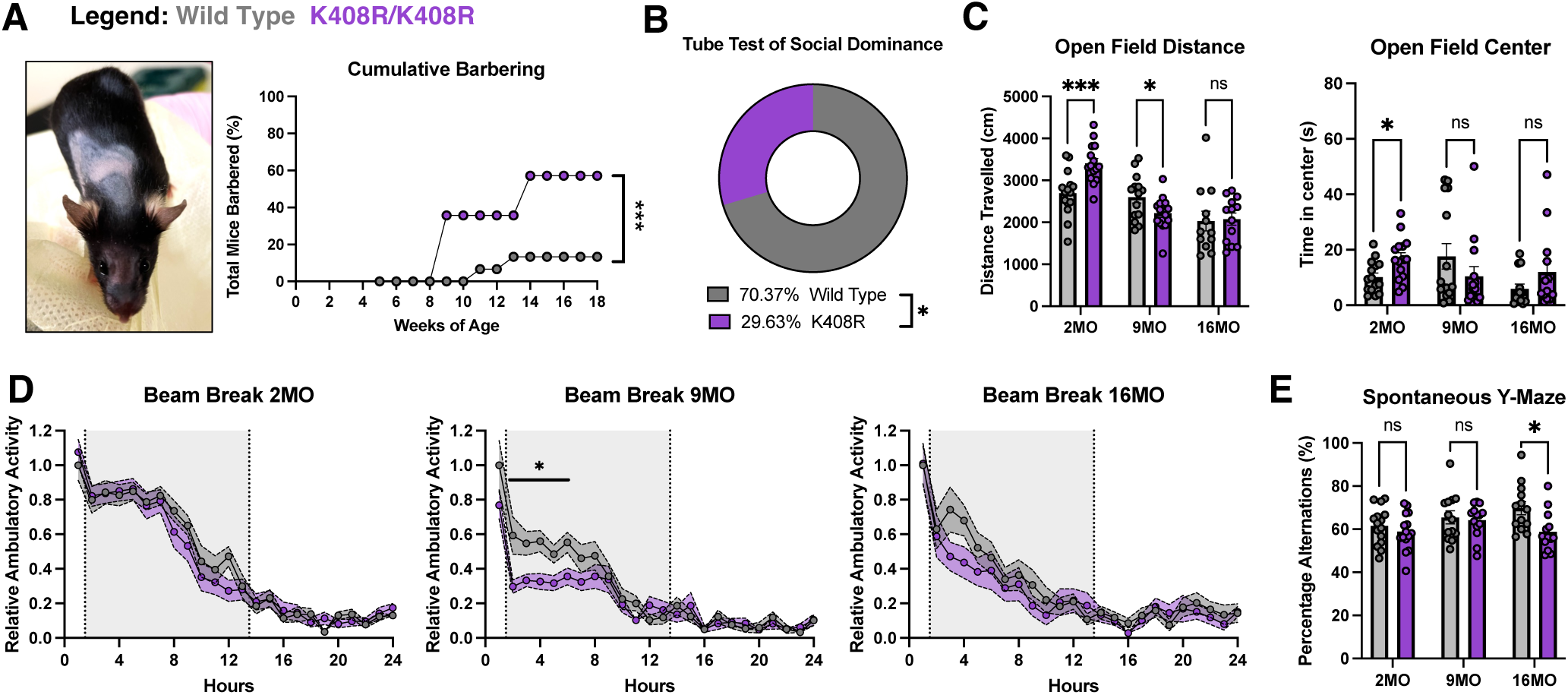
Blocking TDP-43 SUMOylation *in vivo* leads to sex-specific social and cognitive impairment. **(A)** Representative image of a barbered female mouse and quantification of cumulative barbering probability. 2-Way ANOVA with Tukey’s Multiple Comparisons, *** p<0.0005. **(B)** Tube test of social dominance at 16 months of age (16MO) displaying the total head-to-head battles and win percentage by genotype comparing wild type against K408R (K408R/+ and K408R/K408R animals). Binomial Test of Observed vs. Expected with Expected due to random chance set to 50%, * p<0.05. **(C)** Quantification of total distance traveled (cm) and total time in center (s) in the open field test. Mixed Effects Analysis with Tukey’s multiple comparison data presented as mean ± SEM, *** p<0.0005, * p<0.05. **(D)** Quantification of percent alternations in the spontaneous Y-maze test at 2, 9, and 16 months of age (2MO, 9MO, 16MO, respectively). Mixed-Effects Analysis with Tukey’s multiple comparison data presented as mean ± SEM, * p<0.05. **(E)** Quantification of relative ambulatory activity normalized to wild type at 1 hour at 2, 9, and 16 months of age (2MO, 9MO, and 16MO, respectively). Grey shading between hours 2 and 14 indicate “lights off” with respect to day/night light cycle. 2-Way ANOVA with Tukey’s multiple comparison data presented as mean ± SEM, * p<0.05.

### TDP-43^K408R^ mice present with distinct features of TDP-43 pathology related to ALS and FTD

Next, we aimed to characterize regions of the nervous system vulnerable in ALS/FTD to determine how loss of TDP-43 SUMOylation impacts these regions across aging. We did not observe significant differences in cortical thickness of the primary motor cortex nor prefrontal cortex in male or female mice across all timepoints suggesting that loss of TDP-43 SUMOylation does not lead to widespread cortical degeneration occurring during normal aging (Fig. S11A,B). To gain an additional layer of granularity into vulnerable neuron subpopulations in the cortex, we stained for Ctip2 to highlight neurons in layer V, including upper motor neurons, in the cortex and did not observe significant differences in neuron quantity across aging (Fig. S11C,D). We further stained for Cux1 as a marker of layer II/III neurons which can be vulnerable to TDP-43 pathology in FTD and degeneration in Alzheimer’s disease. Again, we did not observe significant changes in male nor female mice across the mouse lifespan supporting the absence of cortical degeneration in TDP-43^K408R^ animals during native aging (Fig. S11C,D). To determine if blocking TDP-43 SUMOylation results in neuroinflammation, we probed for the reactive astrocyte marker Gfap and reactive microglia marker Iba1 and observed no change in neuroinflammation (Fig. S11E-H). Together these data suggest the process of aging does not lead to cortical neurodegeneration nor cortical neuroinflammation.

Cellular and molecular changes are generally thought to precede behavioral changes in mouse models and human disease. Our biochemical and cellular assays show that stress-induced SUMOylation helps to clear TDP-43 during recovery and blocking TDP-43 SUMOylation in neurons leads to impaired recovery from stress and TDP-43 accumulation (Fig. 1E, Fig. 3, Fig. S6A,F). Due to the sex-specific social and cognitive impairment in female mice, we posited that TDP-43 accumulation would reflect the behavioural changes. Consistent with the sex-specific social and cognitive impairment described above, we find that insoluble phosphorylated TDP-43 accumulated in the cortex of female, but not male mice (Fig. S12A,B). Furthermore, this accumulation was observed as early as 2 months of age and was maintained throughout aging, which may imply that impaired TDP-43 proteostasis is underlying the social and cognitive phenotypes observed in female TDP-43^K408R^ animals. Interestingly, we observed significant changes in *Mapt* splicing in the cortex of TDP-43^K408R^ mice, specifically the N2/N0 splice isoform (Fig. S12C). Together, this may suggest that there is some degree of TDP-43 loss of function in the TDP-43^K408R^ mice.

We questioned whether motor neurons in the spinal cord present with early-stage pathology as they are subject to excess excitotoxic stress throughout their lifespan[40]. Surprisingly, we observed cytoplasmic mislocalization, but not nuclear depletion, of TDP-43 accompanied by insoluble, phosphorylated TDP-43 accumulation in the lumbar spinal cord of both male and female mice at 9 months of age (Fig. 5A, Fig. S12D,E). Furthermore, by 16 months of age, male TDP-43^K408R/K408R^ displayed a significant denervation of neuromuscular junctions in the tibialis anterior and a reduction in ChAT positive neurons in the spinal cord compared to littermate controls (Fig. 5B,C, Fig. S12F-H). Together, blocking TDP-43 SUMOylation results in altered proteostasis spinal motor neurons leading to neuromuscular junction denervation and ChAT positive motor neuron loss in an age dependent manner.

**Fig. 5:**
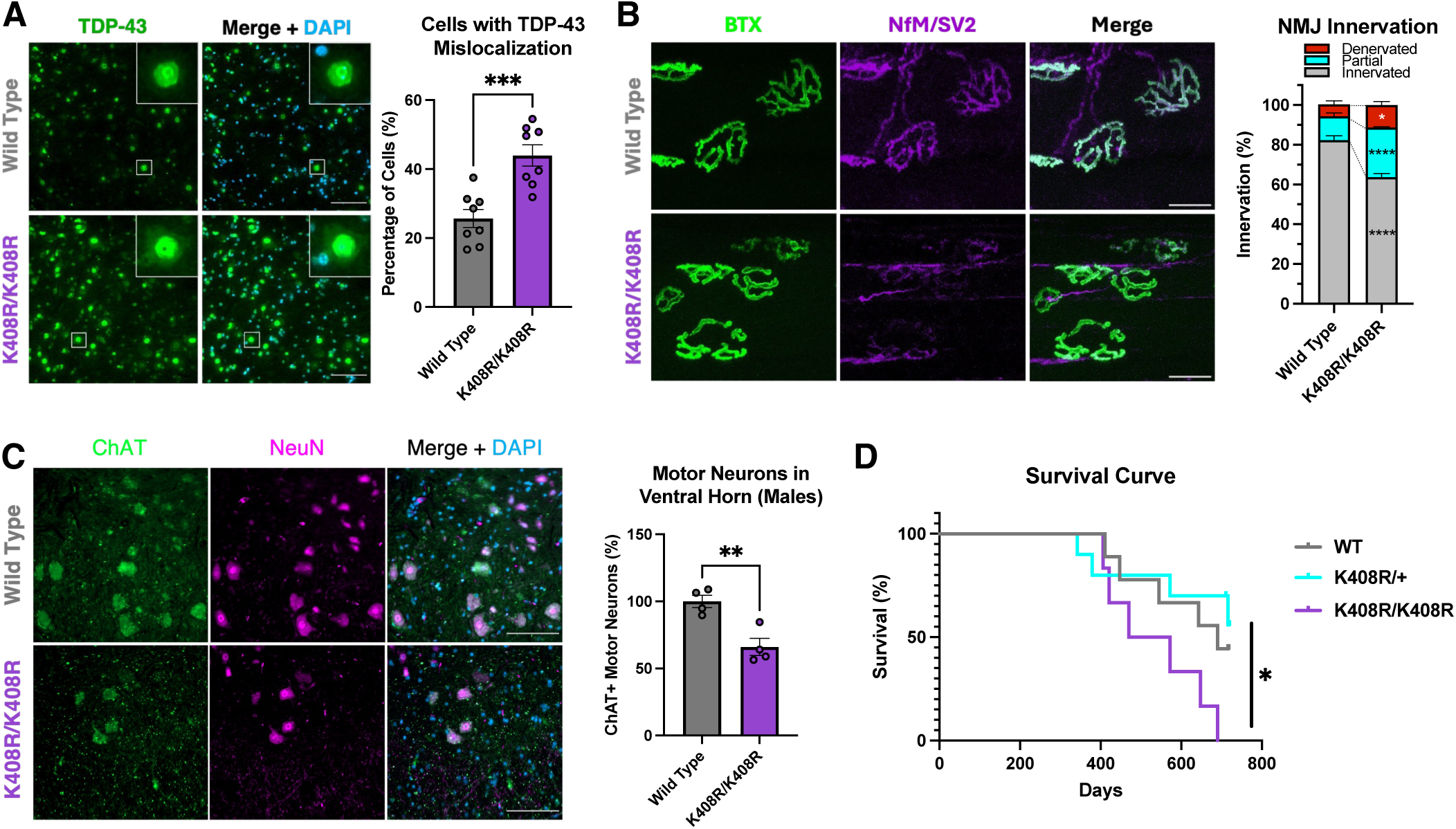
Male TDP-43^K408R^ mice present with age-specific features of ALS. **(A)** Representative images and quantification of TDP-43 mislocalization in the lumbar spinal cord of 9-month-old TDP-43^K408R^ mice. Each datapoint represents the average of 4 serial sections 40 μm apart per individual mouse. Scalebar = 100 μm. (N = 4 per sex/genotype) Unpaired t-test, *** p<0.0005. See Fig. S12C for sex comparisons. **(B)** Representative images and quantification of neuromuscular junction (NMJ) innervation in the tibialis anterior of male 16-month-old TDP-43^K408R^ mice. >80 NMJs were quantified per animal (N = 4 per genotype). 2-Way ANOVA with Tukey’s multiple comparison analysis, *p<0.05, ****p<0.0001. **(C)** Representative images and quantification of ChAT+ motor neurons in the ventral horn of the lumbar spinal cord of male mice. Each datapoint is the average of 4 serial sections spaced 40 μm apart through the lumbar enlargement of the lumbar spinal cord. Unpaired t-test, ** p<0.005. **(D)** Survival curve for male TDP-43^K408R^ mice (Females found in Fig. S7D). Curve comparisons analyzed using Log-Rank test and Gehan-Breslow-Wilcoxon test. *p<0.05.

As the TDP-43^K408R^ mice continued to age beyond our behaviour timepoints we observed a 24.49% decrease in survival specific to male TDP-43^K408R/K408R^ mice with a median survival of 521 days for the TDP-43^K408R/K408R^ compared to 690 days for TDP-43^+/+^ mice (Fig. 5D, Fig. S7D). Mice were typically found dead or euthanized after reaching a humane endpoint. Surprisingly, a few animals presented with classic hindlimb weakness paralysis akin to those observed in other models of ALS (Supplemental Video 1). However, these results should be interpreted with caution due to the low statistical power. Taken together, our data supports that TDP-43 SUMOylation plays a protective role in mediating TDP-43 proteostasis in the CNS and that its blockade may confer a risk for ALS/FTD-like pathogenesis in an age-dependent, and sex-specific manner providing insights into molecular substrates underlying sexually dimorphic clinical features of ALS and FTD.

### Aberrant SUMOylation is a feature of human aging and disease

Although SUMOylation is well understood to play an essential role in the CNS, its involvement in human aging and neurodegenerative diseases remains largely elusive. To assess how SUMOylation is normally affected in human aging, we processed samples from human temporal lobe tissue ranging between 0.5 and 88 years of age from individuals unaffected by neurological diseases (Fig. 6A, Supplementary Table 3, n=55). Complementing our findings that SUMOylation serves as a response to increased proteostatic demands, such as those in our aging TDP-43^K408R^ model, we observed that global SUMOylation was significantly increased in “aged” (>60 years old) human brains (Fig. 6B).

**Fig. 6:**
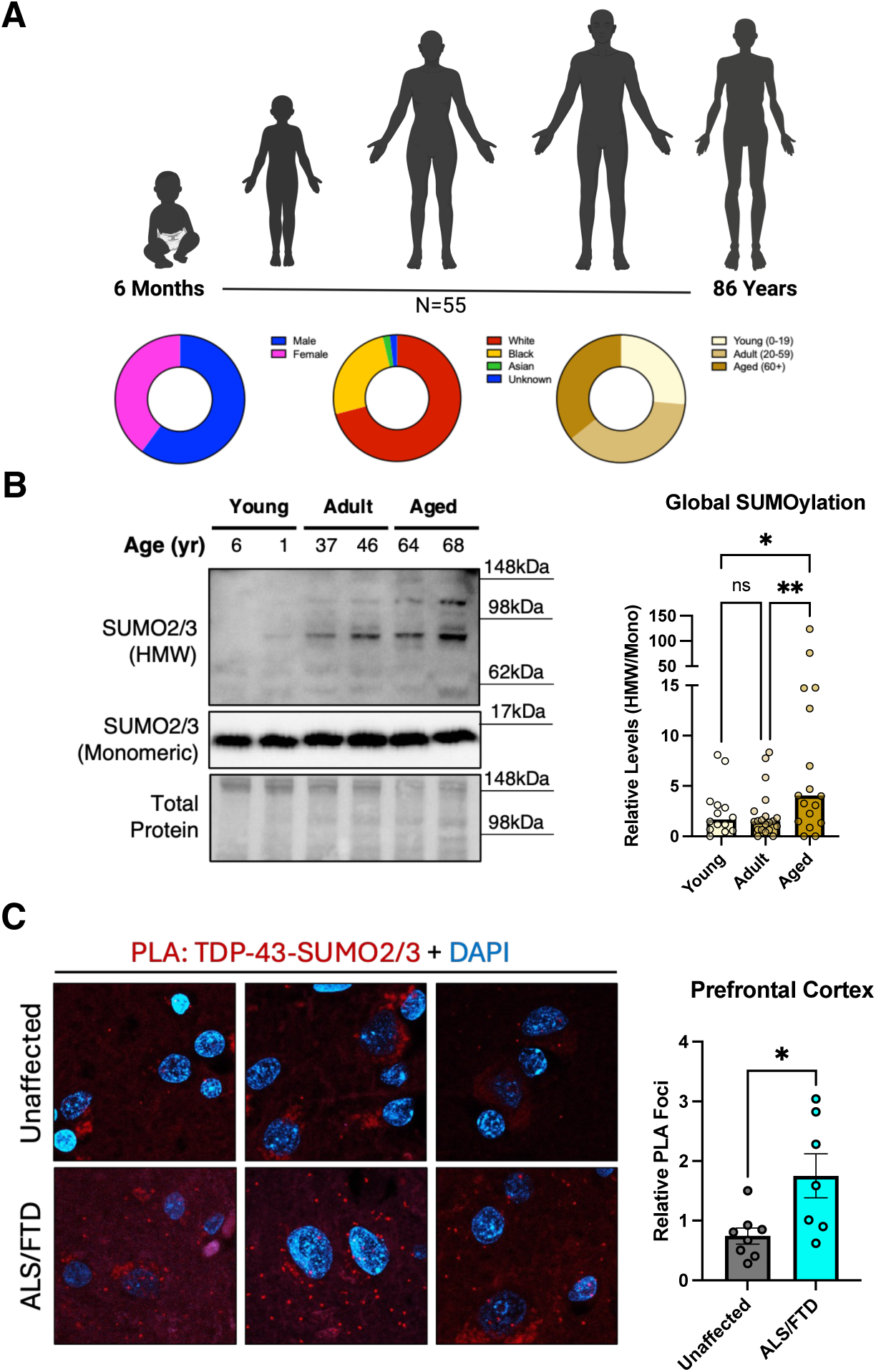
SUMOylation correlates with aging and is enriched in the prefrontal cortex of ALS/FTD patients. **(A)** Demographic composition of temporal lobe samples from aging cohort. **(B)** Representative western blot and box plot of SUMO2/3 across binned age groups. Average Age: “Young” = 6.32yrs, N = 14; “Adult” = 31.11yrs, N = 20; “Aged” = 64.00yrs, N = 19. Kruskall-Wallis with Uncorrected Dunn’s multiple comparisons test. * p<0.05, ** p<0.005. **(C)** Representative images and quantification from proximity ligation assay (PLA) between SUMO2/3 and TDP-43 in the prefrontal cortex from 3 patients diagnosed with ALS/FTD and unaffected controls. (N = 7-8) Unpaired T-test data presented as mean ± SEM, * p<0.05.

To explore the extent of TDP-43 SUMOylation in human ALS/FTD cases, we performed proximity ligation assays against TDP-43 and SUMO2/3 in prefrontal cortex samples from ALS/FTD cases positive for TDP-43 pathology and age/sex matched unaffected controls (Supplementary Table 3). Through semiquantitative analysis, patient samples collected from various sites were assessed for relative TDP-43 pathology by a neuropathologist in a blinded manner. We confirmed there was a significant increase in TDP-43 pathology in the prefrontal cortex of ALS/FTD patients relative to controls (Fig. S13A). When assessing the interaction between TDP-43 and SUMO2/3, we observed a significant increase in interactions in the prefrontal cortex of ALS/FTD patients with TDP-43 pathology suggesting the TDP-43 SUMOylation pathway is engaged in disease (Fig. 6C, Fig. S13B). This may be a result of increased stress burden throughout the prefrontal cortex in those affected by ALS/FTD with TDP-43 pathology compared to age and sex matched counterparts. Taken together, SUMOylation as a stress responsive modification is positively correlated with aging in the human brain, potentially linked to the increase demand imposed by age-associated stress. Additionally, the SUMOylation response is enhanced in the frontal cortex of those affected by ALS/FTD providing further *in vivo* support for its role in disease.

## Discussion

Although ALS and FTD are highly heterogenous diseases, the nearly universal convergence on TDP-43 proteinopathy emphasizes a need to understand early drivers of this pathogenic process. Mutations in various genes can lead to TDP-43 pathology, however the majority of ALS and FTD cases occur sporadically without a clear genetic cause[3, 22].

Additionally, patients living with disease causing mutations do not present with symptoms of disease until later in life suggesting exogenous factors related to aging play a role in phenoconversion[99]. Cellular stressors are well characterized to induce TDP-43 pathology providing a clue as to how exogenous factors may contribute to disease pathogenesis. Given the postmitotic nature of neurons, maintaining proteostasis in response to lifelong stressors is critical for neuronal longevity[1]. This is particularly true for motor neurons which endure constant excitotoxic stress[40]. In this study, we found that SUMOylation maintains TDP-43 proteostasis in the nucleus specifically in response to stress. This modification is critical to preserve neuronal function, the absence of which results in neuronal demise and TDP-43 proteinopathy.

SUMOylation of TDP-43 occurs at K408 in a highly conserved region of the C-terminal domain. This is in contrast to what was previously reported for TDP-43 and SUMO1; where it was suggested, but not demonstrated, that SUMO1 modified TDP-43 at lysine 136[49, 100]. Indeed, mutating K136 to an arginine (R) had no effect on stress-induced TDP-43 SUMOylation by SUMO2/3, though this does not discount the possibility that SUMO1 may modify TDP-43 under conditions where increased SUMO machinery (e.g. UBC9 overexpression) is present. In our hands, expression of TDP-43^K136R^ led to the formation of nuclear TDP-43 puncta as previously reported[50, 51]. However, these puncta were similar to those observed when expressing RNA binding deficient TDP-43. Furthermore, prediction of TDP-43 structure suggests that K136R mutation may impact the structure of the RNA recognition motif and RNA binding. Thus, phenotypes observed in TDP-43 carrying the K136R mutation should be interpreted with caution.

In genetic forms of ALS or FTD caused by mutations in TDP-43, the C-terminal domain harbors the overwhelming majority of disease-causing mutations[101]. Thus, understanding the involvement of the C-terminal domain in TDP-43 biology is critical to uncover pathways that may be linked to disease independent of TDP-43 mutations. Phosphorylation of TDP-43 in the C-terminal domain, particularly S403/404 and S409/410, have garnered much attention in this regard as these residues are characteristically phosphorylated in disease [42]. However, the roles for phosphorylation at these sites remains generally elusive. Few reports have suggested roles in regulating liquid-liquid phase separation and solubility[102, 103]. Our findings suggest that phosphorylation and SUMOylation act antagonistically on TDP-43, indicating potential distinct mechanisms of regulation. We further observed that this region of the C-terminal domain targeted by SUMOylation and phosphorylation is highly conserved throughout vertebrate species. Thus, this PTM-enriched region may play key roles for regulating TDP-43 in vertebrate biology.

Regulation of TDP-43 proteostasis is critical for cells as fluctuations in TDP-43 levels can lead to toxicity in cell and animal models[20, 104–107]. Unsurprisingly, TDP-43 is regulated in a variety of mechanisms including transcriptionally, post-transcriptionally, translationally and post-translationally[11, 20, 108]. Furthermore TDP-43 can be cleared through several pathways including the proteasome, endo-lysosome, autophagy, and extracellular vesicle pathways[109–116]. Here we add another mechanism of regulation through SUMOylation during stress and recovery. We hypothesized that blocking TDP-43 SUMOylation with K408R mutation would result in its accumulation during stress recovery. Interestingly, we observed a significant increase in insoluble phosphorylated TDP-43 in TDP-43^K408R/K408R^ cortical neurons and a delay in stress granule disassembly during stress recovery indicating TDP-43 accumulation. However, this was resolved 3 hours post-stress indicating that the neurons could compensate to clear the aberrantly accumulated TDP-43 within this acute stress paradigm. We questioned whether chronic stress may exacerbate phenotypes and found that chronic stress significantly resulted in increased phosphorylated, insoluble TDP-35 fragments which then recovered upon removal of stress. Thus phosphorylation of TDP-43, in addition to cleavage, may be additional mechanisms to regulate TDP-43 proteostasis in the event of SUMOylation-pathway failure. Thus, SUMOylation may be an early line of defense to regulate TDP-43 in response to stress and blocking TDP-43 SUMOylation shifts proteostasis leading to delayed recovery. Future studies exploring repeated and/or divergent stressors will be crucial in determining the degree to which TDP-43 SUMOylation safeguards from cellular demise.

Previous reports have emphasized the status of altered proteostasis as a hallmark of aging in the CNS, culminating in impaired removal of damaged proteins with age[97]. In TDP-43^K408R/K408R^ mice, TDP-43 was found to accumulate in the CNS in an age-dependent manner, suggesting SUMOylation plays important roles during aging. Supporting these findings, in human temporal lobe samples we similarly observed a positive correlation between global SUMOylation and age supporting SUMOylation as a potential protective mechanism during aging in the CNS. Previous studies on global SUMOylation have focused on the dynamic nature of this modification playing roles during development, and associations between SUMOylation and neurodegeneration gaining traction however roles for SUMOylation in the central nervous system during the process of human aging remain understudied[117, 118]. This may be ascribed, in part, to challenges in studying SUMOylation due to the dynamic nature of the modification and lack of specific tools to study its interactions. We employed a PLA approach to assess the interaction between TDP-43 and SUMO in ALS/FTD patients and found significantly increased interactions in the prefrontal cortex relative to age/sex matched unaffected controls. It was interesting to observe that PLA signal occurred and was increased in both the nucleus and the cytoplasm in ALS/FTD patient samples. We hypothesize that SUMOylation of TDP-43 is an early event regulating TDP-43 proteostasis, however if overwhelmed other systems such as phosphorylation or protein cleavage may help respond to regulate TDP-43. As we are observing late-stage events, SUMOylation in the cytoplasm may be co-occurrent with late-stage regulation of TDP-43. Approaches that provide increased resolution for detecting TDP-43 SUMOylation events are clearly warranted. Recent studies have used RNA aptamers to reveal nuclear changes in TDP-43 in spinal and cortical neurons of ALS patients prior to traditional cytoplasmic pathology in vulnerable cells[19]. Further development of these technologies with respect to TDP-43 SUMOylation may help advance biomarker development that can provide information about TDP-43 stress-response and pathogenesis in ALS/FTD patients.

While the extent of phenotypes observed in TDP-43^K408R^ are relatively mild, they similarly reflect the magnitude of phenotypes observed in mice with significant construct validity to ALS/FTD (i.e. TDP-43^Q331K^, TDP-43^M337V^, and TDP-43^K145R^ models) [45, 119, 120]. These knock in models highlight potential shortcomings of other TDP-43 models relying on overexpression to induce behavioural phenotypes in mice which may not reflect the underlying biology of TDP-43 in ALS/FTD. The emergence of predominantly cognitive FTD-like phenotypes in the TDP-43^K408R^ mice is not unusual in TDP-43 knock in models such as TDP-43^Q331K^ and TDP-43^K145R^ mice which present with features of cognitive dysfunction but lacking the entire constellation of ALS/FTD-like phenotypes[45, 119]. However, we were surprised to observe that the cognitive phenotypes were specific to female mice with no behavioural phenotypes being observed in male mice. This may imply that TDP-43 plays important role regulating sex specific behaviours. Alternatively, female and male mice may experience different forms of life stress that may act upon SUMOylation pathways. For example, we observed differences in social behaviors which may indicate social pathways linked with female behaviour may also be associated with SUMOylation. Alternatively, male mice were frequently separated due to fighting behavior, which could contribute to altered baseline social features, regardless of genotype. Future work exploring sex-specific roles of TDP-43 and/or SUMOylation will help provide mechanistic insight into sex-specific behaviour and experience.

As TDP-43 SUMOylation plays roles in regulating TDP-43 clearance during recovery from stress, blocking TDP-43 SUMOylation in the TDP-43^K408R/K408R^ mouse line disrupts this proteostasis and leads to subsequent mislocalization consistent with loss of nuclear function. Additionally, we observe significant increases in insoluble phosphorylated TDP-43 and motor neuron loss in male mice suggesting that gain of function may be occurring, phenocopying disease pathogenesis. While we did observe some motor impairment in 9-month-old mice correlating with TDP-43 mislocalization, we did not observe significant motor impairment in the male mice at 16 months of age, which may be explained by the overall poor performance during motor tests at this age limiting the sensitivity to detect mild changes. Despite this, we did observe molecular and histological evidence supporting early motor impairment with significant neuromuscular junction denervation and reduction in ChAT positive motor neurons in the ventral horn. Additionally, as the male TDP-43^K408R/K408R^ mice displayed a significant decrease in survival, behavioral analysis at 16 months of age may have preceded the onset of significant motor phenotypes. Future refined studies using key pathological centered around key pathological markers of cellular dysfunction will help uncover specific points of phenoconversion and phenotransition to identify the chronology of events leading to phenotypes associated with neurodegeneration.

We have demonstrated that TDP-43 SUMOylation occurs in response to cellular stress, thus careful characterization of the TDP-43^K408R^ mice is critical to determine baseline phenotypes without the addition of exogenous stressors. We initially hypothesized that aging may be a sufficient stressor to interrogate TDP-43 SUMOylation in mice leading to age-dependent phenotypes. However, the various stressors that individuals with ALS/FTD were previously exposed to during their lifetime (i.e. the “exposome”) are not experienced by mice in vivaria on a short timescale thus mouse aging likely does not faithfully reflect human aging. The TDP-43^K408R^ model will therefore help facilitate the exploration of how the exposome synergizes with TDP-43 to better understand how ALS/FTD relevant stressors that humans may experience throughout aging (e.g. *C9ORF72* expansions, viral infections, and traumatic brain injury) converge on TDP-43 and uniquely drive aspects of neurodegeneration.

## Acknowledgements

This project was supported in part by an NSERC Discovery Grant and Discovery Launch Supplement to M.W.C.R. (RGPIN-2019-04133 and DGECR-2019-00369); the Canada Research Chairs program to M.W.C.R; the ALS Society of Canada in partnership with the Brain Canada Foundation through the Brain Canada Research Fund, with the financial support of Health Canada, for financial support through the ALS Trainee Award Program 2019 (T.R.S.) and the Discovery Grants Program 2021 (M.W.C.R.); the J.P. Bickell Medical Research Fund (M.W.C.R.); Canadian Institute of Health Research Project Grant (PJT-195691, M.W.C.R.). The Eric Poulin Center for Neuromuscular Disease through the Student Translational Research Awards (T.R.S., V.S.G.); The Canadian Institute of Health Research through the CGS-M award (C.E.P., J.L.Z., V.S.G.); The Ontario Graduate Scholarship Program Award (C.E.P.); NSERC Undergraduate Summer Research Award (T.T.N.); Medical Student Summer Research Program Award (T.T.N); Rising Star in ALS Research 2024 Award in memory of Madeleine Blanc through the Brain Canada Foundation (J.L.Z.); Christopher Chiu senior postdoctoral fellowship (P.M.M.); James Hunter and Family ALS Initiative (J.R.). Human tissue was received from the NIH NeuroBioBank at the University of Maryland, University of Pittsburgh, and Sepulveda sites. The authors also thank all members of the Rousseaux lab for important discussions and critical feedback on the manuscript. The authors also thank the following Core facilities from the University of Ottawa and the Ottawa Hospital Research Institute (OHRI) for use of their facility, equipment, and expertise: the Cell Biology and Imaging Acquisition Core (RRID:SCR_021845), STEMCore Laboratories (RRID:SCR_012601), Genome Engineering and Molecular Biology (GEM) Core (RRID:SCR_022954), Animal Behaviour and Physiology Core (RRID:SCR_022882), Louise Pelletier Histology Core (RRID: SCR_021737), Figs. 1, 2, 6, and S4 were generated in part with Biorender.com.

## Author Contributions

Conceptualization: TRS, CEP, MWCR

Methodology: TRS, CEP, TTN, JLZ, VSG, PMM, BN, SMC, JR

Investigation: TRS, CEP, TTN, JLZ, MMH, ACG, VSG, PMM, BN, JR

Visualization: TRS, CEP, JLZ, MWCR

Funding acquisition: TRS, CEP, MWCR

Project administration: JMW, JR, MWCR

Supervision: JR, MWCR

Writing – original draft: TRS, MWCR

Writing – review & editing: TRS, MWCR

## Supplemental Figures

**Fig. S1:**
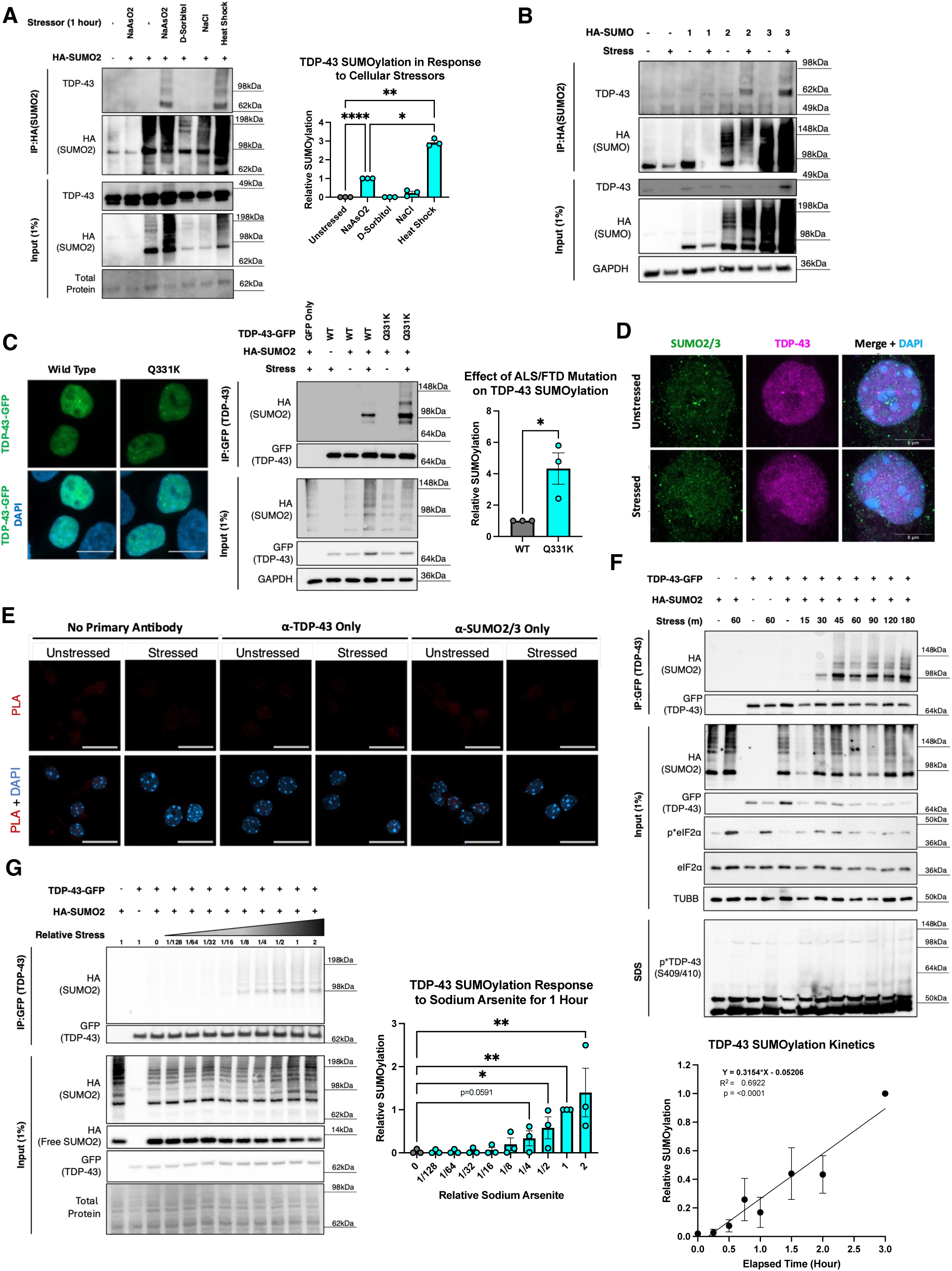
Characterizing stress responsive TDP-43 SUMOylation. **(A)** Representative western blot from SUMOylation assays testing whether various stressors that cause TDP-43 aggregation induce TDP-43 SUMOylation: NaAsO2 (250 μM), D-Sorbitol (400 mM), NaCl (300 mM), Heat Shock (42 °C) in HEK293T HA-SUMO2 stable cells. One-Way ANOVA with Fisher’s LSD test. Data presented as mean ± SEM. *p<0.05, **p<0.005, ****p<0.0001. **(B)** Representative western blot from SUMOylation assays testing the selectivity of SUMO paralogs to SUMOylate TDP-43 in response to 1 hour sodium arsenite (250 μM) in HEK293T cells with transient expression of SUMO paralogs. **(C)** Representative GFP-Trap SUMOylation assay depicting increased SUMOylation with ALS-causing Q331K mutation in response to 250 µM sodium arsenite stress in HEK293T cells with transient expression of HA-SUMO2. Unpaired T-test. *p<0.05. Data presented as mean ± SEM. **(D)** Representative images of SUMO2/3 and TDP-43 in the nucleus of mouse cortical neurons (7 DIV) in unstressed and 1 hour sodium arsenite (250μM) stressed conditions. **(E)** Representative control imaged from Proximity Ligation Assay (PLA) in mouse primary cortical neurons. Scalebar = 20 μm. (N = 4). **(F)** Representative western blot and quantification from GFP-Trap SUMOylation assays testing the kinetics of TDP-43 SUMOylation in response to sodium arsenite stress (250 μM) and its relationship to insoluble phosphorylated TDP-43 in HEK293T HA-SUMO2 stable cells. (N = 3). Linear regression analysis of Relative SUMOylation against time in response to stress. **(G)** Representative dose-response assay to detect TDP-43 SUMOylation after 1 hour of sodium arsenite stress (1 = 250 μM, 1/2 = 125 μM, 1/4 = 62.5 μM, etc.) in HEK293T HA-SUMO2 stable cells. One-Way ANOVA with Fisher’s LSD test. Data presented as mean ± SEM. *p<0.05, **p<0.005, ****p<0.0001.

**Fig. S2:**
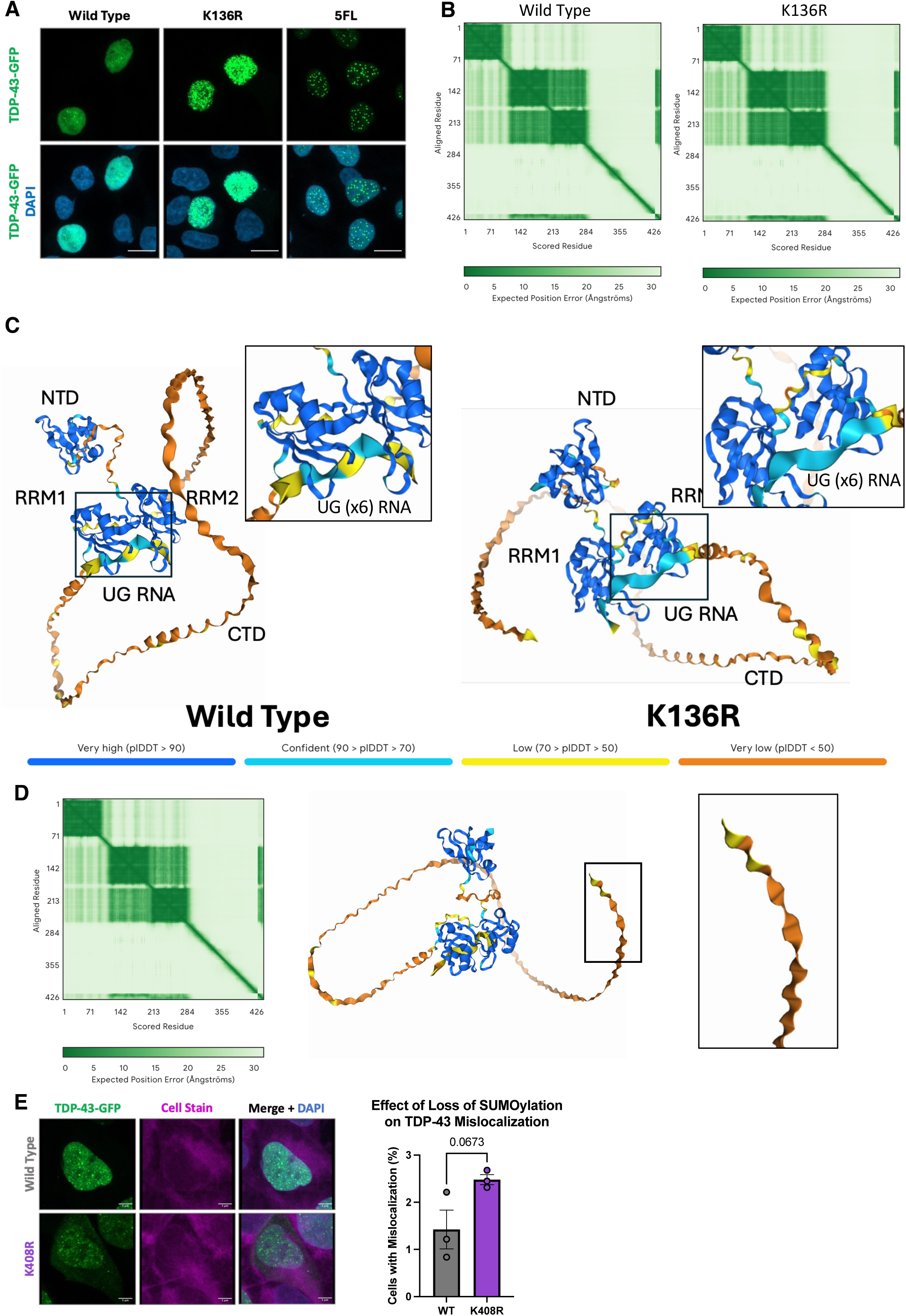
Analysis of TDP-43 lysine to arginine mutation at K136 and K408. **(A)** Representative fluorescent microscopy images expressing TDP-43-GFP with K136R and RNA binding deficient (5FL) mutations. Scale bar = 10 μm. **(B)** Predicted alignment error output from AlphaFold3 prediction of TDP-43 structure (residues 1-414) interacting with UG (x6) RNA (residues 415-426). **(C)** Predicted structure of TDP-43 with annotated domains. Color annotates predicted local difference test (plDDT)j scores. NTD = N-Terminal Domain; RRM1/RRM2 = RNA Recognition Motif 1/2; CTD = C-Terminal Domain. **(D)** Predicted structure of TDP-43^K408R^ highlighting no addition of secondary structure at the C-terminus surrounding residue 408 upon mutation. **(E)** Representative image and quantification of TDP-43-GFP expression denoting mislocalization in a subset of cells overexpressing TDP-43 with K408R mutation. (N=3) Students T-test. Data presented as mean ± SEM.

**Fig. S3:**
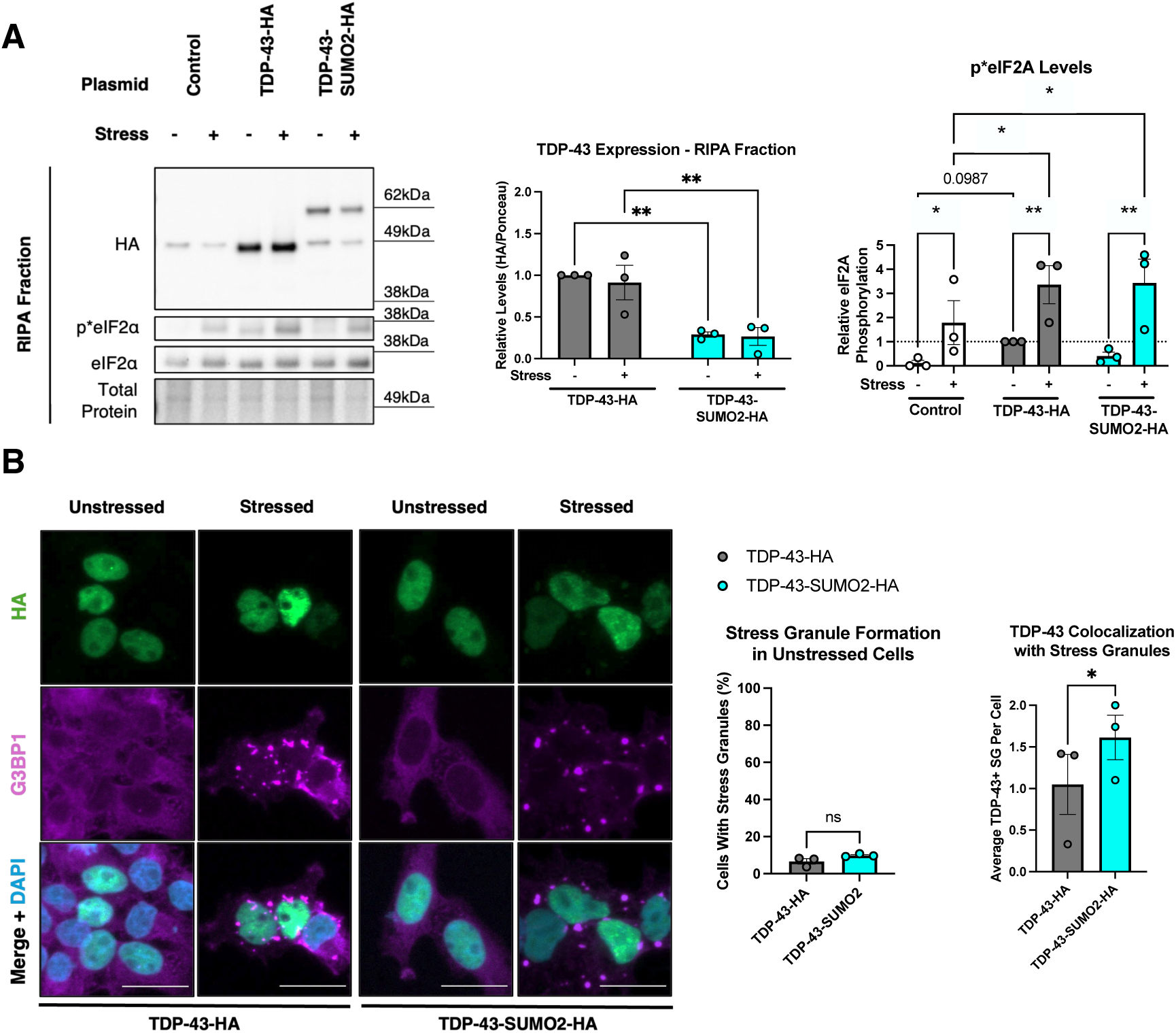
Characterization of SUMO2 fusion to the C-terminus of TDP-43. **(A)** Representative western blot and quantification of TDP-43-HA and TDP-43-SUMO2-HA expression in HEK293T cells. (N=3) 2-way ANOVA with Fishers LSD test. Data presented as mean ± SEM. *p<0.05, **p<0.005. **(B)** Representative immunofluorescence microscopy images of TDP-43-HA and TDP-43-SUMO2-HA fusion in HEK293T cells in response to cellular stress (250 μM sodium arsenite for 1 hour). (N=3) Students T-test. Data presented as mean ± SEM. *p<0.05.

**Fig. S4:**
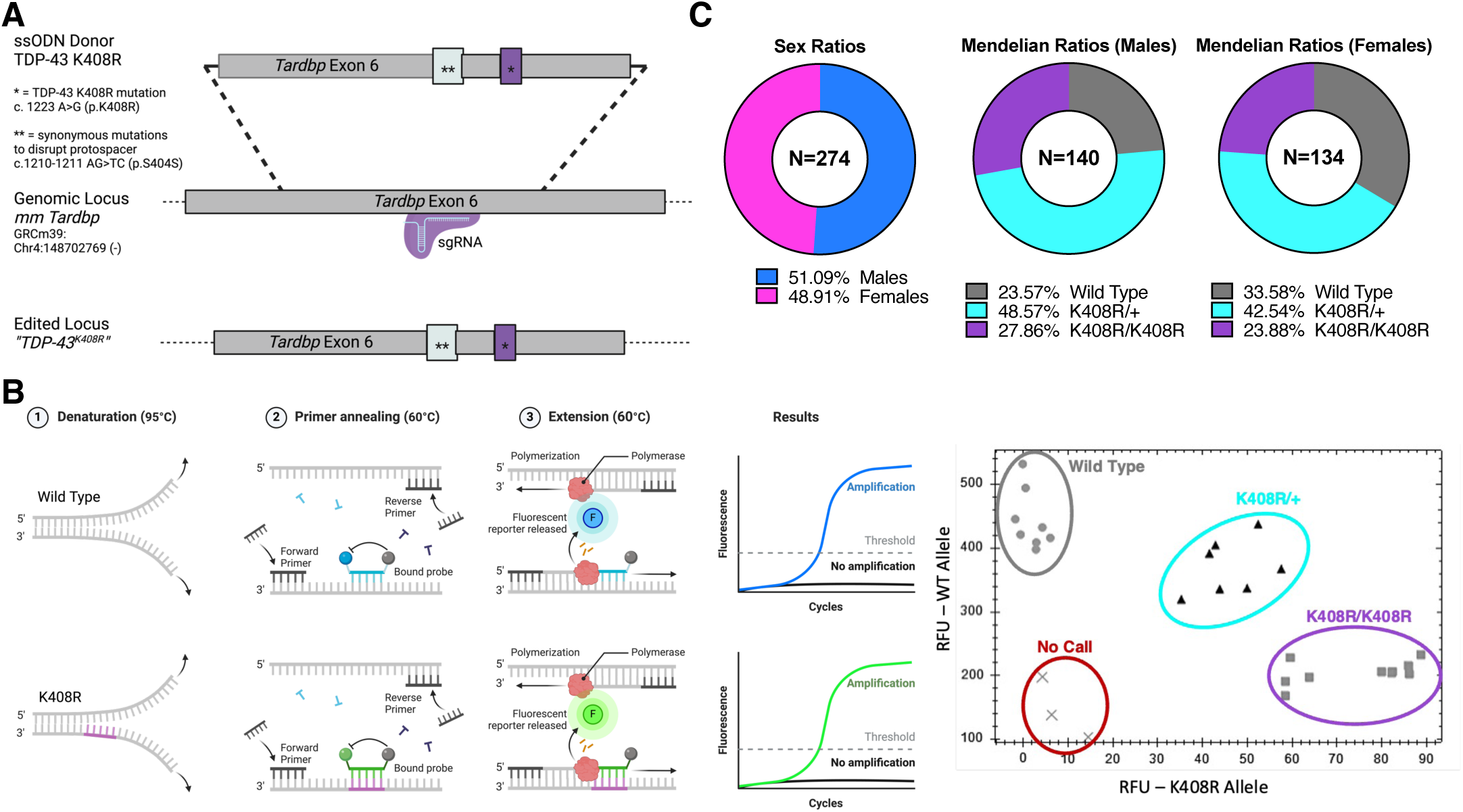
Generation of the TDP-43^K408R^ mouse line. **(A)** Schematic of the knock in approach to edit the endogenous *Tardbp* to express TDP-43^K408R^. **(B)** Schematic of LNA probe based genotyping approach and allelic discrimination assay to genotype TDP-43^K408R^ mouse line. **(C)** Sex and mendelian ratios of F3 generation TDP-43^K408R^ mice from K408R/+ and K408R/+ breeding pairs used to generate behaviour and histology cohorts. Observed vs Expected statistical analysis assuming an expected 1:1 ratio sex ratio and 1:2:1 mendelian ratio.

**Fig. S5:**
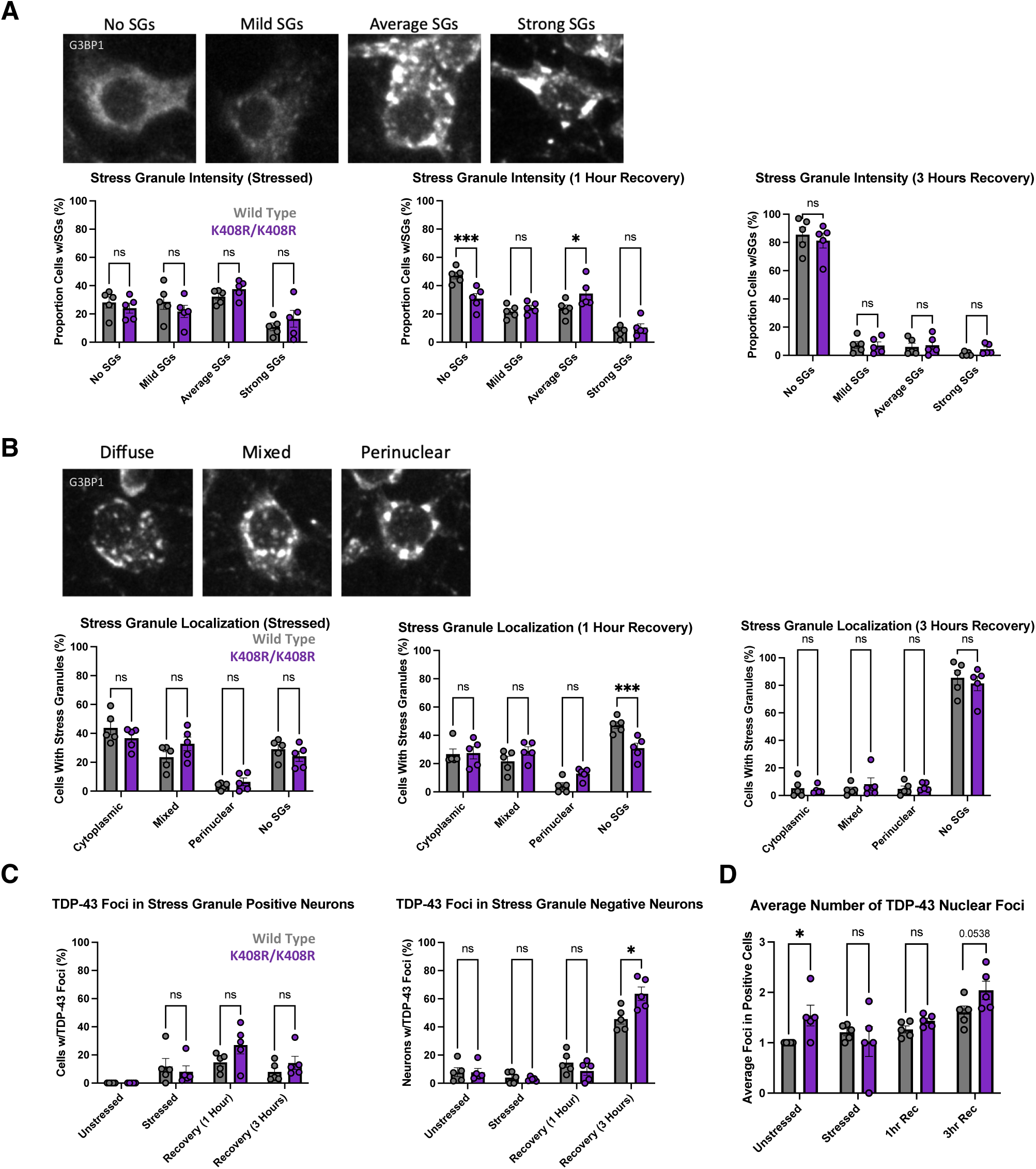
Characterizing the stress response in TDP-43^K408R^ primary cortical neuron cultures. **(A)** Classification and quantification of G3BP1 stress granule (SG) intensity in response to 1 hour sodium arsenite stress (250 μM) and recovery. Data presented as mean ± SEM. (N = 5 per genotype) 2-Way ANOVA with Tukey’s multiple comparisons analysis. *p<0.05, ***p<0.0005. **(B)** Quantification of the localization of G3BP1 stress granules (SGs) in response to 1 hour sodium arsenite stress (250 μM) and recovery. Data presented as mean ± SEM. (N = 5 per genotype) 2-Way ANOVA with Tukey’s multiple comparisons analysis. ***p<0.0005. **(C)** Quantification of the percentage of neurons with positive TDP-43 nuclear foci in G3BP1 stress granule positive and G3BP1 stress granule negative neurons in response to 1 hour sodium arsenite stress (250 μM) and recovery. Data presented as mean ± SEM. (N = 5 per genotype) 2-Way ANOVA with Tukey’s multiple comparisons analysis. ***p<0.0005. **(D)** Quantification of the average number of TDP-43 nuclear foci in neurons positive for TDP-43 nuclear foci in response to 1 hour sodium arsenite stress (250 μM) and recovery. Data presented as mean ± SEM. (N = 5 per genotype) 2-Way ANOVA with Tukey’s multiple comparisons analysis. *p<0.05.

**Fig. S6:**
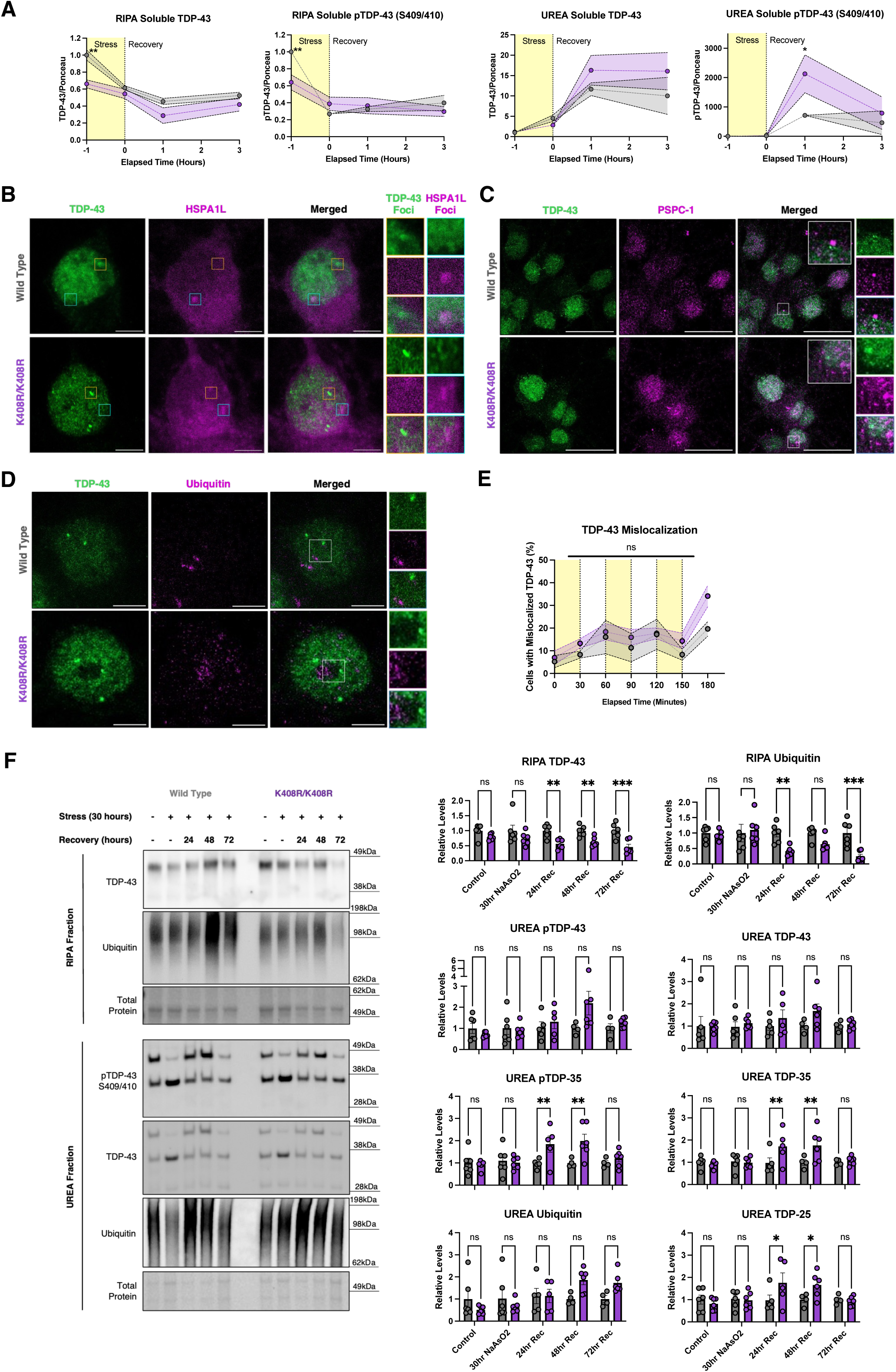
Characterizing the effect of blocking TDP-43 SUMOylation in primary cortical neurons. **(A)** Quantification of RIPA and UREA fractions from mouse primary cortical neurons (7 DIV) during stress (1 hour 250 μM sodium arsenite) and recovery related to Fig. 3B. N=3-5, 2-Way ANOVA with Fisher’s LSD test. Data presented as mean ± SEM, * p<0.05, ** p<0.005. **(B-D)** Representative immunofluorescent images of primary cortical neurons (7 DIV) treated with 250 μM sodium arsenite for 1 hour then recovered for 3 hours to induce TDP-43 foci formation to analyze colocalization with nuclear markers: (B) HSPA1L was visualized through lentiviral transduction of HSPA1L-mRuby2; (C) paraspeckle marker PSPC-1; and (D) Ubiquitin. (N=3). At least 10 cells positive for TDP-43 foci and foci of interest were imaged and analyzed per replicate. **(E)** Quantification of primary cortical neurons with TDP-43 mislocalization in response to repeated stress (30 minutes 250 μM sodium arsenite) and recovery (30 minutes washout) related to Fig. 3E. (N=3) 2-Way ANOVA with Fishers LSD test. Data presented as mean ± SEM. **(F)** Representative western blot and analysis of RIPA soluble and UREA fractions from primary cortical neurons in response to chronic stress (30 hours of 15 μM sodium arsenite) and recovery. Data presented as mean ± SEM. 2-Way ANOVA with Fishers LSD test. *p<0.05, **p<0.005.

**Fig S7:**
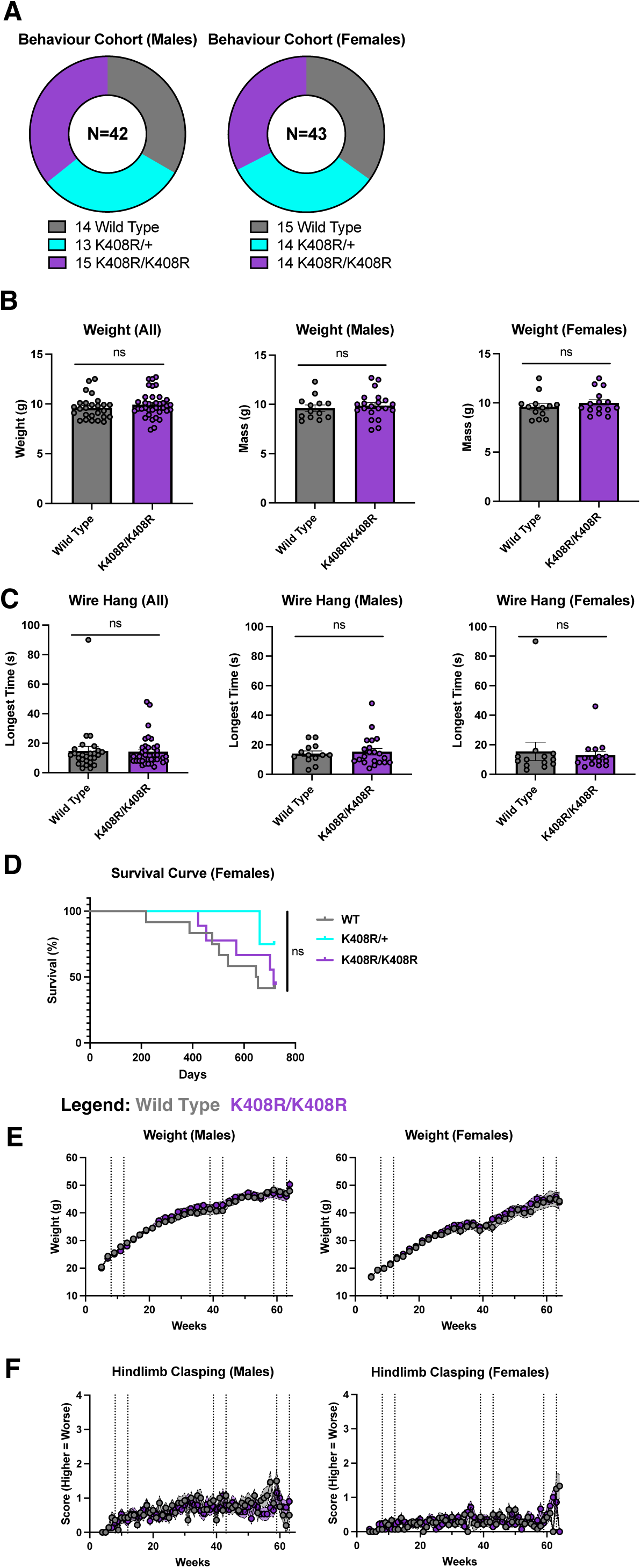
General wellness of TDP-43^K408R^ mice during development and aging. **(A)** Genotype and sex composition of mice subject to behaviour testing. **(B)** Weight of mice at P21. Unpaired T-test (Males); Mann-Whitney test (Females and All) **(C)** Quantification of the duration hanging on a wire at P21. Mann-Whitney test (Males, Females, and All). **(D)** Survival curve for female TDP-43^K408R^ mice (Males found in Fig. 5d). Curve comparisons analyzed using Log-Rank test and Gehan-Breslow-Wilcoxon test. **(E)** Weights of male and female TDP-43^K408R^ mice from 3 weeks to 64 weeks of age. Mixed-effects analysis. **(F)** Hindlimb clasping scores of male and female TDP-43^K408R^ mice from 3 weeks to 64 weeks of age. Mixed-effects analysis. See table S2 for raw data details on statistical tests. Data presented as mean ± SEM

**Fig. S8:**
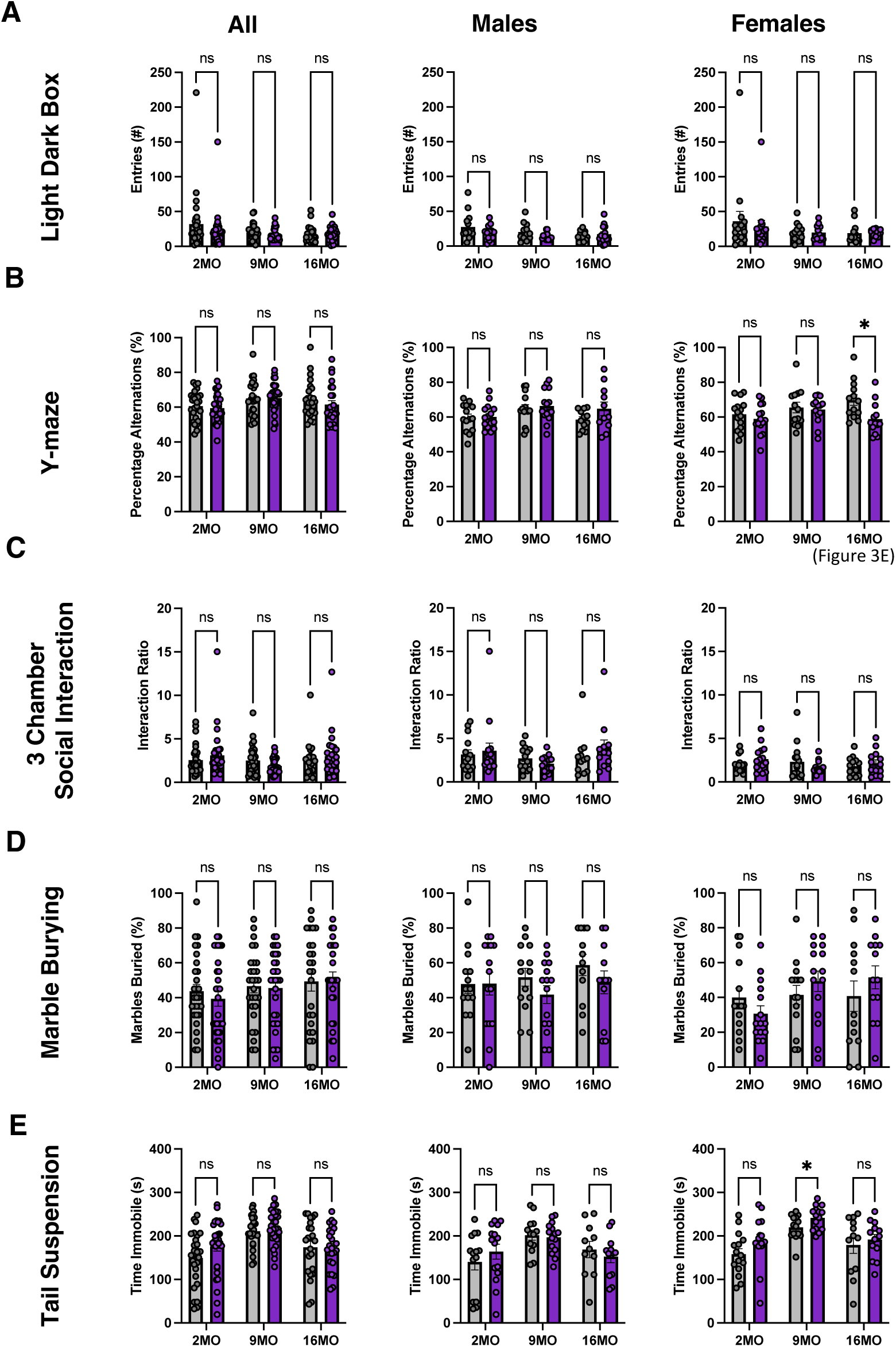
Characterizing the cognitive and social behaviour of TDP-43^K408R^ mice. Quantification of cognitive and social behaviours in the **(A)** Light/Dark Box, **(B)** Y-maze, **(C)** 3 Chamber Social Interaction, **(D)** Marble Burying, and **(E)** Tail Suspension behaviour tasks at 2, 9, and 16 months of age (2MO, 9MO, and 16MO, respectively) in male and female TDP-43^K408R^ mice. All behaviour tests were analyzed with a Mixed-Effects Model with Tukey’s multiple comparison analysis, see table S2. Data presented as mean ± SEM.

**Fig. S9:**
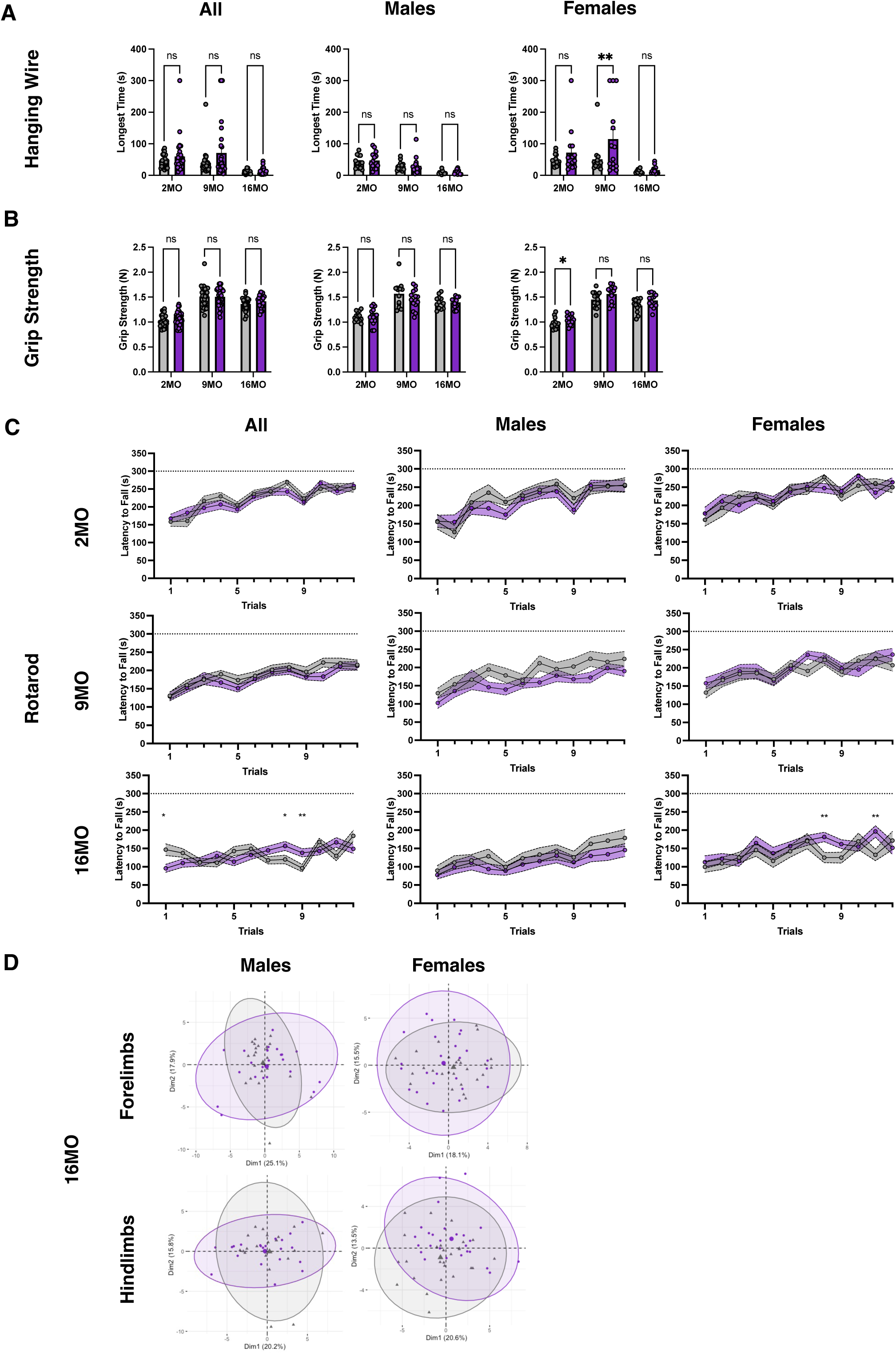
Characterizing the motor behaviour of TDP-43^K408R^ mice. Quantification of motor abilities in the **(A)** Hanging Wire, **(B)** Grip Strength, and **(C)** Rotarod behaviour tasks at 2, 9, and 16 months of age (2MO, 9MO, and 16MO, respectively) in male and female TDP-43^K408R^ mice. **(D)** Digigait behaviour tasks at 16 months of age in male and female TDP-43^K408R^ mice. Behaviour tests were analyzed with 2-Way ANOVA or Mixed-Effects Models, see table S2. Data presented as mean ± SEM.

**Fig. S10:**
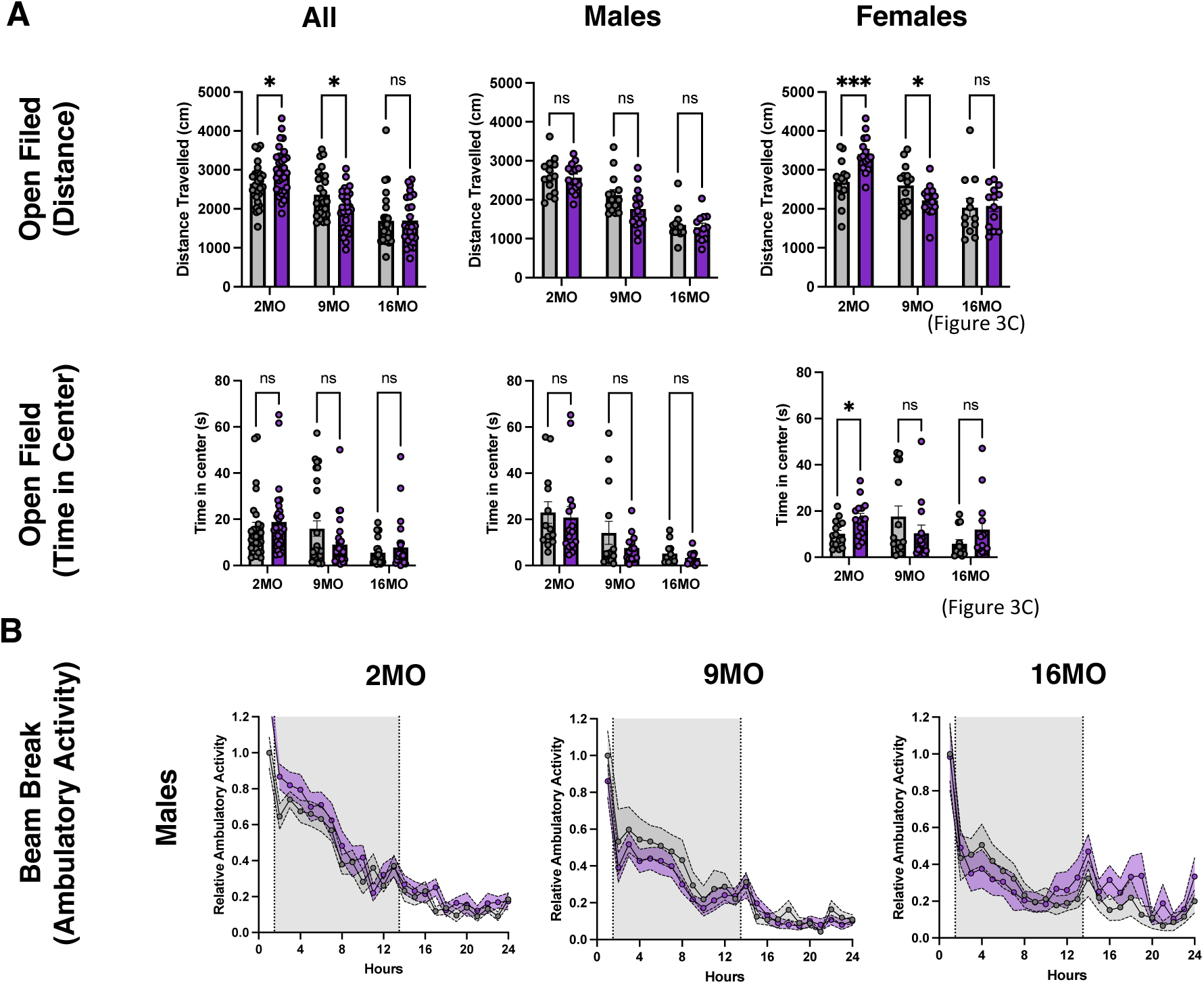
Characterizing the activity of TDP-43^K408R^ mice. Quantification of **(A)** total distance traveled (cm) and time in center (s) during the Open Field behaviour test and **(B)** the relative ambulatory activity (males) of TDP-43^K408R^ mice at 2, 9, and 16 months of age (2MO, 9MO, and 16MO, respectively). Behaviour tests were analyzed with 2-Way ANOVA or Mixed-Effects models, see table S2. Data presented as mean ± SEM.

**Fig. S11.**
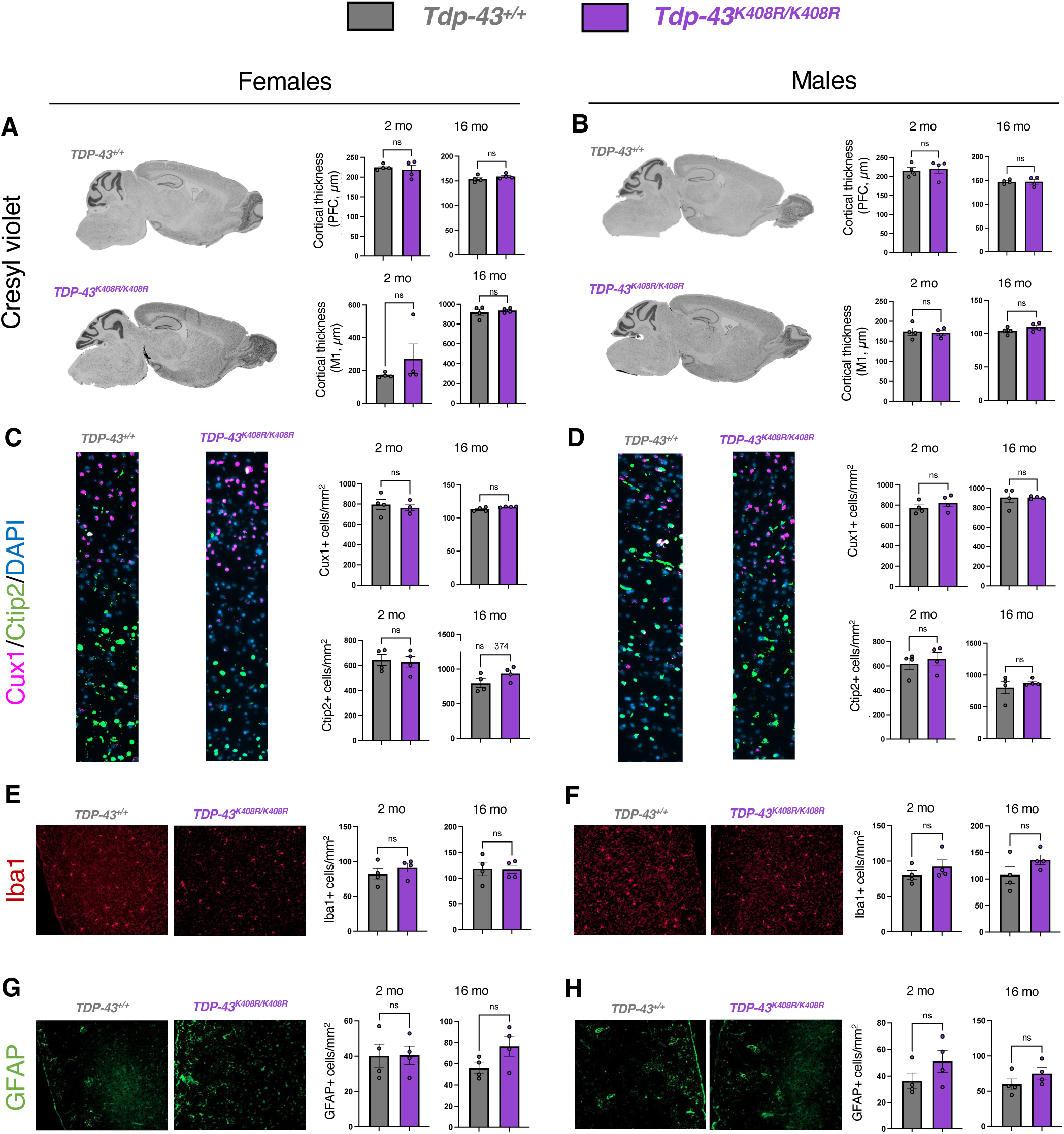
Absence of overt neurodegenerative or neuroinflammatory phenotypes in TDP-43^K408R/K408R^ mice. Cresyl violet staining used to measure cortical thickness in the prefrontal cortex (PFC) and motor area 1 (M1) of female **(A)** and male **(B)** TDP-43^K408R/K408R^ mice and littermate controls. Cux1 and Ctip2 staining and quantification in the motor cortex of female **(C)** and male **(D)** TDP-43^K408R/K408R^ mice and littermate controls. Iba1 staining of microglia **(E, F)** and GFAP staining of astrocytes **(G, H)** in female **(E,G)** and male **(F,H)** TDP-43^K408R/K408R^ mice and littermate controls. Quantification represents evaluation of N=4 mice per genotype, per sex, per time point. Micrographs are representative of the 16-month time point. Students T-test. Data presented as mean ± SEM. ns denotes p>0.05.

**Fig. S12:**
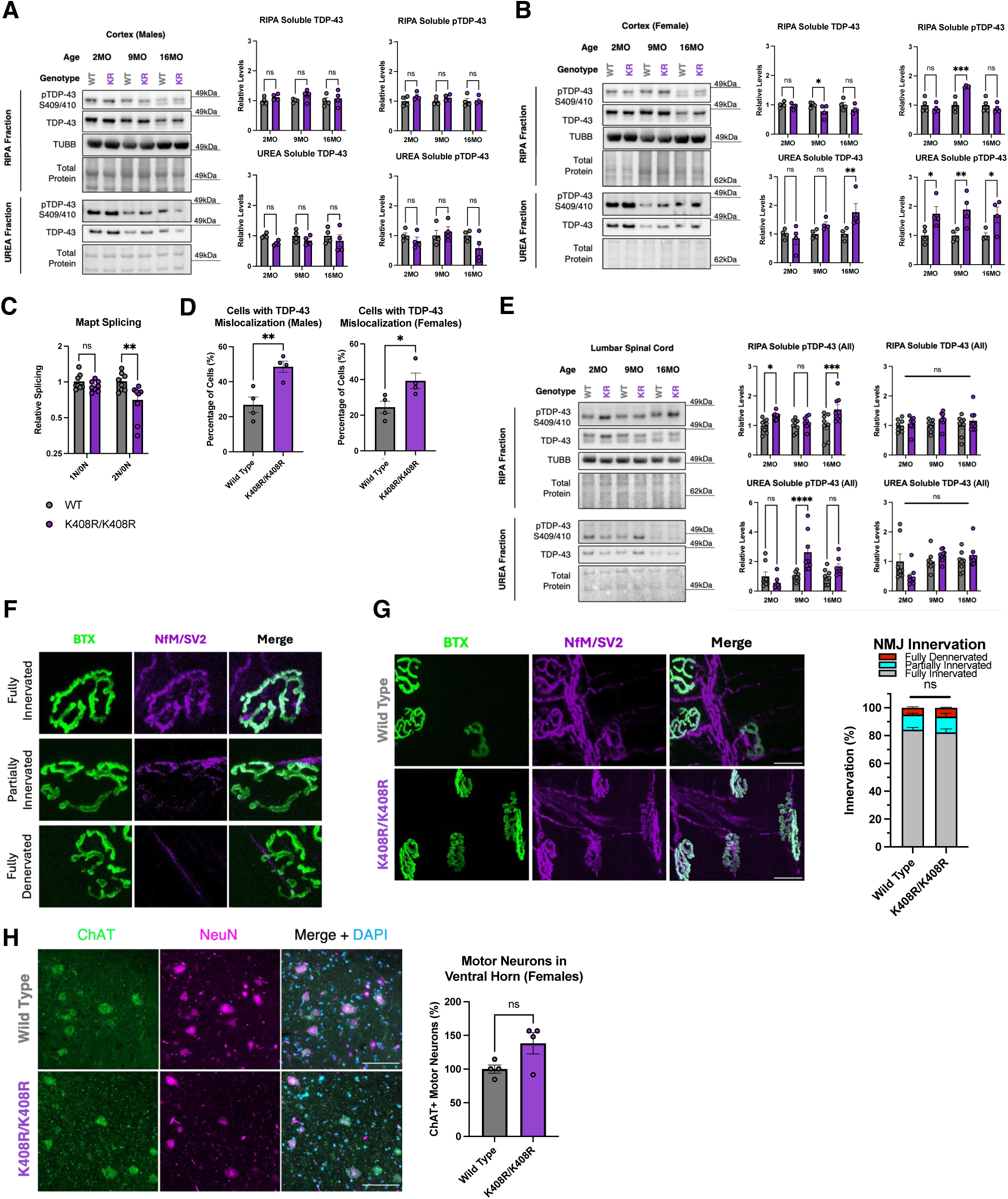
Characterizing the molecular and histological affects of blocking TDP-43 SUMOylation in vivo. **(A)** Representative western blot and quantification of TDP-43 and phosphorylated TDP-43 in the RIPA and UREA fractions of the cortex from male mice at 2, 9, and 16 months of age (2MO, 9MO, and 16MO, respectively). Data presented as mean ± SEM. (N = 4 per genotype) 2-Way ANOVA with Fisher’s LSD test. **(B)** Representative western blot and quantification of TDP-43 and phosphorylated TDP-43 in the RIPA and UREA fractions of the cortex from male mice at 2, 9, and 16 months of age (2MO, 9MO, and 16MO, respectively). Data presented as mean ± SEM. (N = 4 per genotype) 2-Way ANOVA with Fisher’s LSD test. *p<0.05, **p<0.005, ***p<0.0005. **(C)** Quantitative RT-PCR from 16 month old mouse cortex for splicing isoforms of *Mapt*. (N=8, 4 per sex, per genotype) 2-Way ANOVA with Fisher’s LSD Test. Data presented as mean ratios of ΔCt normalized to wild type ± SEM. **p<0.005. **(D)** Quantification of TDP-43 mislocalization in the lumbar spinal cord of 9-month-old from Fig. 5a split by sex. Each datapoint represents the average of 4 serial sections 40 μm apart from an individual mouse. Unpaired t-test. * = p<0.05, ** = p<0.005. **(E)** Representative western blot and quantification of TDP-43 and phosphorylated TDP-43 in the RIPA and UREA fractions of the lumbar spinal cord at 2, 9, and 16 months of age (2MO, 9MO, and 16MO, respectively). Data presented as mean ± SEM. (N = 8 per genotype; 4 per sex/genotype) 2-Way ANOVA with Fisher’s LSD test, *p<0.05, ***p<0.0005, ****p<0.0001 **(F)** Example images of neuromuscular junctions (NMJs) for scoring criteria to assess fully innervated, partially innervated, and fully denervated NMJs. **(G)** Representative images and quantification of neuromuscular junction (NMJ) innervation in the tibialis anterior of female 16-month-old TDP-43^K408R^ mice. >80 NMJs were quantified per animal. Data presented as mean ± SEM. (N = 4 per genotype) 2-Way ANOVA with Tukey’s multiple comparison analysis, *p<0.05, ****p<0.0001. (H) Representative images and quantification of ChAT+ motor neurons in the ventral horn of the lumbar spinal cord of male and female mice. Each datapoint is the average of 4 serial sections spaced 40 μm apart through the lumbar enlargement of the lumbar spinal cord. Data presented as mean ± SEM. Unpaired t-test, ** p<0.005.

**Fig. S13:**
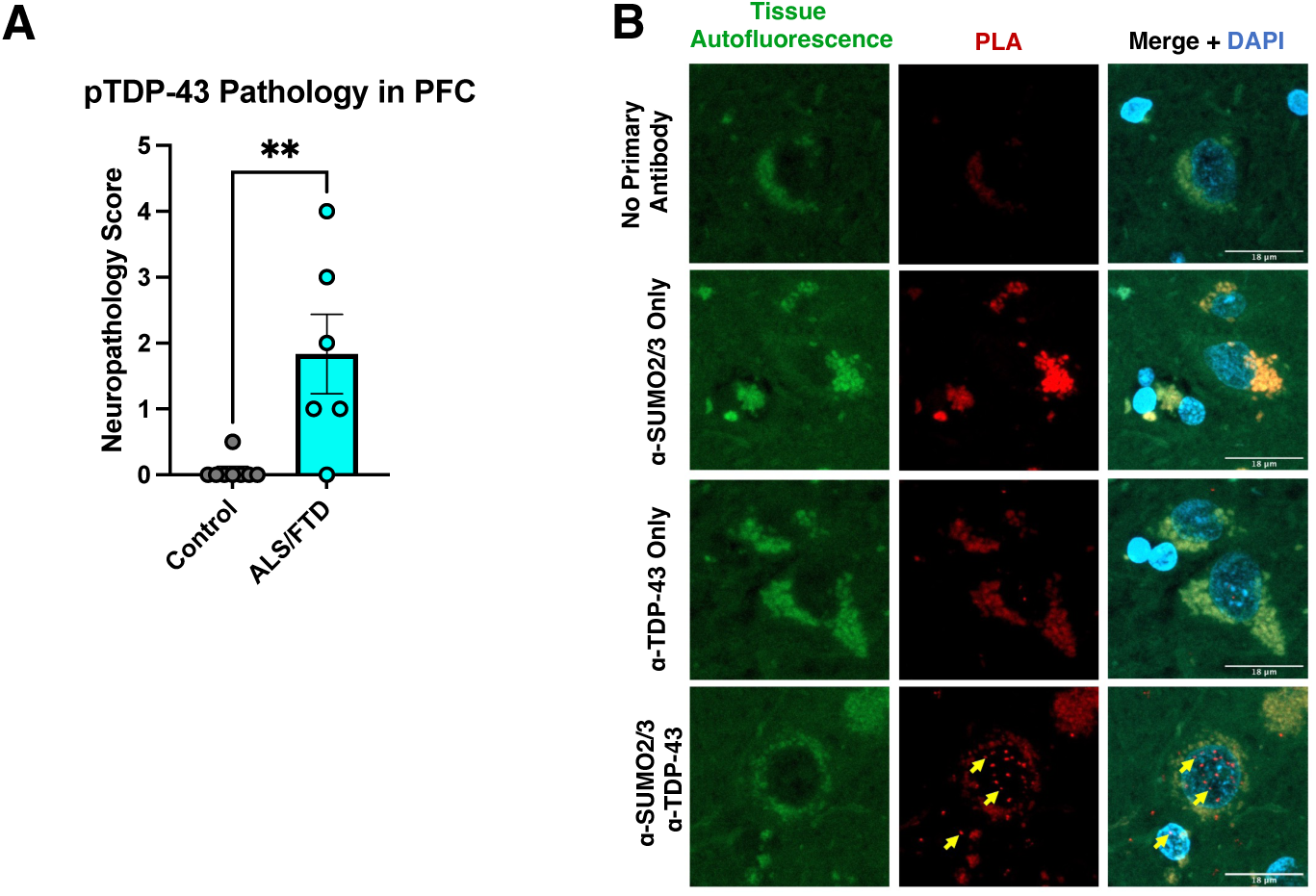
Optimization of proximity ligation assay in human tissue. **(A)** Quantification of neuropathology score of PPFE prefrontal cortex (PFC) tissue sections (5um) from patients diagnosed with ALS/FTD and unaffected controls. Data presented as mean ± SEM. (N = 6-8) Unpaired t-test, ** p<0.005. **(B)** Representative images of proximity ligation assay (PLA) between TDP-43 and SUMO2/3 in human PPFE hippocampus tissue sections (5um). Yellow arrows indicate TDP-43:SUMO2/3 interactions where the signal specific to the PLA channel and not identified in the autofluorescence channel. Scalebar = 18 μm.

## Supplementary Information

Supplementary Tables 1-4

Supplementary Video 1

## Materials and Methods

### Materials availability

Plasmids generated in this study will be deposited to Addgene by the time of publication. Mouse lines generated in this study are available upon request.

### Data and code availability

Human tissue data reported in this paper will be shared by the lead contact upon request.

Any additional information required to reanalyze the data reported in this paper is available from the lead contact upon request.

### Cloning of pEGFP-TDP-43 plasmid and variants

For expression of C-terminally tagged TDP-43-EGFP, human TDP-43 was subcloned out of a wtTDP-43tdTOMATOHA (gift from Zoushang Xu, Addgene # 28205) via XhoI and Kpn1 double digest and inserted into a pEGFP-N3 (Clontech) backbone and validated by sanger sequencing. Site directed mutagenesis was performed using QuikChange II XL (Agilent 200521) following manufacturer instructions. Primers for site directed mutagenesis can be found in table S4. Several TDP-43 variants were generated through the uOttawa Genome Engineering and Molecular Biology facility or designed and ordered through twist bioscience and were subcloned into the pEGFP-TDP-43 backbone for consistency using XhoI and HindIII double digest, see table S4 for plasmids used/generated in this study.

### Cell culture

HEK293T cells were cultured in DMEM outgrowth media containing 10% FBS (unless otherwise stated) and antibiotic/antimycotics at 37°C with 5% CO_2_.

### Generation of pL302-HA-SUMO2 stable HEK293T cell line

To generate a pL302-HA-SUMO2 lentivirus-compatible vector for stable HA-SUMO2 expression, HA-SUMO2 was subcloned from pcDNA-HA-SUMO2 (gift from Guy Salvesen, Addgene #48967) and inserted into the pL302 lentiviral backbone (gift from Jacqueline Burré and Thomas Südhof at Stanford). Briefly, extension PCR was performed to add an Xba1 restriction site to the 5’ and 3’ ends of the HA-SUMO2 insert using PCR amplification using forward primer sequence*: 5’-ggctgcaggtcgactctagaaagcttatggatgcctaccc-3’*, and reverse primer sequence*: 5’-tggctgcaggtcgactctagatgcatgctcgagtcaacct-3’.* The PCR amplicon and pL302 backbone were digested with XbaI and the amplicon was ligated into the pL302 backbone. Colonies were screened by sanger sequencing to ensure proper orientation and copy number of the insert. Next the pL302-HA-SUMO2 plasmid was packaged into lentivirus by co-transfection of psPAX2 (Gift from Didier Trono, Addgene #12259), and pMD2-G (Gift from Didier Trono, Addgene #12260) into HEK293T cells and subsequent media collections. Viral media was filtered through a 0.45 μm filter then concentrated by centrifugation at 100,000G for 2 hours at 4 °C and the pellet was resuspended in 1X PBS. Fresh HEK293T cells were transduced with pL302-HA-SUMO2 virus to generate a stable polyclonal cell line.

### GFP-trap SUMOylation assay

∼250,000 HEK293T and/or pL302-HA-SUMO2 HEK293T cells were seeded (experiment dependent) in a 6-well plate. The following day, plasmids were transfected to express TDP-43-GFP (or mutant variants) and/or HA-SUMO variants and allowed to incubate for 48 hours for optimal expression. Cells were treated with 250 μM NaAsO_2_ for one hour (or variations of time, concentration, or chemical depending on assay), after which cells were immediately collected by scraping the bottom of the well and pelleting the samples by centrifugation at 2500G for 5 minutes at room temperature. The supernatant was then aspirated, and the cells were flash frozen or immediately lysed in 450 uL of ice cold 1X Denaturing Lysis buffer (1X RIPA, supplemented with 1X protease inhibitor, 1X phosphatase inhibitor, 50mM N-Ethylmalamide, and 5% 2-mercaptoethanol) and boiled at 95 °C for 5 minutes. After boiling, 450 uL of ice cold 1X Lysis buffer (1X RIPA, supplemented with 1X protease inhibitor, 1X phosphatase inhibitor, 50mM N-Ethylmalamide) was then added to the sample, and cells were left on ice for 20 minutes with vortexing every 5 minutes for 10 second. Next, the cell lysates are centrifuged at ∼21000G for 20 minutes at 4 °C and during this time, 30 uL of GFP-Trap beads (Bulldog Bio) per sample were washed 3 times in cold 1X RIPA buffer then aliquoted equally into microcentrifuge tubes for each sample. Following the centrifugation, 1-3% of the supernatant was transferred into a microcentrifuge tube to be used as the *Input* sample. The remainder of the supernatant was transferred into a microcentrifuge tube containing the GFP-Trap beads as the immunoprecipitation sample and was then placed on a rotator for 45 minutes at 4 °C. After the incubation, the solution was aspirated, and the beads were washed using cold 1X RIPA buffer 5 times. Samples were eluted in 30 μL of 2X Laemmli buffer containing 10% Beta-Mercaptoethanol and boiled at 85 °C for 10 minutes at 1250 rpm on a thermomixer. The input samples were prepared with 4X Laemmli buffer containing 10% Beta-Mercaptoethanol and then boiled at 85 °C for 10 minutes. Samples were then analyzed by Western Blot. Quantification of SUMOylation assays was performed through densitometry analysis using ImageLab (BioRad) to quantify the volume of the ∼95kDa anti-HA bands from the immunoprecipitation standardized to the anti-GFP pulldown. For quantification of changes in TDP-43 SUMOylation response, conditions were normalized to the 1 hour sodium arsenite treatment for wild type TDP-43.

### SUMOylation assay

∼500,000 HEK293T and/or pL302-HA-SUMO2 HEK293T cells were seeded (experiment dependent) in a 6-well plate. The following day, cells were treated respective of their experiment after which cells were immediately collected by scraping the bottom of the well and pelleting the samples by centrifugation at 2500G for 5 minutes at room temperature. The supernatant was then aspirated, and the cells were flash frozen or immediately lysed in 100 uL of 1% SDS buffer (1% SDS in IP Lysis Buffer supplemented with 1X protease inhibitor, 1X phosphatase inhibitor, 100mM N-Ethylmalamide), vortexed for 20 seconds then boiled at 95C for 10 minutes. After boiling, 900 μL of ice-cold IP Lysis buffer was then added to the sample, and cells were left on ice for 20 minutes with vortexing every 5 minutes for 10 second. Next, the cell lysates are centrifuged at ∼21000G for 20 minutes at 4 C. Following the centrifugation, 1-3% of the supernatant was transferred into a microcentrifuge tube to be used as the *Input* sample. The remainder of the supernatant was transferred into a microcentrifuge tube containing 1 to 10 ug of antibody per sample (See table S5 for concentrations) and incubated overnight rotating at 4 °C. The next day, 50 μL of Protein G Dynabeads (Thermo Fisher Scientific) per sample were washed 3X in ice cold IP Lysis Buffer. The beads were resuspended in 50 μL IP Lysis Buffer per sample which was then added to each of the samples incubated with antibody. The samples were left to rotate at 4 °C for 45 minutes. Next, the solution was added to a magnetic rack kept at 4 °C and the supernatant was collected for *flowthrough* to test immunodepletion or aspirated. The beads were washed 5X in ice cold IP Lysis buffer. Samples were eluted using 5 ug of synthetic HA peptide (Sino Biological) per 1ug of antibody and incubating at 37 °C for 15 minutes at 900rpm on a thermomixer (for anti-HA immunoprecipitations). The input/flowthrough and HA immunoprecipitation samples were prepared with 4X Laemmli buffer containing 10% Beta-Mercaptoethanol and then boiled at 85 °C for 10 minutes. Samples were then analyzed by Western Blot using light chain specific secondary antibodies. Quantification of SUMOylation assays was performed through densitometry analysis using ImageLab (BioRad) to quantify the volume of the ∼65kDa anti-TDP-43 bands from the immunoprecipitation standardized to the loading control of the input. For quantification of changes in TDP-43 SUMOylation response, conditions were normalized to the 1 hour sodium arsenite treatment.

### Serial protein extraction from cell culture

Soluble protein was extracted using RIPA buffer (supplemented with protease inhibitor, phosphatase inhibitor, and 50mM N-Ethylmalamide) during immunoprecipitation assays (taken as *Input* prior to immunoprecipitation) or from lysate in cortical neuron experiments. The remaining pellet after the centrifugation step post-lysis was washed in 1mL RIPA buffer with vortexing for 10 seconds followed by centrifugation at ∼21000G for 20 minutes at 4 °C. The supernatant was carefully removed, and the pellet was resuspended in 300 μL of 2% SDS buffer in PBS for immunoprecipitation assays, or in 300 μL of 8M UREA in PBS with 10mM Tris-HCl pH 7.4. Samples were left to solubilize overnight at room temperature before 2X Laemmli was added prior to Western Blot analysis.

### Western blot analysis

Protein sample prepared in Laemmli loading buffer was loaded onto 8% polyacrylamide gel, TGx Mini-PROTEAN 4-15% precast gel (BioRad), or Bolt Bis-Tris Plus mini-Protein Gel 4-12% (Invitrogen) and run at 100-140 constant voltage in Tris-Glycine or MES buffer for their respective gels. Proteins were transferred onto a 0.45 μm nitrocellulose membrane at a constant 340 mA for 2 hours at 4 °C. The membranes were blocked in 10% milk diluted in TBS-T, washed 5 x 5 minutes in TBS-T then incubated in primary antibody overnight (table S5). The following day the membranes were washed for 5 x 5 minutes in TBS-T, then incubated in secondary antibody for 1-2 hours at room temperature. Finally, the secondary antibody is washed 5 x 5 minutes in TBS-T before being imaged using chemiluminescence Clarity Western ECL or Clarity Max Western ECL on an LAS4000 (GE). Densitometry was performed using the volumes function of the ImageLab (BioRad) software.

### Immunofluorescent microscopy analysis in HEK293T cells

Micro Coverglass #1.5 coverslips (Electron Microscopy Sciences) coverslips were washed in 2M HCl overnight at 55 °C, washed 5 times in sterile H_2_O, then pre-coated with 300 μL poly-D-lysine (10 μg/mL) overnight at 37 °C, then washed with distilled water three times and air-dried at room temperature for at least 2 hours or stored at 4 °C until required. 400 000 HEK293T cells were seeded in a 6 well dish and incubated overnight. 1000 ng of plasmid for protein expression was transfected into cells using Lipofectamine 3000 protocol (Thermo Fisher Scientific). Media was changed 4 hours post transfection to reduce toxicity. The following day transfected HEK293T cells were split onto coverslips plating ∼50 000 cells per coverslip. Cells were allowed to adhere for 24 hours. Cells were fixed using 10% buffered formalin for 10 minutes, then permeabilized in blocking buffer (10% serum, 1% Triton X-100, in 1X PBS) for 1 hour, then primary antibody diluted in blocking buffer was applied overnight (See supplemental table 5 for antibody concentrations). The following day coverslips were washed 5X in 1X PBS for 5 minutes then secondary antibody diluted in blocking buffer was applied for 2 hours at room temperature. Samples were stained with 1X DAPI (and/or 1:10000 dilution of CellMask Membrane Stain) in PBS for 10 minutes. Samples were washed 4 more times in PBS for 5 minutes each. Coverslips were briefly air dried and then mounted on slides using Vectashield Antifade Mounting Medium with DAPI. Z-stack images were obtained on a Zeiss AxioObserverZ1 LSM800 Confocal Microscope at 40× magnification with a 5× digital zoom or 63x magnification with 2X digital zoom through a Z distance of 10-12 μm per image using optimal spacing per slice with dimensions set to 1024 x 1024 pixels with 4X averaging per frame. At least 10 cells were imaged per replicate. Images were analyzed and quantified using ImageJ.

### E3 SUMO ligase screen

One day prior to transfection 62,000 HEK293T or HEK293T stably expressing HA-SUMO2 were plated in a 6 well dish. Samples were co-transfected using 800 ng of pCLIP-Dual-sgRNA for each of the SUMO E3 ligases alongside 1200ng pLenti-Cas9-BLAST and media was changed after 4 hours to minimize toxicity. Forty-eight hours after transfection, cells were selected using 2 μg/mL puromycin for 2-3 days. After selection was complete, cells were recovered in outgrowth media. On day 7 post transfection, 6-well plates were coated with 10 μg/mL poly-D-lysine for 1 hour at 37 °C then washed 3X in 1X PBS. 200,000 cells were plated into a 6 well dish for each knockout condition and respective controls. The following day, 500ng of pEGFP-N3 or pEGFP-TDP-43 was transfected for each respective condition and media was changed after 4 hours. Cells were incubated for 48 hours (10 days post Cas9/sgRNA transfection). Cells were then stressed for 1 hour using 250 μM sodium arsenite and GFP-Trap SUMOylation Assay and Western Blot analysis was performed as previously described. Samples were quantified using densitometry normalizing each condition to the stressed positive control condition on each respective gel.

### AlphaFold3 TDP-43 structure prediction

TDP-43 structure was predicted using the AlphaFold3 server (https://alphafoldserver.com/) using FASTA sequence of Human TDP-43 (Uniprot TADBP_HUMAN, Q13148) with relevant mutations. UG x6 (5’-UGUGUGUGUGUG-3’) RNA sequence was included for structure predictions.[121]

### Conservation analysis of TDP-43 K408

Representative FASTA amino acid sequences of TDP-43 paralogs were aligned for phylogenetic analysis using MUSCLE multiple alignment and PhyML maximum likeliness for phylogenetic tree generation[122]. Pairwise alignment of the C-terminal domain was performed using Geneious Prime 2022.1.1 (https://geneious.com) performing a MUSCLE multiple alignment of 300 FASTA amino acid sequences of TDP-43 paralogs between Humans and Actinopterygii representing the emergence of [human] K408 curated from the OrthoDB database (table S1).

### *TDP-43^K408R^* mouse line generation

At The Centre of Phenogenomics (SickKids, Toronto, ON, Canada), the *TDP-43^K408R^* mice were generated on a C57BL6/N background based on previous methods[123] using CRISPR/Cas9-mediated gene editing of the endogenous *Tardbp* locus. In brief, spCas9 loaded with an sgRNA to target exon 6 of *Tardbp (MGI:2387629; 5’-TGGGGGCTTTGGCTCGAGCA-3’)* was electroporated into embryos alongside a single stranded oligonucleotide (ssODN) repair template (5’-C*TAAATCTACCTAACCTAATAACCAACCTACTAACCACCCCCCACCACCTACATTCCCCAG CCAGAAGAC**c**TAGAATCCATG**ga**CGAGCCAAAGCCCCCATTAAAACCACTGCCCGATCCTG CATTTGATGCTGACCCCCAACCAAGGGGGGC-3’*). Embryos were screened via allelic discrimination (see “TDP-43^K408R^ genotyping” below) to identify founders. Four founders were crossed to C57BL6/N mice to ensure germline transmission of knock-in allele.

### Mouse husbandry

All mouse procedures were carried out in accordance with the Canadian Council on Animal Care and approved by the University of Ottawa Animal Care Committee. All mice were group housed (3-5 per mice cage) with access to food and water *ad libitum* on a standard 12-hour light dark cycle. Exceptions included adult male mice who were separated and single housed if persistent fighting and fight wounds were observed in the cage. All experimental mice were given crinkle paper in addition to the standard nestlet and hut enrichment material. Husbandry was completed by University of Ottawa Animal Care and Veterinary Services except for cohorts actively undergoing behavior testing for which husbandry was completed by the experimenter.

### Mouse wellness monitoring

Mice were monitored weekly starting at age P21 and included assessment of hindlimb clasping, kyphosis, weight, and general health. The hindlimb function was assessed following previous methods [124]. Briefly, the mouse was suspended in the air for approximately 5-10 seconds by the base of the tail. Features of the limbs were assessed to give a score from 0-4. Kyphosis was assessed by allowing the mice to briefly walk on the flat table top in the housing room and visually observing the straightness of the spine. A score of 0-3 was given based on previously described methods[125]. Weight was measured every other week by placing individual mice on a digital weigh scale. Notably, mice exhibiting barbered patches were monitored and recorded.

### Sex and age of experimental mice

Experiments were performed using both male and female mice sufficiently powered to detect sex differences. In some instances where no sex differences were observed, male and female mice were grouped together to assess genotypic differences independent of sex. Experiments involving primary cortical neurons cultures dissected from individual embryos were used involving a mixture of male and female embryos. Experimental animals underwent behavioral testing as early as P21 and as late as 16 months of age. See Table S2 for number of animals used (distinguished by sex and genotype), raw behavior data, and specific statistical analyses for each behavior experiment.

### TDP-43^K408R^ genotyping

Tail samples were collected prior to weaning and again postmortem for genomic DNA (gDNA) isolation and genotyping. Tails were solubilized in 300 μL solubilization buffer (10X SET, 100 mM NaCl, 100 μg/mL Proteinase K (Bio Basic PB0451-250)) at 55 °C overnight. Cell debris was precipitated by adding 150 μL of “Tail Salts Buffer” (4.31M NaCl, 0.63M HCl, 10 mM Tris-HCl, pH7.4) then samples were centrifuged at ∼21,000*g* for 15 minutes at 4 °C. Supernatant containing gDNA was transferred into 600 μL of chilled 100% ethanol to precipitate nucleic acids which were then pelleted by centrifugation for 10 minutes at ∼21,000*g* at 4 °C. The supernatant was carefully removed, and the pellet was washed in 600 μL of chilled 70% ethanol then centrifuged at ∼21,000*g* at 4 °C for 5 minutes. The supernatant was carefully removed, and residual supernatant was left to evaporate for 5 minutes. The gDNA pellet was resuspended in double distilled H2O at 55 °C for 10 minutes. Genotyping reaction was prepared in a 10 μL reaction containing ∼5ng of gDNA, 250nM *K408R Genotyping Forward* primer (5’-*CCACCATTCTAAATCTACCTAACCTAATA-3’*), 250nM *K408R Genotyping Reverse* primer (5’-*GGATCGGGCAGTGGTTTTA-3’*), 125nM wild type Locked Nucleic Acid (LNA) probe (*HEX-TCT+A+A+GT+CT+TCT+GGC-IowaBlack FQ),* 125nM K408R LNA probe (*FAM-TCT+A+G+GT+CT+T+CT-IowaBlack FQ),* and 2X PerfeCTa qPCR Tough Mix (QuantaBio). Reactions were run on a BioRad CFX96 qPCR thermocycler with the following cycling parameters: Initial annealing at 95 °C for 2 minutes, followed by 40 cycles of 95 °C for 15 seconds then 60 °C for 60 seconds. Results were analyzed via allelic discrimination in the BioRad Maestro software. See Fig. S2b.

### Primary cortical neuron cultures

Pregnant mice were euthanized between gestation E13.5–15.5 with 120 mg/kg Pentobarbital Sodium (Bimeda-MTC, 8015E) delivered via intraperitoneal injection. The embryos were removed and placed into ice cold 1X PBS. For each embryo, the cortices were carefully isolated and meninges removed and then placed in ice cold HBSS (Sigma Aldrich). Cortices were dissociated for 20 minutes with trypsin (Thermo Scientific) at room temperature on a rotator before trypsin inhibitor with DNase solution was added to quench the reaction. Cells were pelleted at 2,500 x *g* for 5 minutes at 4°C. The supernatant was carefully removed and the pellet was washed with trypsin inhibitor plus DNase solution. Cortical neurons were pelleted at 2,500 x *g* for 5 minutes at 4°C. The supernatant was carefully removed and the neurons were resuspended in 1 mL of ice cold outgrowth media (Neurobasal media (Thermo Scientific), supplemented with 1X B-27 (Thermo Scientific), 1X N-2 (Thermo Scientific), 500 μM L-Glutamine (Wisent Bioproducts), and 0.5% penicillin/streptomycin (GE Healthcare Life Sciences)). Culture dishes and coverslips were prepared in advanced coated with 50 μg/mL Poly-D-Lysine overnight at 37 °C specific to each experiment detailed below. Cultures were maintained for 7-9 days *in vitro* prior (DIV) to experimentation.

### Murine primary neuron culture maintenance

Primary cortical neurons were dissected and maintained in Neurobasal media (Thermo Scientific), supplemented with 1X B-27 (Thermo Scientific), 1X N-2 (Thermo Scientific), 500 μM L-Glutamine (Wisent Bioproducts), and 0.5% penicillin/streptomycin (GE Healthcare Life Sciences) at 37 °C with 5% CO_2_. Each neuron culture was harvested from a single embryo representing one biological replicate. The sex of the embryos/cultures were not determined thus experiments contain a mix of both male and female replicates.

### Endogenous SUMOylation assay from ex vivo embryonic mouse brains

Mouse embryos were harvested following *Cortical Primary Neuron Culture* methods described above. Brains were dissected from embryos and stored in ice cold HBSS (Sigma Aldrich). Brains were cut using a razor blade down the midline evenly to separate hemibrains and each hemibrain was transferred into a 1.5 mL microcentrifuge tube containing neurobasal complete media used in cortical *Primary Cortical Neuron Culture Maintenance* described above. One hemibrain was subjected to 42 °C heat shock for 1 hour while the complementary hemibrain was maintained at 37 °C for 1 hour as a control. Samples were briefly centrifuged at 2000g for 2 minutes to pellet hemibrain, then supernatant was carefully removed. The hemibrain pellets were immediately lysed in 100 μL of 1% SDS buffer (1% SDS in IP Lysis Buffer supplemented with 1X protease inhibitor, 1X phosphatase inhibitor, 100mM N-Ethylmalamide), and homogenized with a pestle then boiled at 95 °C for 10 minutes. After boiling, 900 μL of ice-cold IP Lysis buffer was then added to the sample then samples were vortexed aggressively for 10 seconds and passed through a 18G insulin needle. Lysates were left on ice for 20 minutes with vortexing every 5 minutes for 10 second to ensure complete lysis. Next, the lysates were centrifuged at ∼21000G for 20 minutes at 4 °C. Following the centrifugation, 1-3% of the supernatant was transferred into a microcentrifuge tube to be used as the *Input* sample. The remainder of the supernatants were equally divided (∼450 uL each) into two separate 1.5 mL microcentrifuge tubes containing 10 μg of normal mouse IgG (Alpha Diagonstic 20008-250) or 10 ug of anti-SUMO2/3 clone [8A2] (Abcam ab8137) per sample. Samples were incubated overnight rotating at 4 °C. The next day, 50 μL of Protein G Dynabeads (Thermo Fisher Scientific) per sample were washed 3X in ice cold IP Lysis Buffer. The beads were resuspended in 50 μL IP Lysis Buffer per sample which was then added to each of the samples incubated with antibody. The samples were left to rotate at 4 °C for 45 minutes. The solution was added to a magnetic rack kept at 4 °C and the supernatant was collected for *flowthrough* to test immunodepletion or aspirated. The beads were washed 5X in ice cold IP Lysis buffer. Samples were eluted using 30uL (at 1mg/mL) of synthetic SUMO2/3 peptide (Amino Acid Sequence: *IRFRFDGQPI,* synthesized through Genscript) per sample and incubating at 37 °C for 15 minutes at 900rpm on a thermomixer. The input/flowthrough and immunoprecipitation samples were prepared with 4X Laemmli buffer containing 10% Beta-Mercaptoethanol and then boiled at 85 °C for 10 minutes. Samples were then analyzed by Western Blot using light chain specific secondary antibodies. Quantification of SUMOylation assays was performed through densitometry analysis using ImageLab (BioRad) to quantify the volume of the ∼62kDa anti-TDP-43 bands from the immunoprecipitation standardized to the loading control of the input. For quantification of changes in TDP-43 SUMOylation response, conditions were normalized to the 1 hour sodium arsenite treatment.

### Immunofluorescence in primary cortical neurons

Micro Coverglass #1.5 coverslips (Electron Microscopy Sciences) coverslips were washed in 2M HCl overnight at 55 °C, washed 5 times in sterile H_2_O, then pre-coated with poly-D-lysine (50 μg/mL) overnight at 37 °C, then washed with distilled water three times and air-dried at room temperature for at least 2 hours. Primary mouse cortical neurons were seeded at 75,000-100,000 cells per coverslip and cultured as described above. On day 7, neurons were fixed for 10 minutes using 10% phosphate buffered formalin for (Fisher Chemical, SF100-4) followed by 3 x 5-minutes washes in 1 mL of 1X PBS. Neurons were blocked in 500 μL of blocking buffer (1% Triton X-100, 10% cosmic calf serum in 1X PBS) for 1 hour, then incubated in 300 μL of primary antibody diluted in blocking buffer overnight at 4 °C. The following day, the neurons were washed for 5 x 5-minutes in 1 mL of 1X PBS then incubated in 300 μL of secondary antibody diluted in blocking buffer for 2 hours at room temperature. Neurons were then washed in 1 mL 1X PBS, then stained with DAPI for 10 minutes at room temperature followed by 4 x 5-minutes washes in 1X PBS before being mounted using antifade fluorescence mounting media (Dako, S3023). Z-stack images were obtained on a Zeiss AxioObserverZ1 LSM800 Confocal Microscope at 40× magnification with a 2× digital zoom through a Z distance of 10-12 μm per image using optimal spacing per slice with dimensions set to 1024 x 1024 pixels with 4X averaging per frame. At least 50 cells were imaged and quantified per replicate. For TDP-43 foci colocalization, Z-stack images were obtained on a Zeiss AxioObserver Z1 LSM880 AiryScan2 confocal microscope at 63x magnification with 3X digital zoom at 512 x 512 resolution. At least 10 cells presenting with TDP-43 foci were imaged per replicate. Images were analyzed and quantified using ImageJ.

### TDP-43 colocalization with HSPA1L

HSPA1L colocalization experiment was designed similar to previous approaches that identified TDP-43 forming anisosomes in response to stress[96]. HSPA1L-mRuby2 was designed and synthesized in a pTwist-Lenti-Puro backbone (Twist Bioscience). Next the pTwist-Lenti-HSPA1L-mRuby2 plasmid was packaged into lentivirus by co-transfection of psPAX2 (Gift from Didier Trono, Addgene #12259), and pMD2-G (Gift from Didier Trono, Addgene #12260) at equimolar concentrations into HEK293T cells. After 24 hours, media was changed and discarded. Media was collected 48 hour and 72 hours post transfection. Viral media was filtered through a 0.45 μm filter then concentrated by centrifugation at 100,000g for 2 hours at 4 °C and the pellet was resuspended in 1X PBS. Cortical primary neurons were cultured as previously described and lentiviral infection was performed at the time of plating (0 DIV). Cells were cultured to 7 DIV then were stressed with 250 μM sodium arsenite for 1 hour and recovered for 3 hours to induce TDP-43 foci formation. Cells were fixed in 10% formalin for 10 minutes and immunoflurescent staining was performed for TDP-43 as described above. Z-stack images were obtained on a Zeiss AxioObserver Z1 LSM880 AiryScan2 confocal microscope at 63x magnification with 3X digital zoom at 512 x 512 resolution. At least 10 cells presenting with TDP-43 foci and HSPA1L foci were imaged per replicate. Images were analyzed and quantified using ImageJ.

### Proximity ligation assay in primary cortical neurons

Primary mouse cortical neurons were cultured and fixed as described in *primary cortical neuron cultures* and *immunofluorescence in primary cortical neurons*. Fixed coverslips were transferred into 12-well plates and outlined with a hydrophobic pen and blocked using 40 μL of Duolink blocking buffer (Sigma Aldrich, DUO82007) at 37 °C for 1 hour. Coverslips were then washed for 3 × 5 minutes in 1 mL 1X PBS. Next, coverslips were incubated in 300 μL of primary antibody (anti-TDP-43 1:750, PTGLabs; anti-SUMO2/3 [8A2] 1:500 Sigma) diluted in blocking buffer (1.5% Triton X-100, 10% cosmic calf serum in 1X PBS) overnight at 4 °C. The next day, the coverslips were washed for 3 × 5 minutes in 1 mL of Duolink Wash Buffer A (0.01 M Tris-Base, 0.15 M NaCl, 0.05% Tween-20, pH 7.4) followed by incubation at 37°C for 1 hour in 40 μL of dilute probe mixture containing a 1:5 dilution of Duolink PLA MINUS (Sigma Aldrich, DUO82004) and PLUS probes (Sigma Aldrich, DUO82002) in antibody diluent (Sigma Aldrich, DUO82008). Duolink PLA probes were diluted in antibody diluent (Sigma Aldrich, DUO82008) at a 1:5 dilution. Coverslips were washed for 3 × 5 minutes in Duolink Wash Buffer A and then incubated in 40 μL of ligase (Sigma Aldrich, DUO82027) diluted in 1X ligation buffer (Sigma Aldrich, DUO82009) at a 1:40 dilution at 37 °C for 30 minutes. The coverslips were then washed for 3 x 5-minutes washes in Duolink Wash Buffer A and then incubated in 40 μL of polymerase (Sigma Aldrich, DUO82028) diluted 1:80 in 1X amplification buffer (Sigma Aldrich, DUO82011) at 37°C for 90 minutes. Next, the coverslips were washed 2 × 10 minutes in Duolink Wash Buffer B (0.2 M Tris-Base, 0.1 M NaCl, pH 7.5) and then again in 1 mL of Duolink Wash Buffer B diluted at 1:100 in 1X PBS for 1 minute. Coverslips were briefly air dried and then mounted on slides using Vectashield Antifade Mounting Medium with DAPI. Z-stack images were obtained on a Zeiss AxioObserverZ1 LSM800 Confocal Microscope at 63× magnification with a 5× digital zoom through a Z distance of 10-12 μm per image using optimal 0.27 μm spacing per slice with dimensions set to 512 × 512 pixels with 2X averaging per frame. ∼5 images were randomly obtained around the coverslip sampling >15 cells per condition per replicate. Images were analyzed and quantified using the Spots function on the Imaris (ver. 9.9.1 Bitplane, Switzerland) software.

### Behavior testing general information

All adult behavior testing was performed in the University of Ottawa Behaviour and Physiology Core. During behavior testing periods the mice were minimally disturbed: food and water were only added/changed as needed and cages were changed by the experimenter with a transfer of old bedding material. Apart from nesting and Beam Break testing, all behavior tests were performed in the light phase of the cycle. The experimenter was blinded to the genotype of the mice during behavior testing and until the mice were end pointed. Mice were brought to the testing room to habituate in dim white light at least 30 minutes prior to commencing testing with the exceptions of nest building and fear conditioning for which there was no habituation period. The behavior cohort consisted of males and females with 12-15 animals per sex per genotype. These mice had been backcrossed three times to C57BL/6N background. All mice used for histology, biochemistry, and RNA analyses performed the fear conditioning test 1 week prior to dissections to control for potential molecular changes induced by this behavior test.

### Developmental testing

A small battery of behavior tests was completed at P8 and P21. At P8, these tested included (I) weight, (II) righting reflex, (III) hindlimb suspension, and (IV) forelimb suspension. (I) Weight was measured every other week by placing individual mice on a digital weigh scale. (II) Mice were placed on their back on a tabletop and timed for how long it took them to right themselves onto their paws. (III) Mice were placed facing downward in a 50mL Falcon tube (VWR 21008-940) and timed for their latency to fall. (IV) Mice were placed to grasp a wire suspended by a pencil holder and timed for latency to fall. At P21, hanging wire was performed in which the mouse was allowed to grasp a wire 30 cm off the ground and time for its latency to fall. Each mouse completed three trials with a maximum time of 60 seconds.

### Beam break

The Beam Break test was used to assess general locomotor activity and habituation to a novel cage environment. The apparatuses used to record and analyze the mice activity included the Micromax analyzer/Fusion Software (Omnitech Electronics; Columbus, OH, USA) at the 2-month time point and the Photobeam Activity System (San Diego Instruments; San Diego, CA, USA) at the 9 and 16-month timepoints. Clean cages with only a thin layer of corncob bedding, food, and water were loaded into the recording frame. Mice were singly housed in one of these cages 2 hours prior to the start of their dark cycle. For a 24-hour period at the standard 12-hour dark/light cycle, the activity of the mice was monitored by infrared beam emitters and receptors. When the test was completed, the sum of the infrared beam breaks in 5-minute, 1-hour, and 24-hour sampling bins was analyzed.

### Digigait

The Digigait treadmill Imager and Analysis Software (Mouse Specifics Inc.) were used to record and analyze parameters of gait, respectively. Mice were placed on the unmoving treadmill surface and once recording was started the speed was increased to 18 cm/s with 0 degree incline. A 3 second video of each mouse walking continuously, with no stopping or starting, was captured. A mouse was excluded from the test if it was unable to walk for 3 seconds at the 18 cm/s speed. Digiait data was analyzed using FactoMinR [126] and visualized using FactoExtra [127] packages in R (version 4.2.3).

### Fear conditioning

Contextual and cued fear conditioning were tested using a 3-day protocol to assess associative fear learning and memory. Naïve age and sex matched mice were initially used to determine the optimal shock value. On day 1 (training), the mice were placed into a Phenotyper box (Noldus Information Technology) with a grid shock floor (Med Associates). The testing room was set to 60 lux light level and 70 dB of white noise (context A). The mice were left in the apparatus for 6 minutes during which they receive 3 tone-shock pairings (30 seconds tone co-terminated with a 2 second foot shock). On day 2 (Context), the mice were placed in the same apparatus (Context A) with no tone or foot shock delivered for the 6-minute trial. On day 3 (cue), the animals were placed in the test apparatus with an altered context (context B) including red light, no white noise, vanilla scent, textured mat covering the shock floor and plastic inserts in the apparatus. The mice are allowed to explore this context with no tone for 3 minutes, and then are presented with the same tone from Day 1 for the last 3 minutes. The time freezing was analyzed with EthoVision software for all 3 testing days.

### Grip strength

A grip strength meter (Chatillon DFE II, Columbus Instruments) was used to assess the maximal forelimb grip strength of the mice. The grip strength meter was rotated vertically and temporarily mounted on a flat surface prior to testing. The mouse was brought near the triangular attachment and allowed to grasp the lower bar with its forepaws. The mouse was pulled directly downward, by the tail, in one smooth motion. Each mouse was tested, and the grip strength recorded for 5 consecutive trials with a 1-minute intertrial interval. If a trial was deemed unsuccessful by the blinded experimenter, the trial was redone. Causes of an unsuccessful trial included the mouse prematurely losing its grip on the bar or grasping the bar with its hindfoot. The average grip strength for the 5 successful trials was analyzed.

### Hanging wire

The hanging wire test was used to assess muscle strength and coordination. A metal wire was secured to the top of a tall plastic box with padding on the bottom. The mouse was brought near and allowed to grasp the wire with its forepaws. A timer was started once the experimenter released the mouse to allow it to hang freely. When the mouse fell from the wire the time was recorded. Mice had three trials and were allowed to hang for a maximum of 600 seconds with a 60 second intertrial interval in between. If a mouse hung for 600 seconds, it did not complete any additional trials. The maximum hanging time from the trials was used for analysis.

### Light/dark box

The light dark paradigm was used to assess anxiety-like or disinhibited, exploratory behavior. The testing apparatus (Med Associates) has two equal sized rectangular compartments the mouse can move freely between. The one is fully illuminated whereas the other is covered by a black plastic insert. The position of the mouse in the apparatus testing field is recorded using infrared beams. To start the test, the mice are placed into the lit side and allowed to explore for 10 minutes. The time spent in each compartment and number of entries into each compartment was recorded and analyzed by Activity Monitor software (Med Associates).

### Marble burying

The marble burying assay can assess the motor function required to bury objects as well as cognitive changes related to apathy or perseverance. Cages were filled with 10cm of fresh corncob bedding and one mouse was placed to habituate for 5 minutes. After habituation, 20 glass marbles were laid out evenly in a 4 by 5 pattern. The mouse was returned to the cage and left alone for 30 minutes in 60 lux light. With the completion of the trial, the number of marbles buried by at least two thirds was scored by a blinded experimenter.

### Nest building

Nest building is an innate behavior of mice in their daily lives. This complex behavior requires both executive planning and sensorimotor coordination. Directly following Beam Break testing, a single square nestlet (5 cm^2^ cotton pad) was placed in each Beam Break cage for 16 – 18 hours, of which 12 of these hours was during the dark cycle. At the end of the test, images were taken of the nests and scored blindly for quality as previously described on a scale from 1-5.

### Open field

The open field test was used to assess anxiety and locomotor activity in a novel environment. The apparatus consists of white plastic square arenas measuring 45cm on each side. The mice were placed in a corner of the arena and allowed to freely explore for 10 minutes with light levels at 300 lux. The distance travelled, time spent in the corners and center of the field was recorded and analyzed by EthoVision software (Noldus Information Technology).

### Rotarod

The rotarod (IITC Life Science, Ugo Basile) was used to test the motor performance of mice including coordination and resistance to fatigue. Mice were placed in the stationary rotarod bar for 10 seconds before the rotarod program was initiated. The bar accelerated from 4 rpm to 40 rpm for 5 minutes and the latency to fall for each mouse was recorded. The time was stopped when the mouse fell from the bar or rotated passively. Mice were completed four trials per day, with a 10-minute intertrial interval in their home cage, for 3 consecutive days.

### Spontaneous Y-maze

The spontaneous Y-Maze test was used to assess spatial working memory. The Y-shaped maze has three identical arms at 120°C around a center point triangle. The mice were placed in the center point and allowed to freely explore the arms for 8 minutes. The movement of the mouse was tracked including the sequence of arm entries by EthoVision (Noldus Information Technology). An alternation is defined as the mouse making consecutive, sequential entries into each of the three arms without revisiting an arm. The alternation index was calculated as (number of alternations/(total number of arm entries minus two)) and reported as a percent.

### Tail suspension

The automated Tail Suspension apparatus (Med Associates) was used to assess apathetic-like behaviors of the mice. Mice were taped and suspended by the tail to a vertical steel bar which measures strain gauge as the mouse moves during a 6-minute trial. The cumulative time spent immobile, hanging passively (below lower threshold) was measured by the Tail Suspension software.

### Three chamber sociability

The Three Chamber test was conducted to measure the sociability of mice when allowed to interact with a novel mouse or a similarly sized inanimate object. The testing apparatus is a 19 x 45 cm plastic box divided into three equal chambers with clear plastic wall dividers. The two external chambers each have a single weighted metallic mesh pencil holder and the central chamber is empty. The mice are habituated to the apparatus for 5 minutes by being placed in the central chamber and allowed to freely explore and enter all chambers. The mouse has a 5-minute intertrial interval in its home cage. In the test trial, a sex and age matched wild type mouse (social target) is placed beneath one mesh pencil holder, and an inanimate plastic toy (non-social target) is placed beneath the other. To commence the test trial, the mouse is paced in the central chamber. The time spent in each chamber and interacting with the social or non-social target is recorded and analyzed by EthoVision software (Noldus Information Technology).

### Tube test of social dominance

Tube test was used to assess social dominance and within cage social hierarchy. Cage mates were paired against each other for testing in a round-robin design. The tube test was conducted on a flat tabletop, which the mice were allowed to run around on for 5 minutes prior to commencing the testing. One foot of vinyl tubing was used. Mice were habituated to the tube prior to testing on the same day by encouraging them to run through the empty tube from either side 5 times. A blinded experimenter placed a mouse on either end of the tube, and released their tales when they completely entered the tube. The first mouse to step its hind paws out of the tube lost the battle. The battle was redone if after 2 minutes no mouse had won.

### Collection and preparation of mouse tissue

Mice were harvested for both histology and biochemistry/molecular analyses to reduce total animal numbers. Mice were first acclimated for 2 hours prior to dissection to help control for activity-induced changes and all dissections were performed at zeitgeber hours ZT6 to ZT11 to control for potential circadian affects. Mice were randomly dissected by cage to limit batch effects and additional stress. Mice were anesthetized using isoflurane inhalation, blood was collected, and then mice were sacrificed via decapitation and the brains were quickly isolated, weighted, then cut into 2 hemispheres where the left hemisphere was drop fixed in 10% buffered formalin and the right hemisphere was dissected by region and flash frozen on dry ice. Spinal cords were removed using hydraulic extrusion and the lumbar spinal cord was isolated. The lumbar enlargement was isolated for histology whereas the remaining lumbar spinal cord was used for molecular analysis. Tibialis anterior and soleus muscles were isolated and drop fixed in 10% buffered formalin for imaging studies and gastrocnemius muscle was flash frozen.

### Preparation of mouse tissue for analysis by histology

Tissue was collected as described in *Collection and preparation of mouse tissue*. Following fixation for 48 hours, samples were transferred to a 70% ethanol solution and sent to the Louise Pelletier Histology Core facility at the University of Ottawa for paraffin embedding and microtome sectioning. The samples were sectioned at a thickness of 5 µm. Serial sectioning was performed at four levels within the tissue each separated by 40 µm. For staining, slide-mounted sections were deparaffinized in 100% xylenes (Fisher Scientific X3P-1GAL) for 10 minutes, then transferred into a second container of fresh 100% xylenes for an additional 10 minutes. The slides were then rehydrated in descending ethanol solutions: two 5 minutes treatments in 100% ethanol, 5 minutes in 70% ethanol, and 5 minutes in 50% ethanol. The slides were then immersed in 1X PBS for at least 5 minutes to rinse residual ethanol.

### Cresyl violet staining

*Slides containing PPFE mounted sections were slowly dipped 20 times in 70% ethanol followed by 95% ethanol, 100% ethanol, 95% ethanol, and 70% ethanol. Slides were then submerged in ddH2O for 1 minute. Next the sections were incubated in 0.25% Cresyl Violet stain for 2 minutes. Samples were then dipped 10 times in ddH2O followed by 70% ethanol, 95% ethanol, 0.25% glacial acetic acid, 95% ethanol, and finally 100% ethanol. Permount was applied to sections and mounted with coverslips*.

### Immunofluorescence analysis in mouse tissue

Antigen was performed by placing slides in 1X sodium citrate buffer (2.94g sodium citrate 0.5mL Tween-20 in 1L 1X PBS, pH6) at 95°C for 30 minutes then rinsed in 1X PBS. Tissue sections were then blocked in blocking buffer containing 1% Triton-X and serum (5% horse serum, 5% cosmic calf serum, or 1% bovine serum albumin) in 1X PBS for 2 hours at room temperature. Next the tissue sections were incubated with primary antibody (table S5) overnight at 4°C. The next day, the sections were washed twice in PBS 1X + 0.1% Triton-X for 5 minutes each and then 3X in 1X PBS for 5 minutes each. The slides were then incubated in secondary antibody at room temperature for 2 hours. The slides were then washed in 1X PBS + 0.1% Triton-X for 5 x 5-minutes, before mountin with #1.5 coverslips (Thermo Fisher Scientific 12-544E) and fluorescent mounting media (Dako S3023). The coverslips were sealed with clear nail polish and allowed to dry before imaging. Images of each ventral horn were taken on a Zeiss Axio Imager 2 at 20X magnification. Cells were counted manually per image using Fiji (ImageJ).

### Neuromuscular junction analysis

Tissue was collected as described in *Collection and preparation of mouse tissue*. After 24 hours of fixation, tibialis anterior muscles were stored in a solution of 1X PBS. Muscle bundles were carefully teased apart from the tissue into 4-5 bundles with forceps. The muscle bundles were transferred to 6-well plate cell culture plate containing 1% Triton-X in 1X PBS overnight at 4°C under gentile agitation for permeabilization. The next day, the muscle bundles were washed three times in 1X PBS for 5 minutes at room temperature. They were placed in a blocking buffer solution of 4% bovine serum albumin and 1% Triton-X in 1X PBS at 4°C overnight. The following day, the muscles were incubated in primary antibody (BTX and NfL/SV2 cocktail, see table S5) diluted in blocking buffer solution overnight at 4°C. The next day, the muscle bundles were washed with 1X PBS three times for 5 minutes and incubated with secondary antibody diluted in blocking buffer solution at room temperature for 2 hours. The muscle bundles were then washed 3 times in 1X PBS for 5 minutes. The bundles were then placed on slides, mounted with VECTASHIELD® HardSet™ Antifade Mounting Medium (#H-1400), and #1.5 coverslips placed above the muscle bundles. The slides were left at room temperature overnight in the dark to allow the mounting media to cure. Images were taken on a Zeiss LSM800 Confocal Microscope at 20X through a Z-stack with optimal spacing (0.61 µm). Images were taken throughout the entirety of the muscle tissue such that at least 80 neuromuscular junctions per animal were captures. Images were scored manually per image using Fiji (ImageJ). See Fig. S8E for scoring examples.

### SUMOylation analysis in human brain tissue

20 mg of frozen human brain tissue was homogenized in 300 μL of RIPA buffer supplemented with protease inhibitor, phosphatase inhibitor, and 50mM N-Ethylmalamide and vortexed every 5 minutes for 20 minutes while incubating on ice. The samples were centrifuged at 21000G for 20 minutes at 4°C and the supernatant (RIPA soluble fraction) was carefully removed. 4X laemmli buffer was added and samples were boiled for 5 minutes at 95 °C then stored overnight at 4 °C. The remaining pellet after the centrifugation step post-lysis was washed in 1mL RIPA buffer with vortexing for 10 seconds followed by centrifugation at ∼21000G for 20 minutes at 4 °C. The supernatant was carefully removed, and the pellet was resuspended in 300 μL of of 8M UREA in PBS with 10mM Tris-HCl pH 7.4. Samples were vortexed and left to solubilize overnight at room temperature before 4X Laemmli was added prior to Western Blot analysis. RIPA and UREA samples were randomly loaded onto gels (separated by fraction) with 4 control samples consistently loaded (aged 20-30) on each gel for normalization. Western blot was performed as described above. Samples were standardized to ponceau signal then normalized to control samples.

### Collection and preparation of FFPE human brain tissue

Frontal cortex tissues were collected antemortem from three sporadic ALS/FTLD patients (a 74-year-old male, 67-year-old female and 59-year-old female) obtained through the ALS Clinic at Sunnybrook Health Sciences Centre, Toronto. ALS was diagnosed using the revised El Escorial Criteria (Brooks et al., 2000) and informed consent was obtained with approval from the local ethical review board. The presence of TDP-43 proteinopathy within the frontal cortex was verified through immunohistochemical labeling using rat anti-phosphorylated TDP-43 (p409/410) antibody on formalin fixed paraffin embedded sections. Genetic analyses confirmed absence of mutations in key ALS/FTD-associated genes: *C9orf72*, *SOD1*, *FUS* and *TARDBP*.

### Neuropathological assessment of all human PFC tissues from various sites

*Human FFPE PFC samples collected from the NIH Neurobiobank, CHEO, and University of Toronto were independently assessed and scored by a neuropathologist to verify TDP-43 pathology. Immunohistochemistry staining was performed on FFPE tissue sections using the Leica bond system. Sections were pre-treated using sodium citrate buffer (pH 7.0, epitope retrieval solution 1) for 20 minutes and then incubated using a 1:2000 dilution of anti-phosphorylated TDP-43 S409/410 (Cosmo Bio Co. Ltd, #TIP-PTD-P07) for 40 minutes at room temperature and detected using an HRP conjugated compact polymer system. Slides were then stained using DAB as the chromogen, counterstained with Hematoxylin, mounted on slides and covered with coverslips. TDP-43 pathology was assessed by a score of 0-4 looking for neuronal cytoplasmic inclusions and glial cytoplasmic inclusions, particularly within layers II and V/VI. Scoring: 0 = negative, 1 = very rare inclusions, 2 = inclusions readily visible, 3= moderate density of inclusions, 4= highest density of inclusions*.

### Proximity ligation assay in human FFPE tissue

Formalin fixed paraffin-embedded human brain sections (5 μm sections) were deparaffinized using two 10-minute washes with xylenes, two 5-minute washes with 100% ethanol, one 5-minute wash with 70% ethanol, and one 5-minute was with 50% ethanol. Deparaffinized sections then underwent sodium citrate antigen retrieval (10mM sodium citrate, 0.05% Tween-20, pH 6) for 2 hours at 80°C. Sections were then blocked with Duolink blocking buffer (Sigma Aldrich, DUO82007) for 1 hour at room temperature. Next, sections were incubated with primary antibodies (anti-TDP-43 1:750, PTGLabs; anti-SUMO2/3 [8A2] 1:500 Sigma) diluted in blocking buffer (1.5% Triton X-100, 10% cosmic calf serum in 1X PBS) overnight at 4°C. The following day, the sections were washed in Duolink Wash Buffer A (0.01M Tris-Base, 0.15M NaCl, 0.05% Tween-20, pH 7.4), followed by incubation in Duolink PLA MINUS (Sigma Aldrich, DUO82004) and PLUS probes (Sigma Aldrich, DUO82002) at 37°C for 1 hour. Duolink PLA probes were diluted in antibody diluent (Sigma Aldrich, DUO82008) at a 1:5 dilution. Sections were washed in Duolink Wash Buffer A and then incubated in ligase (Sigma Aldrich, DUO82027) at 37°C for 30 minutes. Ligase was diluted in 1X ligation buffer (Sigma Aldrich, DUO82009) at a 1:40 dilution. Then, the sections were washed in Duolink Wash Buffer A and then incubated in polymerase (Sigma Aldrich, DUO82028) at 37°C for 90 minutes. Polymerase was diluted in 1X amplification buffer (Sigma Aldrich, DUO82011) at a 1:80 solution. The sections were then washed in Duolink Wash Buffer B (0.2M Tris-Base, 0.1M NaCl, pH 7.5) and then again in Duolink Wash Buffer B diluted at 1:1000. Finally, sections were incubated with DAPI at room temperature for 15 minutes. DAPI was diluted at 1:1000 in PBS. Fluorescent mounting media (Dako, S3023) and #1.5 coverslips were then placed over the sections and sealed using clear nail polish. Images were taken on a Zeiss LSM800 Confocal Microscope. 5 images per tissue were randomly collected from grey matter at 63X magnification and 2X digital zoom through a Z stack of 10-12 μm with optimal spacing (0.27 μm) at a resolution of 512 x 512 pixels. PLA foci were manually counted per image using Fiji (ImageJ).

### Statistical analyses

Statistical tests were performed using PRISM 10.2.2. Test type was selected based on the number of comparisons made, repeated measures, and gaussian distribution of the data. Levels of statistical significance are indicated in figure legends.

## References

1. Hipp MS, Kasturi P, Hartl FU (2019) The proteostasis network and its decline in ageing. Nature Reviews Molecular Cell Biology 2019 20:7 20:421–435

2. Spires-Jones TL, Attems J, Thal DR (2017) Interactions of pathological proteins in neurodegenerative diseases. Acta Neuropathologica 2017 134:2 134:187–205

3. Hardiman O, Al-Chalabi A, Chio A, Corr EM, Logroscino G, Robberecht W, Shaw PJ, Simmons Z, Van Den Berg LH (2017) Amyotrophic lateral sclerosis. Nature Reviews Disease Primers 2017 3:1 3:1–19

4. Grossman M, Seeley WW, Boxer AL, et al (2023) Frontotemporal lobar degeneration. Nature Reviews Disease Primers 2023 9:1 9:1–19

5. Burrell JR, Halliday GM, Kril JJ, Ittner LM, Götz J, Kiernan MC, Hodges JR (2016) The frontotemporal dementia-motor neuron disease continuum. The Lancet 388:919–931

6. Nelson PT, Dickson DW, Trojanowski JQ, et al (2019) Limbic-predominant age-related TDP-43 encephalopathy (LATE): consensus working group report. Brain 142:1503–1527

7. Meneses A, Koga S, O’Leary J, Dickson DW, Bu G, Zhao N (2021) TDP-43 Pathology in Alzheimer’s Disease. Molecular Neurodegeneration 2021 16:1 16:1–15

8. Nicks R, Clement NF, Alvarez VE, et al (2023) Repetitive head impacts and chronic traumatic encephalopathy are associated with TDP-43 inclusions and hippocampal sclerosis. Acta Neuropathol 145:395–408

9. Thammisetty SS, Pedragosa J, Weng YC, Calon F, Planas A, Kriz J (2018) Age-related deregulation of TDP-43 after stroke enhances NF-κB-mediated inflammation and neuronal damage. J Neuroinflammation 15:1–15

10. Coyne AN, Baskerville V, Zaepfel BL, Dickson DW, Rigo F, Bennett F, Patrick Lusk C, Rothstein JD (2021) Nuclear accumulation of CHMP7 initiates nuclear pore complex injury and subsequent TDP-43 dysfunction in sporadic and familial ALS. Sci Transl Med 13:1923

11. Suk TR, Rousseaux MWC (2020) The role of TDP-43 mislocalization in amyotrophic lateral sclerosis. Mol Neurodegener 15:45

12. Ma XR, Prudencio M, Koike Y, et al (2022) TDP-43 represses cryptic exon inclusion in the FTD–ALS gene UNC13A. Nature 2022 603:7899 603:124–130

13. Brown AL, Wilkins OG, Keuss MJ, et al (2022) TDP-43 loss and ALS-risk SNPs drive mis-splicing and depletion of UNC13A. Nature 2022 603:7899 603:131–137

14. Melamed Z, López-Erauskin J, Baughn MW, et al (2019) Premature polyadenylation-mediated loss of stathmin-2 is a hallmark of TDP-43-dependent neurodegeneration. Nat Neurosci 1

15. Klim JR, Williams LA, Limone F, et al (2019) ALS-implicated protein TDP-43 sustains levels of STMN2, a mediator of motor neuron growth and repair. Nat Neurosci 22:167–179

16. Gasset-Rosa F, Lu S, Yu H, Chen C, Melamed Z, Guo L, Shorter J, Da Cruz S, Cleveland DW (2019) Cytoplasmic TDP-43 De-mixing Independent of Stress Granules Drives Inhibition of Nuclear Import, Loss of Nuclear TDP-43, and Cell Death. Neuron 102:339–357.e7

17. Baughn MW, Melamed Z, López-Erauskin J, et al (2023) Mechanism of STMN2 cryptic splice-polyadenylation and its correction for TDP-43 proteinopathies. Science (1979) 379:1140–1149

18. Agra Almeida Quadros AR, Li Z, Wang X, et al (2024) Cryptic splicing of stathmin-2 and UNC13A mRNAs is a pathological hallmark of TDP-43-associated Alzheimer’s disease. Acta Neuropathologica 2024 147:1 147:1–18

19. Spence H, Waldron FM, Saleeb RS, et al (2024) RNA aptamer reveals nuclear TDP-43 pathology is an early aggregation event that coincides with STMN-2 cryptic splicing and precedes clinical manifestation in ALS. Acta Neuropathol 147:1–15

20. Hergesheimer RC, Chami AA, De Assis DR, Vourc’h P, Andres CR, Corcia P, Lanznaster D, Blasco H (2019) The debated toxic role of aggregated TDP-43 in amyotrophic lateral sclerosis: A resolution in sight? Brain 142:1176–1194

21. Zou ZY, Zhou ZR, Che CH, Liu CY, He RL, Huang HP (2017) Genetic epidemiology of amyotrophic lateral sclerosis: a systematic review and meta-analysis. J Neurol Neurosurg Psychiatry 88:540–549

22. Ghasemi M, Brown RH (2018) Genetics of Amyotrophic Lateral Sclerosis. Cold Spring Harb Perspect Med. 10.1101/CSHPERSPECT.A024125

23. Mackenzie IR, Nicholson AM, Sarkar M, et al (2017) TIA1 Mutations in Amyotrophic Lateral Sclerosis and Frontotemporal Dementia Promote Phase Separation and Alter Stress Granule Dynamics. Neuron 95:808–816.e9

24. Walker AK, Soo KY, Sundaramoorthy V, et al (2013) ALS-associated TDP-43 induces endoplasmic reticulum stress, which drives cytoplasmic TDP-43 accumulation and stress granule formation. PLoS One 8:e81170

25. Tam OH, Rozhkov N V., Shaw R, et al (2019) Postmortem Cortex Samples Identify Distinct Molecular Subtypes of ALS: Retrotransposon Activation, Oxidative Stress, and Activated Glia. Cell Rep 29:1164–1177.e5

26. Wolozin B, Ivanov P (2019) Stress granules and neurodegeneration. Nat Rev Neurosci 20:649–666

27. Lin MT, Beal MF (2006) Mitochondrial dysfunction and oxidative stress in neurodegenerative diseases. Nature 443:787–795

28. Ratti A, Gumina V, Lenzi P, et al (2020) Chronic stress induces formation of stress granules and pathological TDP-43 aggregates in human ALS fibroblasts and iPSC-motoneurons. Neurobiol Dis 145:105051

29. D’Amico E, Factor-Litvak P, Santella RM, Mitsumoto H (2013) Clinical perspective on oxidative stress in sporadic amyotrophic lateral sclerosis. Free Radic Biol Med 65:509–527

30. Colombrita C, Zennaro E, Fallini C, Weber M, Sommacal A, Buratti E, Silani V, Ratti A (2009) TDP-43 is recruited to stress granules in conditions of oxidative insult. J Neurochem 111:1051–1061

31. Zuo X, Zhou J, Li Y, et al (2021) TDP-43 aggregation induced by oxidative stress causes global mitochondrial imbalance in ALS. Nature Structural & Molecular Biology 2021 28:2 28:132–142

32. Ueda T, Takeuchi T, Fujikake N, et al (2024) Dysregulation of stress granule dynamics by DCTN1 deficiency exacerbates TDP-43 pathology in Drosophila models of ALS/FTD. Acta Neuropathol Commun 12:1–15

33. Iguchi Y, Katsuno M, Takagi S, Ishigaki S, Niwa J ichi, Hasegawa M, Tanaka F, Sobue G (2012) Oxidative stress induced by glutathione depletion reproduces pathological modifications of TDP-43 linked to TDP-43 proteinopathies. Neurobiol Dis 45:862–870

34. Liu-Yesucevitz L, Bilgutay A, Zhang YJ, et al (2010) Tar DNA binding protein-43 (TDP-43) associates with stress granules: Analysis of cultured cells and pathological brain tissue. PLoS One 5:e13250

35. McDonald KK, Aulas A, Destroismaisons L, Pickles S, Beleac E, Camu W, Rouleau GA, Velde C Vande (2011) TAR DNA-binding protein 43 (TDP-43) regulates stress granule dynamics via differential regulation of G3BP and TIA-1. Hum Mol Genet 20:1400–1410

36. Luan W, Wright AL, Brown-Wright H, Le S, San Gil R, Madrid San Martin L, Ling K, Jafar-Nejad P, Rigo F, Walker AK (2023) Early activation of cellular stress and death pathways caused by cytoplasmic TDP-43 in the rNLS8 mouse model of ALS and FTD. Molecular Psychiatry 2023 28:6 28:2445–2461

37. Fung G, Shi J, Deng H, Hou J, Wang C, Hong A, Zhang J, Jia W, Luo H (2015) Cytoplasmic translocation, aggregation, and cleavage of TDP-43 by enteroviral proteases modulate viral pathogenesis. Cell Death & Differentiation 2015 22:12 22:2087–2097

38. Ash PEA, Stanford EA, Al Abdulatif A, et al (2017) Dioxins and related environmental contaminants increase TDP-43 levels. Mol Neurodegener 12:1–14

39. Dubinski A, Gagne M, Peyrard S, Gordon D, Talbot K, Vande Velde C (2023) Stress granule assembly in vivo is deficient in the CNS of mutant TDP-43 ALS mice. Hum Mol Genet 32:319–332

40. Odierna GL, Vucic S, Dyer M, Dickson T, Woodhouse A, Blizzard C (2024) How do we get from hyperexcitability to excitotoxicity in amyotrophic lateral sclerosis? Brain 147:1610–1621

41. Zhang P, Fan B, Yang P, Temirov J, Messing J, Kim HJ, Taylor JP (2019) Chronic optogenetic induction of stress granules is cytotoxic and reveals the evolution of ALS-FTD pathology. Elife. 10.7554/eLife.39578

42. Neumann M, Kwong LK, Lee EB, et al (2009) Phosphorylation of S409/410 of TDP-43 is a consistent feature in all sporadic and familial forms of TDP-43 proteinopathies. Acta Neuropathol 117:137–149

43. Neumann M, Sampathu DM, Kwong LK, et al (2006) Ubiquitinated TDP-43 in Frontotemporal Lobar Degeneration and Amyotrophic Lateral Sclerosis. Science (1979) 314:130–133

44. Arai T, Hasegawa M, Akiyama H, et al (2006) TDP-43 is a component of ubiquitin-positive tau-negative inclusions in frontotemporal lobar degeneration and amyotrophic lateral sclerosis. Biochem Biophys Res Commun 351:602–611

45. Necarsulmer J, Simon J, Evangelista B, et al (2023) RNA-binding deficient TDP-43 drives cognitive decline in a mouse model of TDP-43 proteinopathy. Elife. 10.7554/ELIFE.85921.2

46. McGurk L, Gomes E, Guo L, Mojsilovic-Petrovic J, Tran V, Kalb RG, Shorter J, Bonini NM (2018) Poly(ADP-Ribose) Prevents Pathological Phase Separation of TDP-43 by Promoting Liquid Demixing and Stress Granule Localization. Mol Cell 71:703–717.e9

47. Henley JM, Craig TJ, Wilkinson KA (2014) Neuronal SUMOylation: mechanisms, physiology, and roles in neuronal dysfunction. Physiol Rev 94:1249–1285

48. Wilkinson KA, Henley JM (2010) Mechanisms, regulation and consequences of protein SUMOylation. Biochemical Journal 428:133–145

49. Maraschi AM, Gumina V, Dragotto J, Colombrita C, Mompeán M, Buratti E, Silani V, Feligioni M, Ratti A (2021) SUMOylation Regulates TDP-43 Splicing Activity and Nucleocytoplasmic Distribution. Mol Neurobiol 58:5682–5702

50. Maraschi A, Gumina V, Dragotto J, Colombrita C, Mompeán M, Buratti E, Silani V, Feligioni M, Ratti A (2021) SUMOylation Regulates TDP-43 Splicing Activity and Nucleocytoplasmic Distribution. Mol Neurobiol 58:5682–5702

51. Maurel C, Chami AA, Thépault RA, Marouillat S, Blasco H, Corcia P, Andres CR, Vourc’h P (2020) A role for SUMOylation in the Formation and Cellular Localization of TDP-43 Aggregates in Amyotrophic Lateral Sclerosis. Mol Neurobiol 57:1361–1373

52. Marino R, Buccarello L, Hassanzadeh K, Akhtari K, Palaniappan S, Corbo M, Feligioni M (2023) A novel cell-permeable peptide prevents protein SUMOylation and supports the mislocalization and aggregation of TDP-43. Neurobiol Dis 188:106342

53. Pichler A, Melchior F (2002) Ubiquitin-Related Modifier SUMO1 and Nucleocytoplasmic Transport. Traffic 3:381–387

54. Wang L, Wansleeben C, Zhao S, Miao P, Paschen W, Yang W (2014) SUMO 2 is essential while SUMO 3 is dispensable for mouse embryonic development. EMBO Rep 15:878–885

55. Enserink JM (2015) Sumo and the cellular stress response. Cell Div 10:1–13

56. Seyfried NT, Gozal YM, Dammer EB, Xia Q, Duong DM, Cheng D, Lah JJ, Levey AI, Peng J (2010) Multiplex SILAC analysis of a cellular TDP-43 proteinopathy model reveals protein inclusions associated with SUMOylation and diverse polyubiquitin chains. Molecular and Cellular Proteomics 9:705–718

57. Dammer EB, Fallini C, Gozal YM, et al (2012) Coaggregation of RNA-binding proteins in a model of TDP-43 proteinopathy with selective RGG motif methylation and a role for RRM1 ubiquitination. PLoS One. 10.1371/JOURNAL.PONE.0038658

58. Dewey CM, Cenik B, Sephton CF, Dries DR, Mayer P, Good SK, Johnson BA, Herz J, Yu G (2011) TDP-43 Is Directed to Stress Granules by Sorbitol, a Novel Physiological Osmotic and Oxidative Stressor. Mol Cell Biol 31:1098–1108

59. Chang HY, Hou SC, Way T Der, Wong CH, Wang IF (2013) Heat-shock protein dysregulation is associated with functional and pathological TDP-43 aggregation. Nat Commun 4:1–11

60. Suk TR, Nguyen TT, Fisk ZA, et al (2023) Characterizing the differential distribution and targets of Sumo1 and Sumo2 in the mouse brain. iScience 26:106350

61. Lu J, Wu T, Zhang B, Liu S, Song W, Qiao J, Ruan H (2021) Types of nuclear localization signals and mechanisms of protein import into the nucleus. Cell Communication and Signaling 19:1–10

62. Lange A, Mills RE, Devine SE, Corbett AH (2008) A PY-NLS Nuclear Targeting Signal Is Required for Nuclear Localization and Function of the Saccharomyces cerevisiae mRNA-binding Protein Hrp1. Journal of Biological Chemistry 283:12926–12934

63. Sun Y, Zhao K, Xia W, et al (2020) The nuclear localization sequence mediates hnRNPA1 amyloid fibril formation revealed by cryoEM structure. Nature Communications 2020 11:1 11:1–8

64. Mann JR, Gleixner AM, Mauna JC, et al (2019) RNA Binding Antagonizes Neurotoxic Phase Transitions of TDP-43. Neuron 102:321–338.e8

65. Keiten-Schmitz J, Wagner K, Piller T, Kaulich M, Alberti S, Müller S (2020) The Nuclear SUMO-Targeted Ubiquitin Quality Control Network Regulates the Dynamics of Cytoplasmic Stress Granules. Mol Cell 79:54–67.e7

66. Kagey MH, Melhuish TA, Wotton D (2003) The polycomb protein Pc2 is a SUMO E3. Cell 113:127–137

67. Garcáa-Gutiérrez P, Juárez-Vicente F, Gallardo-Chamizo F, Charnay P, García-Domínguez M (2011) The transcription factor Krox20 is an E3 ligase that sumoylates its Nab coregulators. EMBO Rep 12:1018–1023

68. Oh SM, Liu Z, Okada M, Jang SW, Liu X, Chan CB, Luo H, Ye K (2009) Ebp1 sumoylation, regulated by TLS/FUS E3 ligase, is required for its anti-proliferative activity. Oncogene 2010 29:7 29:1017–1030

69. Eisenhardt N, Chaugule VK, Koidl S, et al (2015) A new vertebrate SUMO enzyme family reveals insights into SUMO-chain assembly. Nature Structural & Molecular Biology 2015 22:12 22:959–967

70. Payne F, Colnaghi R, Rocha N, et al (2014) Hypomorphism in human NSMCE2 linked to primordial dwarfism and insulin resistance. J Clin Invest 124:4028–4038

71. Kahyo T, Nishida T, Yasuda H (2001) Involvement of PIAS1 in the Sumoylation of Tumor Suppressor p53. Mol Cell 8:713–718

72. Kotaja N, Karvonen U, Jänne OA, Palvimo JJ (2002) PIAS Proteins Modulate Transcription Factors by Functioning as SUMO-1 Ligases. Mol Cell Biol 22:5222– 5234

73. Nakagawa K, Yokosawa H (2002) PIAS3 induces SUMO-1 modification and transcriptional repression of IRF-1. FEBS Lett 530:204–208

74. Sachdev S, Bruhn L, Sieber H, Pichler A, Melchior F, Grosschedl R (2001) PIASy, a nuclear matrix–associated SUMO E3 ligase, represses LEF1 activity by sequestration into nuclear bodies. Genes Dev 15:3088–3103

75. Chu Y, Yang X (2010) SUMO E3 ligase activity of TRIM proteins. Oncogene 2011 30:9 30:1108–1116

76. Pichler A, Gast A, Seeler JS, Dejean A, Melchior F (2002) The Nucleoporin RanBP2 Has SUMO1 E3 Ligase Activity. Cell 108:109–120

77. Reynolds A, Qiao H, Yang Y, et al (2013) RNF212 is a dosage-sensitive regulator of crossing-over during mammalian meiosis. Nature Genetics 2013 45:3 45:269–278

78. Liang Q, Deng H, Li X, Wu X, Tang Q, Chang T-H, Peng H, Rauscher FJ, Ozato K, Zhu F (2011) Tripartite Motif-Containing Protein 28 Is a Small Ubiquitin-Related Modifier E3 Ligase and Negative Regulator of IFN Regulatory Factor 7. The Journal of Immunology 187:4754–4763

79. Yamashita D, Moriuchi T, Osumi T, Hirose F (2016) Transcription Factor hDREF Is a Novel SUMO E3 Ligase of Mi2α*. Journal of Biological Chemistry 291:11619–11634

80. Karvonen U, Jääskeläinen T, Rytinki M, Kaikkonen S, Palvimo JJ (2008) ZNF451 Is a Novel PML Body- and SUMO-Associated Transcriptional Coregulator. J Mol Biol 382:585–600

81. Warner LE, Mancias P, Butler IJ, McDonald CM, Keppen L, Koob KG, Lupski JR (1998) Mutations in the early growth response 2 (EGR2) gene are associated with hereditary myelinopathies. Nature Genetics 1998 18:4 18:382–384

82. Garcáa-Gutiérrez P, Juárez-Vicente F, Gallardo-Chamizo F, Charnay P, García-Domínguez M (2011) The transcription factor Krox20 is an E3 ligase that sumoylates its Nab coregulators. EMBO Rep 12:1018–1023

83. Nishida T, Yasuda H (2002) PIAS1 and PIASxα Function as SUMO-E3 Ligases toward Androgen Receptor and Repress Androgen Receptor-dependent Transcription. Journal of Biological Chemistry 277:41311–41317

84. Liang Q, Deng H, Li X, Wu X, Tang Q, Chang T-H, Peng H, Rauscher FJ, Ozato K, Zhu F (2011) Tripartite Motif-Containing Protein 28 Is a Small Ubiquitin-Related Modifier E3 Ligase and Negative Regulator of IFN Regulatory Factor 7. The Journal of Immunology 187:4754–4763

85. Zhao Q, Xie Y, Zheng Y, Jiang S, Liu W, Mu W, Liu Z, Zhao Y, Xue Y, Ren J (2014) GPS-SUMO: A tool for the prediction of sumoylation sites and SUMO-interaction motifs. Nucleic Acids Res 42:W325–W330

86. Ren J, Gao X, Jin C, Zhu M, Wang X, Shaw A, Wen L, Yao X, Xue Y (2009) Systematic study of protein sumoylation: Development of a site-specific predictor of SUMOsp 2.0. Proteomics 9:3409–3412

87. Eisenhardt N, Chaugule VK, Koidl S, et al (2015) A new vertebrate SUMO enzyme family reveals insights into SUMO-chain assembly. Nature Structural & Molecular Biology 2015 22:12 22:959–967

88. Lumpkin RJ, Gu H, Zhu Y, Leonard M, Ahmad AS, Clauser KR, Meyer JG, Bennett EJ, Komives EA (2017) Site-specific identification and quantitation of endogenous SUMO modifications under native conditions. Nature Communications 2017 8:1 8:1–11

89. Hendriks IA, Lyon D, Su D, Skotte NH, Daniel JA, Jensen LJ, Nielsen ML (2018) Site-specific characterization of endogenous SUMOylation across species and organs. Nature Communications 2018 9:1 9:1–17

90. Chen S, Francioli LC, Goodrich JK, et al (2023) A genomic mutational constraint map using variation in 76,156 human genomes. Nature 2023 625:7993 625:92–100

91. Kraemer BC, Schuck T, Wheeler JM, Robinson LC, Trojanowski JQ, Lee VMY, Schellenberg GD (2010) Loss of Murine TDP-43 disrupts motor function and plays an essential role in embryogenesis. Acta Neuropathol 119:409–419

92. Khalfallah Y, Kuta R, Grasmuck C, Prat A, Durham HD, Vande Velde C (2018) TDP-43 regulation of stress granule dynamics in neurodegenerative disease-relevant cell types. Sci Rep 8:7551

93. Sidibé H, Khalfallah Y, Xiao S, et al (2021) TDP-43 stabilizes G3BP1 mRNA: relevance to amyotrophic lateral sclerosis/frontotemporal dementia. Brain. 10.1093/brain/awab217

94. Shelkovnikova TA, Kukharsky MS, An H, Dimasi P, Alexeeva S, Shabir O, Heath PR, Buchman VL (2018) Protective paraspeckle hyper-assembly downstream of TDP-43 loss of function in amyotrophic lateral sclerosis. Mol Neurodegener 13:1– 17

95. Wang C, Duan Y, Duan G, et al (2020) Stress Induces Dynamic, Cytotoxicity-Antagonizing TDP-43 Nuclear Bodies via Paraspeckle LncRNA NEAT1-Mediated Liquid-Liquid Phase Separation. Mol Cell 79:443–458.e7

96. Yu H, Lu S, Gasior K, et al (2021) HSP70 chaperones RNA-free TDP-43 into anisotropic intranuclear liquid spherical shells. Science (1979). 10.1126/SCIENCE.ABB4309/SUPPL_FILE/ABB4309_YU_SM.PD F

97. Hipp MS, Kasturi P, Hartl FU (2019) The proteostasis network and its decline in ageing. Nature Reviews Molecular Cell Biology 2019 20:7 20:421–435

98. Geertsma HM, Suk TR, Ricke KM, Horsthuis K, Parmasad JLA, Fisk ZA, Callaghan SM, Rousseaux MWC (2022) Constitutive nuclear accumulation of endogenous alpha-synuclein in mice causes motor impairment and cortical dysfunction, independent of protein aggregation. Hum Mol Genet 31:3613–3628

99. Yu B, Pamphlett R (2017) Environmental insults: Critical triggers for amyotrophic lateral sclerosis. Transl Neurodegener 6:1–10

100. Maurel C, Chami AA, Thépault RA, Marouillat S, Blasco H, Corcia P, Andres CR, Vourc’h P (2020) A role for SUMOylation in the Formation and Cellular Localization of TDP-43 Aggregates in Amyotrophic Lateral Sclerosis. Mol Neurobiol 57:1361–1373

101. Kirola L, Mukherjee A, Mutsuddi M (2022) Recent Updates on the Genetics of Amyotrophic Lateral Sclerosis and Frontotemporal Dementia. Molecular Neurobiology 2022 59:9 59:5673–5694

102. Wang A, Conicella AE, Schmidt HB, et al (2018) A single N-terminal phosphomimic disrupts TDP-43 polymerization, phase separation, and RNA splicing. EMBO J 37:e97452

103. Brady OA, Meng P, Zheng Y, Mao Y, Hu F (2011) Regulation of TDP-43 aggregation by phosphorylation andp62/SQSTM1. J Neurochem 116:248–259

104. Wegorzewska I, Bell S, Cairns NJ, Miller TM, Baloh RH (2009) TDP-43 mutant transgenic mice develop features of ALS and frontotemporal lobar degeneration. Proc Natl Acad Sci U S A 106:18809–18814

105. Campbell KM, Xu Y, Patel C, et al (2021) Loss of TDP-43 in male germ cells causes meiotic failure and impairs fertility in mice. Journal of Biological Chemistry. 10.1016/j.jbc.2021.101231

106. Milstead RA, Link CD, Xu Z, Hoeffer CA (2023) TDP-43 knockdown in mouse model of ALS leads to dsRNA deposition, gliosis, and neurodegeneration in the spinal cord. Cerebral Cortex (New York, NY) 33:5808

107. Yang C, Qiao T, Yu J, et al (2022) Low-level overexpression of wild type TDP-43 causes late-onset, progressive neurodegeneration and paralysis in mice. PLoS One 17:e0255710

108. Buratti E (2018) TDP-43 post-translational modifications in health and disease. Expert Opin Ther Targets. 10.1080/14728222.2018.1439923

109. Chatterjee M, Özdemir S, Fritz C, et al (2024) Plasma extracellular vesicle tau and TDP-43 as diagnostic biomarkers in FTD and ALS. Nature Medicine 2024 30:6 30:1771–1783

110. Feiler MS, Strobel B, Freischmidt A, et al (2015) TDP-43 is intercellularly transmitted across axon terminals. Journal of Cell Biology 211:897–911

111. Iguchi Y, Eid L, Parent M, et al (2016) Exosome secretion is a key pathway for clearance of pathological TDP-43. Brain 139:3187–3201

112. Leibiger C, Deisel J, Aufschnaiter A, Ambros S, Tereshchenko M, Verheijen BM, Büttner S, Braun RJ (2018) TDP-43 controls lysosomal pathways thereby determining its own clearance and cytotoxicity. Hum Mol Genet 27:1593–1607

113. Liu G, Byrd A, Warner AN, Pei F, Basha E, Buchanan A, Buchan JR (2019) Cdc48/VCP and Endocytosis Regulate TDP-43 and FUS Toxicity and Turnover. Mol Cell Biol. 10.1128/mcb.00256-19

114. Liu G, Coyne AN, Pei F, Vaughan S, Chaung M, Zarnescu DC, Buchan JR (2017) Endocytosis regulates TDP-43 toxicity and turnover. Nat Commun 8:2092

115. Scotter EL, Vance C, Nishimura AL, et al (2014) Differential roles of the ubiquitin proteasome system and autophagy in the clearance of soluble and aggregated TDP-43 species. J Cell Sci 127:1263–1278

116. Wang X, Fan H, Ying Z, Li B, Wang H, Wang G (2010) Degradation of TDP-43 and its pathogenic form by autophagy and the ubiquitin-proteasome system. Neurosci Lett 469:112–116

117. Yau TY, Molina O, Courey AJ (2020) SUMOylation in development and neurodegeneration. Development. 10.1242/DEV.175703

118. Princz A, Tavernarakis N (2017) The role of SUMOylation in ageing and senescent decline. Mech Ageing Dev 162:85–90

119. White MA, Kim E, Duffy A, et al (2018) TDP-43 gains function due to perturbed autoregulation in a Tardbp knock-in mouse model of ALS-FTD. Nat Neurosci 21:1138

120. Fratta P, Sivakumar P, Humphrey J, et al (2018) Mice with endogenous TDP-43 mutations exhibit gain of splicing function and characteristics of amyotrophic lateral sclerosis. EMBO J. 10.15252/embj.201798684

121. Abramson J, Adler J, Dunger J, et al (2024) Accurate structure prediction of biomolecular interactions with AlphaFold 3. Nature 2024 630:8016 630:493–500

122. Dereeper A, Guignon V, Blanc G, et al (2008) Phylogeny.fr: robust phylogenetic analysis for the non-specialist. Nucleic Acids Res 36:W465–W469

123. Gertsenstein M, Nutter LMJ (2018) Engineering Point Mutant and Epitope-Tagged Alleles in Mice Using Cas9 RNA-Guided Nuclease. Curr Protoc Mouse Biol 8:28– 53

124. Miedel CJ, Patton JM, Miedel AN, Miedel ES, Levenson JM (2017) Assessment of Spontaneous Alternation, Novel Object Recognition and Limb Clasping in Transgenic Mouse Models of Amyloid-β and Tau Neuropathology. JoVE (Journal of Visualized Experiments) 2017:e55523

125. Guyenet SJ, Furrer SA, Damian VM, Baughan TD, la Spada AR, Garden GA (2010) A Simple Composite Phenotype Scoring System for Evaluating Mouse Models of Cerebellar Ataxia. JoVE (Journal of Visualized Experiments) e1787

126. Lê S, Josse J, Husson F (2008) FactoMineR: An R Package for Multivariate Analysis. J Stat Softw 25:1–18

127. Alboukadel K, Fabian M (2020) Extract and Visualize the Results of Multivariate Data Analyses [R package factoextra version 1.0.7]. https://CRAN.R-project.org/package=factoextra. Accessed 8 Apr 2024

